# A new approach of dissecting genetic effects for complex traits

**DOI:** 10.1101/2020.10.16.336180

**Authors:** Meng Luo, Shiliang Gu

**Author notes:** Correspondence and requests for materials should be addressed to Meng Luo or Shiliang Gu.

## Abstract

During the past decades, genome-wide association studies (GWAS) have been used to successfully identify tens of thousands of genetic variants associated with complex traits included in humans, animals, and plants. All common genome-wide association (GWA) methods rely on population structure correction to avoid false genotype and phenotype associations. However, population structure correction is a stringent penalization, which also impedes the identification of real associations. Here, we used recent statistical advances and proposed iterative screen regression (ISR), which enables simultaneous multiple marker associations and shown to appropriately correction population stratification and cryptic relatedness in GWAS. Results from analyses of simulated suggest that the proposed ISR method performed well in terms of power (sensitivity) versus FDR (False Discovery Rate) and specificity, also less bias (higher accuracy) in effect (PVE) estimation than the existing multi-loci (mixed) model and the single-locus (mixed) model. We also show the practicality of our approach by applying it to rice, outbred mice, and A.thaliana datasets. It identified several new causal loci that other methods did not detect. Our ISR provides an alternative for multi-loci GWAS, and the implementation was computationally efficient, analyzing large datasets practicable (n>100,000).

## Introduction

Genome-wide association studies (GWASs) have been increasingly prominent in detecting genetic variants associated with complex traits and disease, while the identified variants significantly explain only a fraction of total phenotypic variance, resulting in the so-called ‘missing heritability’, but adventitiously pinpointing biological mechanisms^1,2^. Commonly, the individuals used in GWA studies are not related to each other, some degrees of confounding cryptic relatedness and population stratification are inevitable. Simultaneously, there is another confounding existing, which is the genetic background (non-genetic factor), such that the population structure control does not do well in very complex cases^3^. If these happen can lead to spurious associations (there is only correlated with the phenotype and markers, but not substantially associated with causal variants) between the phenotype and unlinked candidate loci (Mendel’s Second Law) ^4,5^, which brought about the challenge problem that how to efficiently conquer test for associations in the presence of population structure (including cryptic relatedness and population stratification) and genetic background.

During the past two decades, there are many solutions to the problem of population structure, including genomic control(GC)^6,7^, structured association (SA)^8–10^, regression control (RC)^11,12^, principal components adjustment (PCA)^13,14^ and mixed regression models(MRM)^15–17^. In the regression control and principal components adjustment approaches, population structure both are taken into account by including covariates in the regression model. In the absence of ascertainment bias, RC performed similarly to GC and SA, while being computationally fast and allowing the flexibility of the regression framework which including backward (stepwise) selection and shrinkage penalty approach^12^. Howbeit, with ascertainment bias, the RC approach substantially outperformed GC^5^. These proposals only perform well in simple cases, however, show poorly when the population structure is more complex^18^.

Incontrovertibly, the current method that linear mixed model (LMM) has extensively used for GWA studies, having been shown to perform well in humans, plants, and animals^19–21^. The linear mixed model that included approximate methods P3D^22^, EMMAx^23^, and GRAMMAR-Gamma^24^, also exact methods EMMA^16^, FaST-LMM^25^, GEMMA^26^, and so forth. It both models the genotype effect as a random term in a linear mixed model, by explicitly involving a similarity matrix (called genomic relationship matrix (GRM)^27^) or covariance structure between the individuals, which it can synchronously correct the population structure and the genetic background. As these mixed-model methods that perform pretty well, but GWAS power remains limited^28^. On the one hand, it both are based on single-locus tests, while the most complex traits controlled by several substantial effects loci and numerous polygenes with minor effects, these univariate linear mixed model approaches may not be adequate, especially in complicated individual relatedness^29^. The inflation of single-locus association test is expected for complex traits, even in the absence of population structure^30^. On the other hand, compared with the traditional linkage mapping, by including multiple cofactors in the genetic model (multiple-loci test) is a prominent alternative and indisputable, which the main feature is the ability to control genomic background effects. Also, a multi-loci test of association has shown outperform single-locus analysis of association^31–33^. However, the main problem in GWAS that the number of subjects, *n*, is in the hundreds or thousands, while *p* could be a range of millions of genetic features. Moreover, the number of loci (gene) exhibiting a detectable association with a trait is minimal. It is a fundamental problem in high-dimensional variable selection. Several methods have been developed to address these issues, such as LASSO^34,35^, stepwise regression^36–38^, penalized logistic regression^39^ and penalized multiple regression ^40^.

For the past decades, based on these methods, where several new multi-locus methodologies have been developed. For example, MLMM^33^, where stepwise mixed-model regression with forwarding inclusion and backward elimination, showed the advantage of computationally efficient and outperform the univariate mixed model for GWAS. LMM-Lasso^41^, where combines the benefits of established linear mixed models (LMM) with sparse Lasso regression. Some of the others, BSLMM^42^, MRMLM^43^, and FASTmrEMMA^44^, both are based on the mixed model. Recently, FarmCPU^28^ and QTCAT^45^ are not based on a mixed model. However, FarmCPU by iterating usage of fixed and random effect models, which improved the power and computation time both than the univariate and multivariate mixed model. QTCAT combining those highly correlated markers, which cannot be distinguished for their contribution to the phenotype and enabling simultaneous correction of the population structure and also reflects the polygenic nature of complex traits better than single-marker methods and outperform traditional linkage mapping.

Whereas hypothesis tested, have been changed by the use of a genomic relationship matrix as the random effect to correct for population structure and infinitesimal genetic background. Where we focus on test multiple loci to effects the phenotype that is neither explained by population structure nor by the genetic background^45^. It is problematic that the trait model assumptions to corroborate in reality, which ultimately leads to failures in identifying causal loci^29,45–47^. Here we introduce a new unique variable selection procedure of regression statistic method, call Iterative screen regression. We formulated a new regression information criterion (RIC) and used this criterion as the objective function of the entire variable screen process. We evaluate various model selection criteria through simulations, which suggest that the proposed ISR method performs well in terms of FDR and power. Finally, we show the usefulness of our approach by applying it to *A. thaliana* and mouse data.

## Results

### Method overview

An overview of our method is provided in the Methods section, with details provided in the Supplementary Note. Briefly, we offered a new regression statistics method and combined a unique variable screening procedure (Fig.1).

**Fig. 1.**
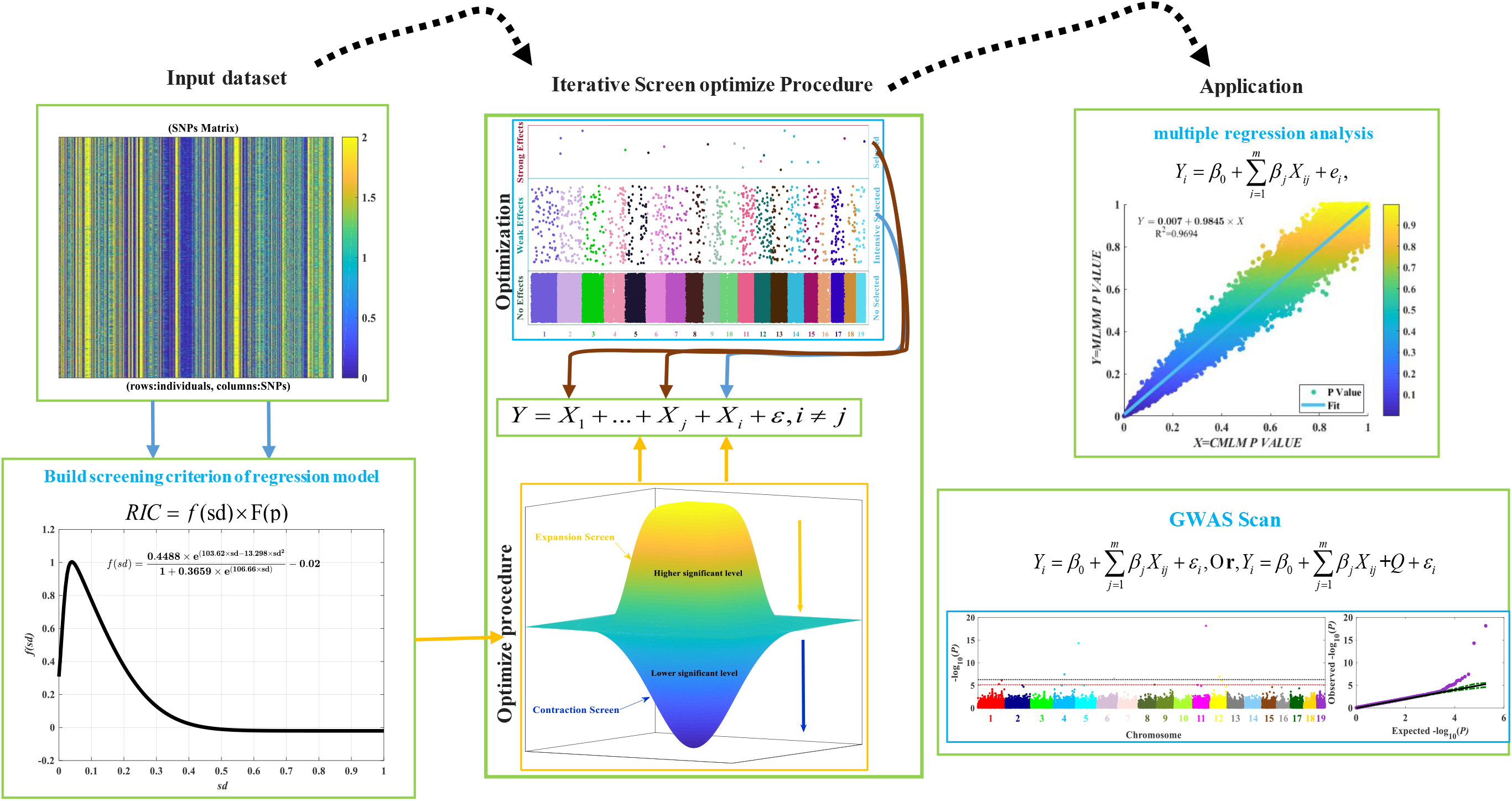
Schematic overviewof model-basedis repeatedly screening stepwiseregressionfor GWAS. The first input dataset with markers (SNPs) matrixrepresentingindividual genotypes (rows) of a population with alleles (0, 2, and 1, missing genotypes will be replaced by the mean genotype or imputed by others complicate algorithm) per marker (columns). Secondly, we formulated a regression information criterion (RIC, objective function) as the screening criterion of the regression model. Combined, the repeatedly proposed screen optimizes the procedure, which mainly included expansion screen and contraction select two -steps (Supplementary Note). The third, apply it to multiple regression analysis and genome-wide association study scan.

### Simulations

We first compare the performance of ISR with several other commonly used association mapping methods using simulations. A total of six different methods are included for comparison: (1) CMLM^48^; (2) LMM (GEMMA) and LM^26^; (3) MLMM^33^; (4) FarmCPU^28^; (5) FASTmrEMMA^44^; (6) FaSTLMM^49^; (7) PLINK (Fisher’s exact test)^50^.

To make our simulations as real as possible, we used genotypes from an existing two model species (*A. thaliana* and mouse), the previous dataset had been widely used to as simulating data including all the above comparison methods. GWAS dataset was simulated by adding phenotypic effects to real genotypic data from *A. thaliana* data under two different scenarios (I, II) ( Methods section for details): a 10-locus model and a 100-locus model. These scenarios have already been simulated in previously^33,44^. In scenarios I, the power for each causal SNPs was defined as the proportion of samples where the causal SNPs were detected (the P-value is smaller than the designated threshold. See the methods all character, Supplementary Table 1). Where we can see that with different heritabilities by each casual loci SNPs, such that multi-locus model including ISR, FASTmrEMMA, FarmCPU outperformed than the mixed model including single-locus(LMM, CMLM) and multi-locus(MLMM); moreover, ISR detected the small effect by the casual loci own more power than the others methods, especially the mixed model (GEMMA, MLMM, CMLM) (Fig.2a), as the following simulation also showed the same phenomena. According to this, all methods’ precision—here defined as the percentage of true positives of all reported loci, where ISR at a level of 5% Bonferroni correction outperformed than the others methods was 92.41%, 80.77%, 78.48%, 68.20%, 65.92% and 65.58% (Fig.2b), respectively. Although the FASTmrEMMA detected the most casual loci, the true positive only almost equal ISR methods detected, while the FDR bigger than ISR nearly 2.8 times. In a word, ISR performs high power and low FDR in the sample trait than other methods. For the sophisticated trait included, 100 locus model is shown, at a different level of heritability 0.25 (low), 0.5 (middle), and 0.75 (high), which can be summarized as follows. First, methods that use a kinship term to correct population structure outperform comparable methods that do not (FASTmrEMMA, FarmCPU, GEMMA, MLMM, CMLM versus LM, respectively). Second, the mixed model, including the single-locus and multi-locus model performed almost equivalent. Third, in low-level heritability, ISR comparable the mixed model (FASTmrEMMA, GEMMA, MLMM, CMLM) and outperformed than FarmCPU and LM. While in the middle and high-level heritability, the performance of ISR more than FarmCPU and other methods (Fig.3 and Supplementary Figs.1-2).

**Fig. 2.**
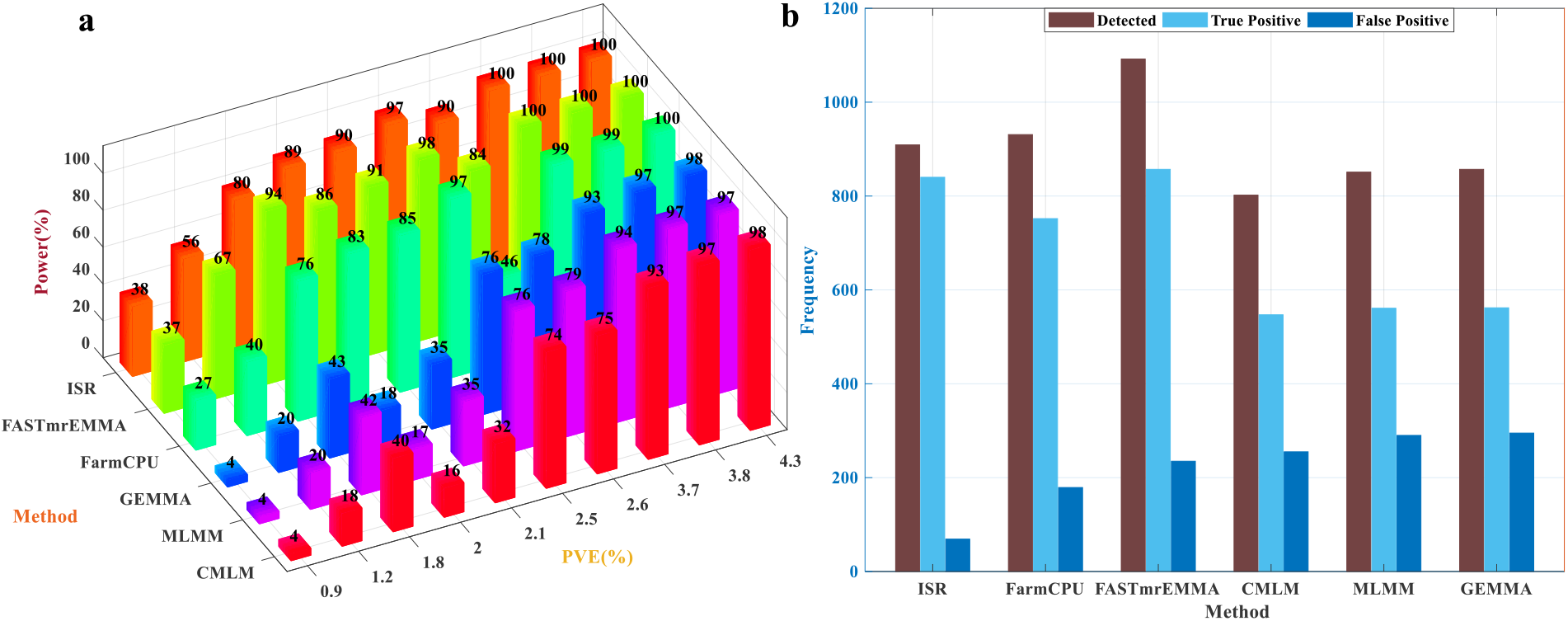
Comparison of ISR with the single-locus and multi-locus approaches. **(a)** The detected power in a different proportionof phenotypicvariation explained (PVE) by genotyped SNPs (10 casual loci) and without considered the window size (means, the 0kb window size) and 100 replicates. **(b)** Compared the number of detected, true positive and false positive, also the FDR in the different genetic models.

**Fig.3.**
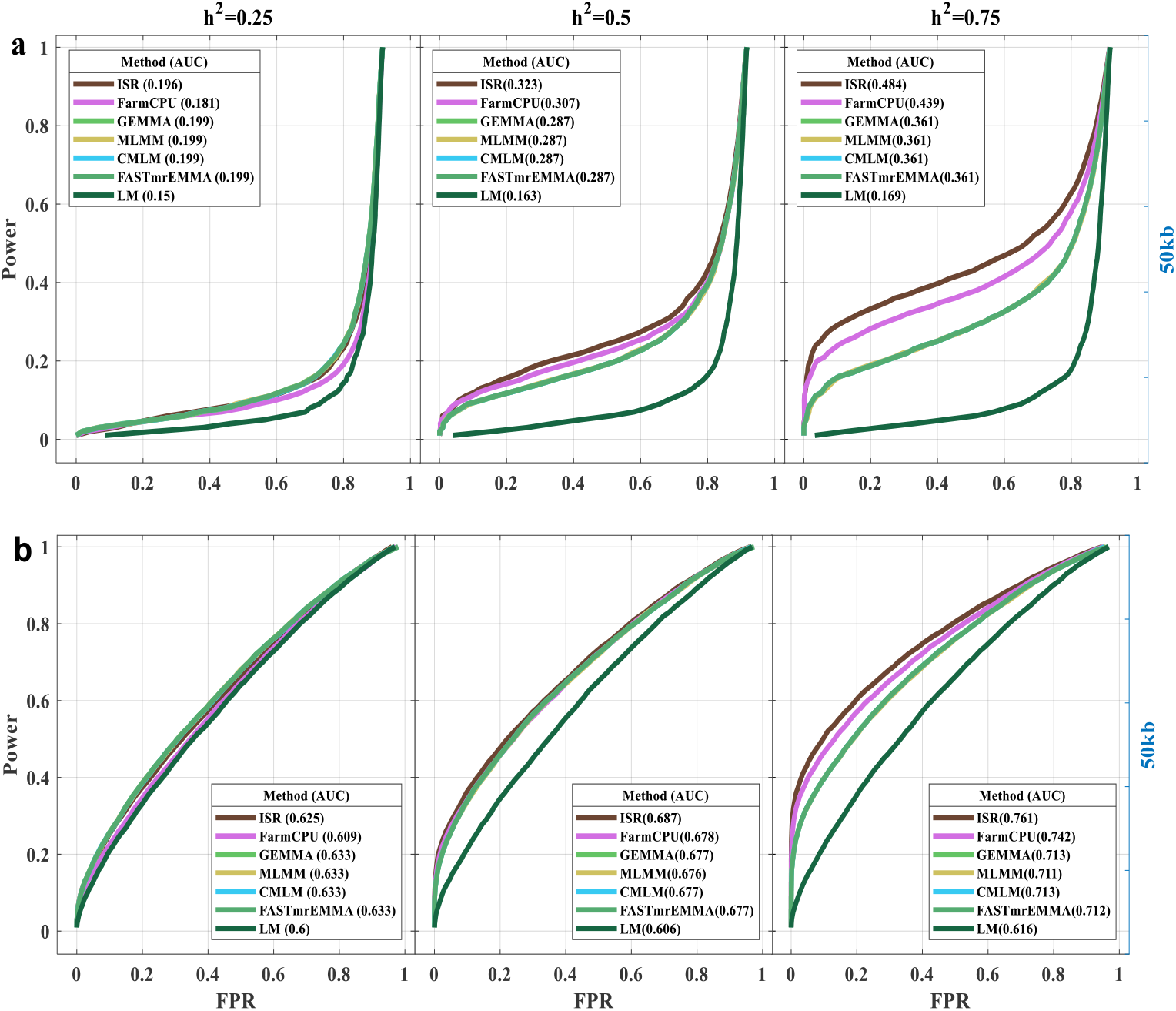
Performances of TPR (Power) versus FDR and FPR in Arabidopsis dataset. A receiver operating characteristic curve for seven methods were performed to test Power/FDR (a) and Power/FPR (b) in the second simulation additive genetic effects controlled by 100 causal loci with three phenotypic heritabilities 0.25(left), 0.5(middle) and 0.75(right), including ISR, FarmCPU, GEMMA, MLMM, CMLM, FASTmrEMMA, and LM methods. The casuallociwere randomly sampled fromall the SNPs in each dataset. Power was examined under different levels of FDR and FPR. A causal SNP was considered to be detected if an SNP within 50 kb on either sidewas determined to have a significant association (results for other window sizes are given in Supplementary Figs.1-2), otherwise, is considered a false positive. The performance of detecting associations is measured by the area under the curve (AUC), where a higher value indicates better performance.

The first CFW mice dataset simulation (scenarios III). Where the phenotype variation controlled by 50-locus is shown (Supplementary Figs.3-5), in which at different levels of heritability 0.25 (low), 0.5 (middle), and 0.75 (high) settings can be summarized as follows. On the one hand, ISR performed well regarding power versus FDR and FPR than other methods. On the other hand, the multi-locus (mixed) model outperformed than single-locus (mixed) model (ISR, FarmCPU, MLMM, FASTmrEMMA versus GEMMA, CMLM, LM). Moreover, with the increase of heritability (0.25~0.75), ISR performs well that get lower FDR, while other methods almost unchanged (Supplementary Fig.6). The second simulation (scenarios IV), another controlled by the 100-locus model, is shown (Fig.4) but only set a level of heritability was 0.5. The performance of the used full dataset is the same as random sample data from all genome. On the one hand, ISR also performed well regarding power versus FDR and FPR than other methods. On the other hand, the multi-locus (mixed) model outperformed than single-locus (mixed) model (ISR, FarmCPU, MLMM, FASTmrEMMA versus GEMMA, CMLM, LM). It is indicated that randomly choose the SNPs from all cover genome (scenarios I-III) or using all genome datasets for simulation, and both results were identically ^33^.

**Fig.4.**
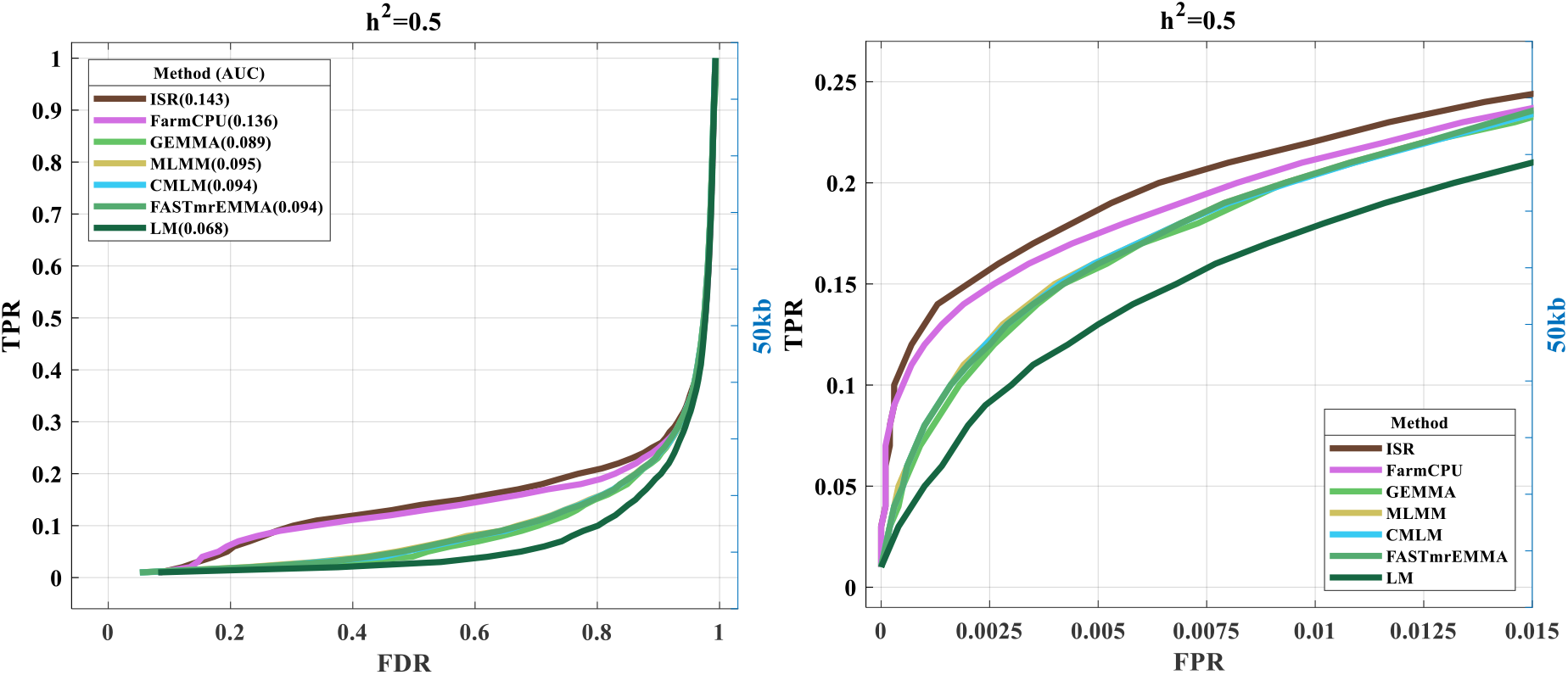
Performances of TPR(Power) versus FDRand FPR in full CFW mice genome dataset. The fourth simulation additive genetic effects are controlled by 100 causal loci with a phenotypic heritability 0.5. Here, a causal SNP was considered to be detected if an SNP within 50 kb on either side was determined to have a significant association, otherwise, is considered a false positive. The Area Under the Curves (AUC) is also displayed separately for TPR (power) versus FDR. The performance of detecting associations is measured by the area under the curve (AUC), where a higher value indicates better performance.

The last two simulations (scenarios V(1-2)) using a human dataset derived from PLINK2^50^ (details seeing the Methods section). Compared to the power, ISR had a significantly larger AUC than FarmCPU, FaSTLMM, and PLINK-Fisher for both TPR versus FDR and TPR versus FPR in both simulations (Fig.5b). PLINK-Fisher had a smaller AUC than ISR, FarmCPU, and FaSTLMM for both comparisons. Especially, FarmCPU only had a significantly larger AUC than FaSTLMM and PLINK-Fisher for TPR versus FDR, not TPR versus FPR in the first simulation (Fig.5b). In other words, FarmCPU had a similar AUC with FaSTLMM and PLINK-Fisher for controlling FPR (Type I error). On the other hand, except PLINK-Fisher that other methods detected power higher along with the samples 10 times increased(1000~10000, Fig.5(a,b)). This situation held true as above two model species (Arabidopsis and mouse).

**Fig.5.**
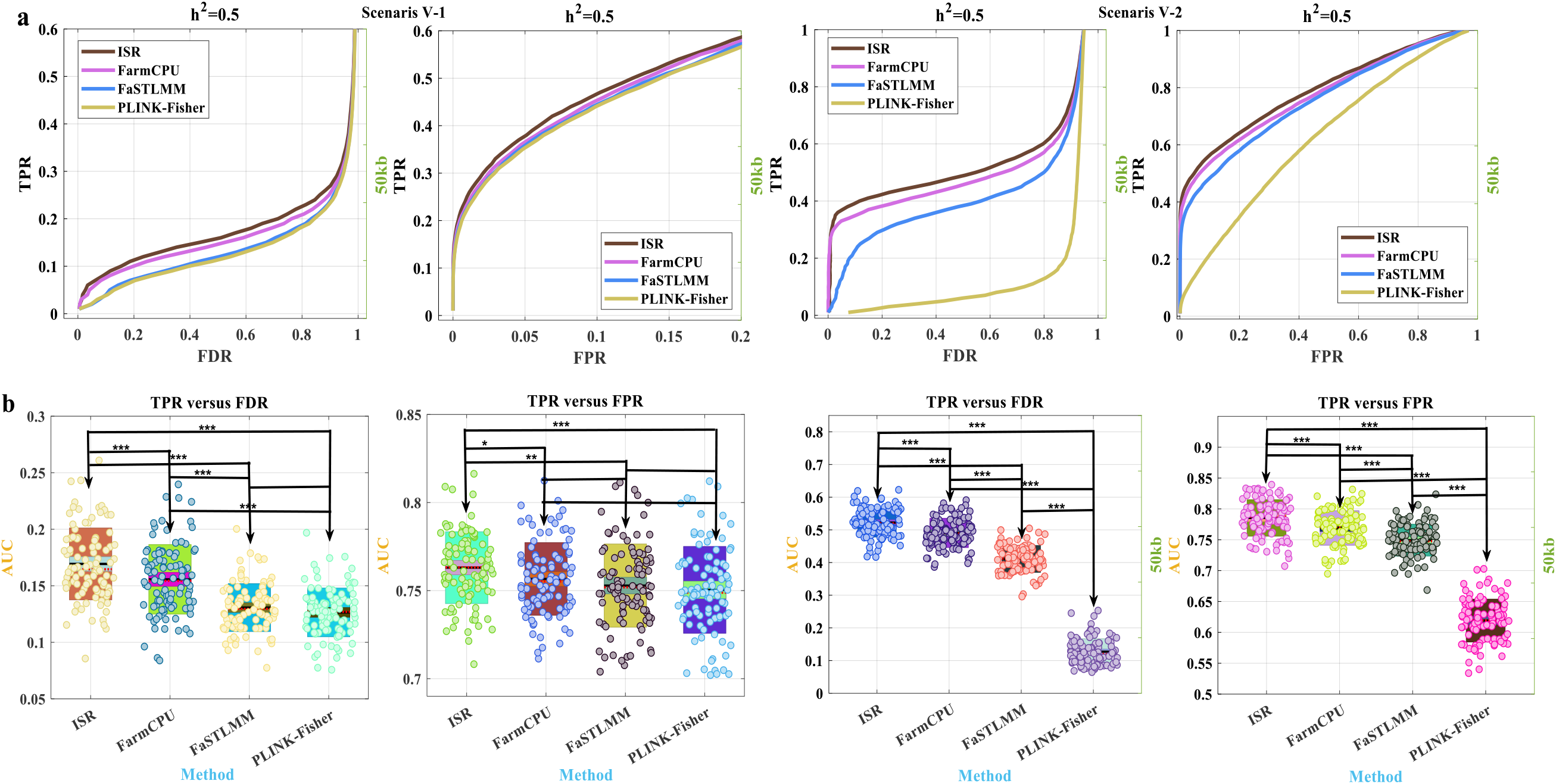
Statistical power and area under the curve to detect causal loci in the fifth simulation scenarios. Statistical power was defined as the proportion of simulated markers detected at cost defined by either False Discovery Rate (FDR) or False Positive Rate (FPR, Type I error).( a) The two types of ReceiverOperating Characteristic (ROC) curves are displayed separately for TPR (true positive rate, power) versus FDR and FPR (the two simulations of Scenarios V (1 −2)). (b) The Area Under the Curves (AUC) are also displayed separately for TPR (true positive rate, power) versus FDR and FPR for 100 simu lations. Four GWAS methods (ISR, FarmCPU, FaSTLMM, and PLINK-Fisher) were compared with phenotypes simulated from real genotypes in humans. The simulated phenotypes had a heritability of 50%, controlled by 100 SNPs. These markers were randomly sampled from the available 100000 (88025) Single Nucleotide Polymorphism (SNPs). (b) .To specify the multiple comparison procedures using Least Significant Difference (LSD) after ANOVA. Here, ‘*’ represents a significant level of 0.05; ‘**’ represents a significant level of 0.01; ‘***’ represents a significant level of 0.001.

### Estimated Effect (PVE)

Generally, if there are environmental factors that influence the phenotype and are correlated with genotype (e.g., due to population structure), then these would undoubtedly affect estimates of SNPs effect, and consequently also affect estimates of other quantities, including the PVE (the total proportion of variance in phenotype explained, or SNPs heritability)^42,51^. So, except comparing the detected power, the accuracy of estimated effect (or PVE) also is one of the keys to whether the model performs well or not. Here, we used the root of mean square error (RMSE) as the accuracy of the PVE estimates obtained by each methods^42^.

In the Arabidopsis simulation dataset (scenarios I). The first simulation set (sparse genetic architecture, which assumes effects are sparse, fixed ten casual SNPs) result showed that ISR, GEMMA, and CMLM significantly perform more stable and accurate (lower RMSE) than other methods (Fig.7), which another two methods (FarmCPU, and FASTmrEMMA) presenting downward bias of PVE estimate. Summarizes the resulting of PVE estimates with six methods. Apparently, except FarmCPU, multi-loci (mixed) model estimated more accuracy than the single-locus mixed model (ISR, FASTmrEMMA versus GEMMA, CMLM). Where the single-locus mixed model is presenting upward bias and tends to decrease along with the PVE (heritability) increased, whereas compared with FarmCPU that tends to downward bias. Furthermore, ISR and FASTmrEMMA accuracy tend to lower along with the increase of heritability (Figs. 7 and Supplementary Fig.7). Where the multiple-loci (mixed) model (ISR and FASTmrEMMA) with lower RMSE estimates of PVE presenting stable and only in the small PVE (low heritability), which tend to less downward bias, on the other hand, detected large effect loci by all methods equally well, while ISR and FarmCPU could expand its findings to loci with smaller effect sizes. Moreover, ISR is more efficient in finding small effect loci along with the increase of heritability (Figs. 6, 7, 8, 9). Human dataset simulation showed the same results, in which ISR had the lowest RMSE than others did three methods (Fig.9).

**Fig.6.**
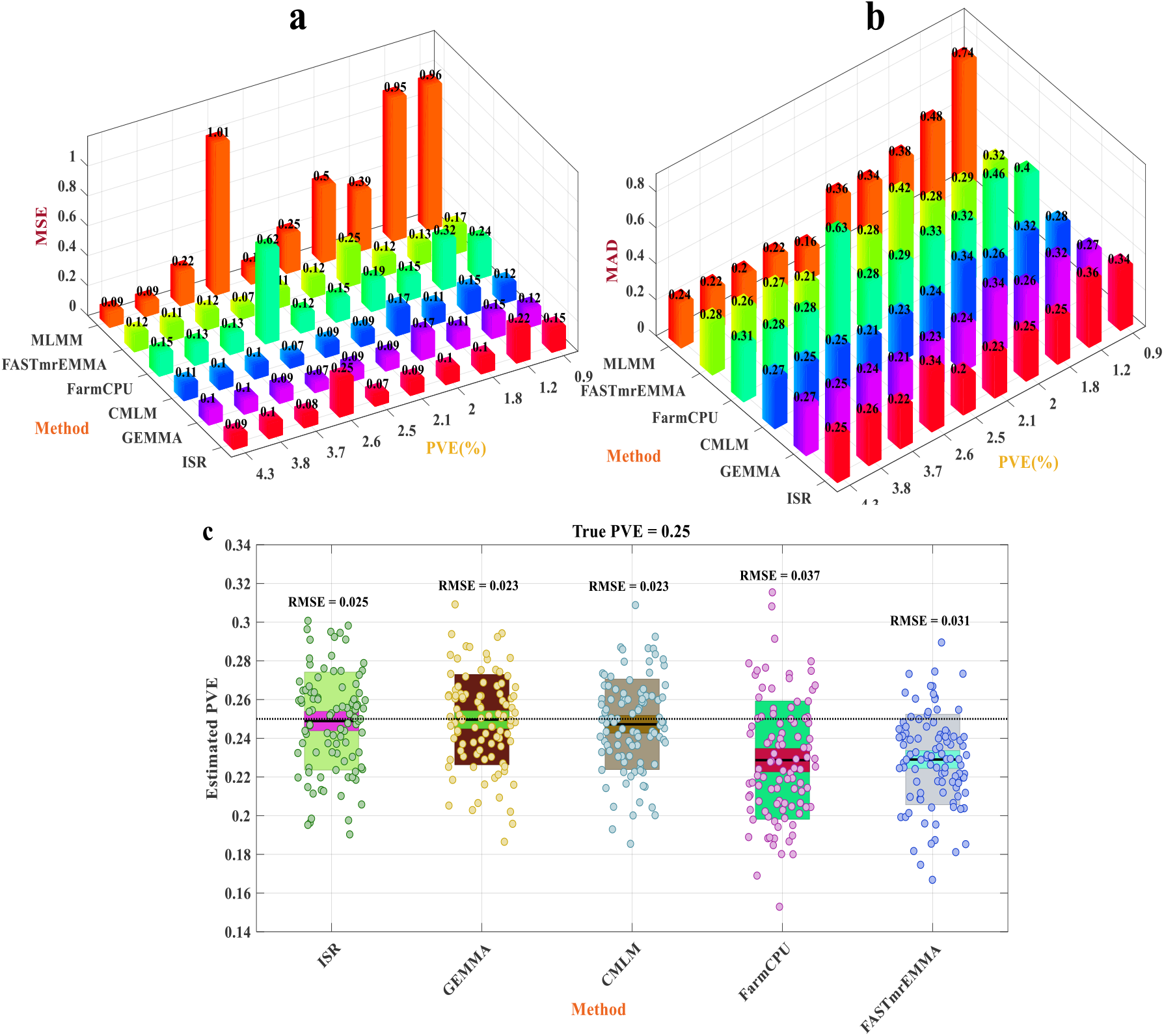
Comparison of accuracy for estimated SNPs effect (and PVE) ISR with other six methods. To measure thebiasof fixed ten casualSNPs effect estimate, where MSE(**a**) and MAD(**b**) were used to compare that in ten different PVE (%). A method with a small MSE (or MAD) is preferable to a method with a large MSE (or MAD)^44^. **(c)** as described^71^, which boxplot showed the small middle patch with a 95% confidence interval (a range of values youcan be 95% confident contains the true mean) for the mean (solid middle line), and the large patch was the SD (standard deviation, where the average difference between the data pointsand their mean). The data points with 100 replicates. Performance of estimating PVE is measured by the root of mean square error (RMSE), where a lower value indicates better performance. The true PVEs are shown as the horizontal dash lines.

**Fig.7.**
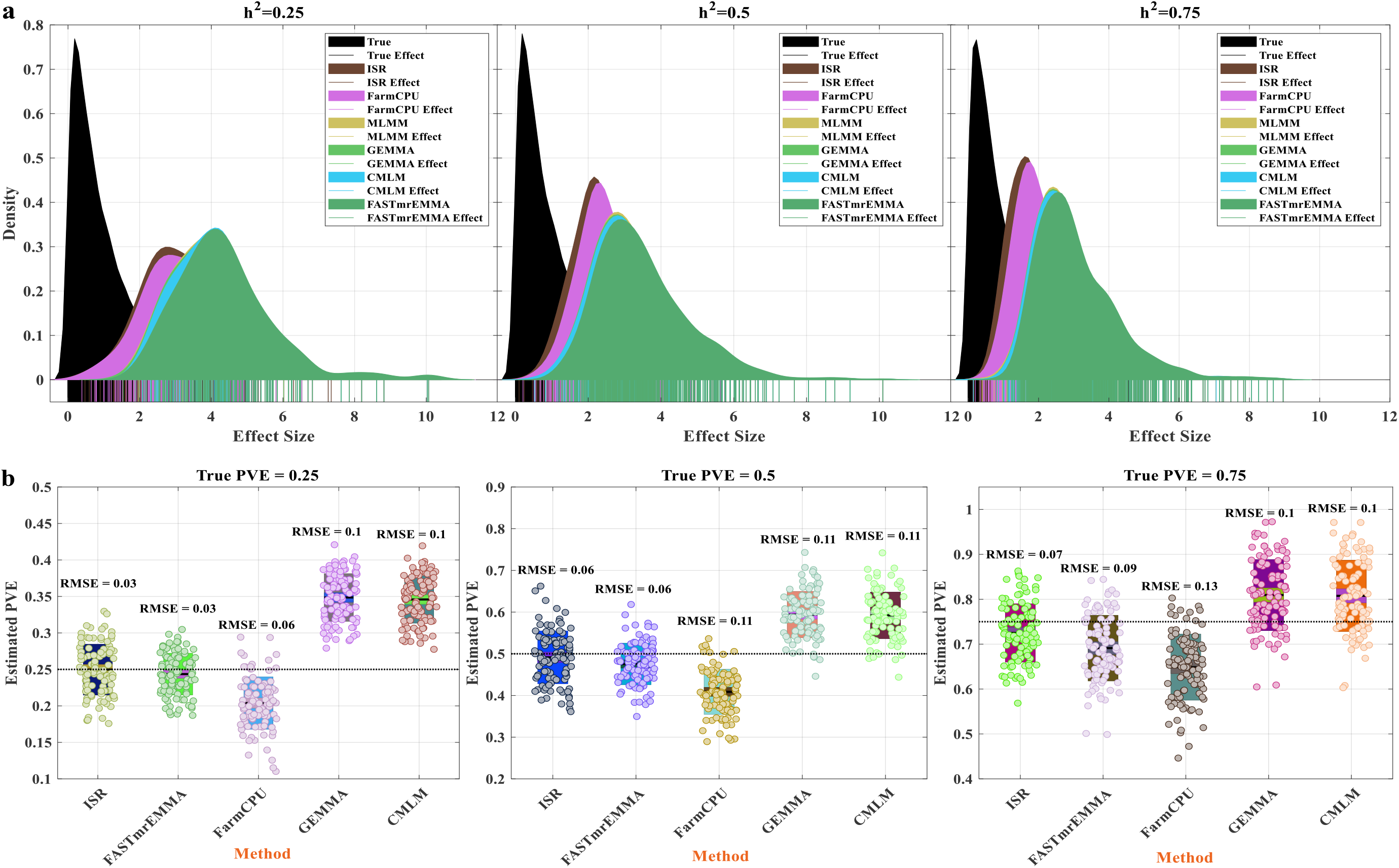
Comparison of detected effect and PVE estimates from five methods in the second simulation scenarios. The distribution of all simulated effects (all true effect) and the distribution of effects of loci identified (100 casual loci within 100 simulations, and only true positive) by six methods. The solid line shows the effect size by different methods. (a) The phenotype with 25%, 50%, and 75% of PVE from left to right, respectively; (b)The bottom boxplot has explained the variance of the loci effect estimated by ISR, FASTmrEMMA, FarmCPU, GEMMA, and CMLM within the 100 simulations. Performance of estimating PVE is measured by the root of mean square error (RMSE), where a lower value indicates better performance. The true PVEs are shown as the horizontal dash lines.

**Fig.8.**
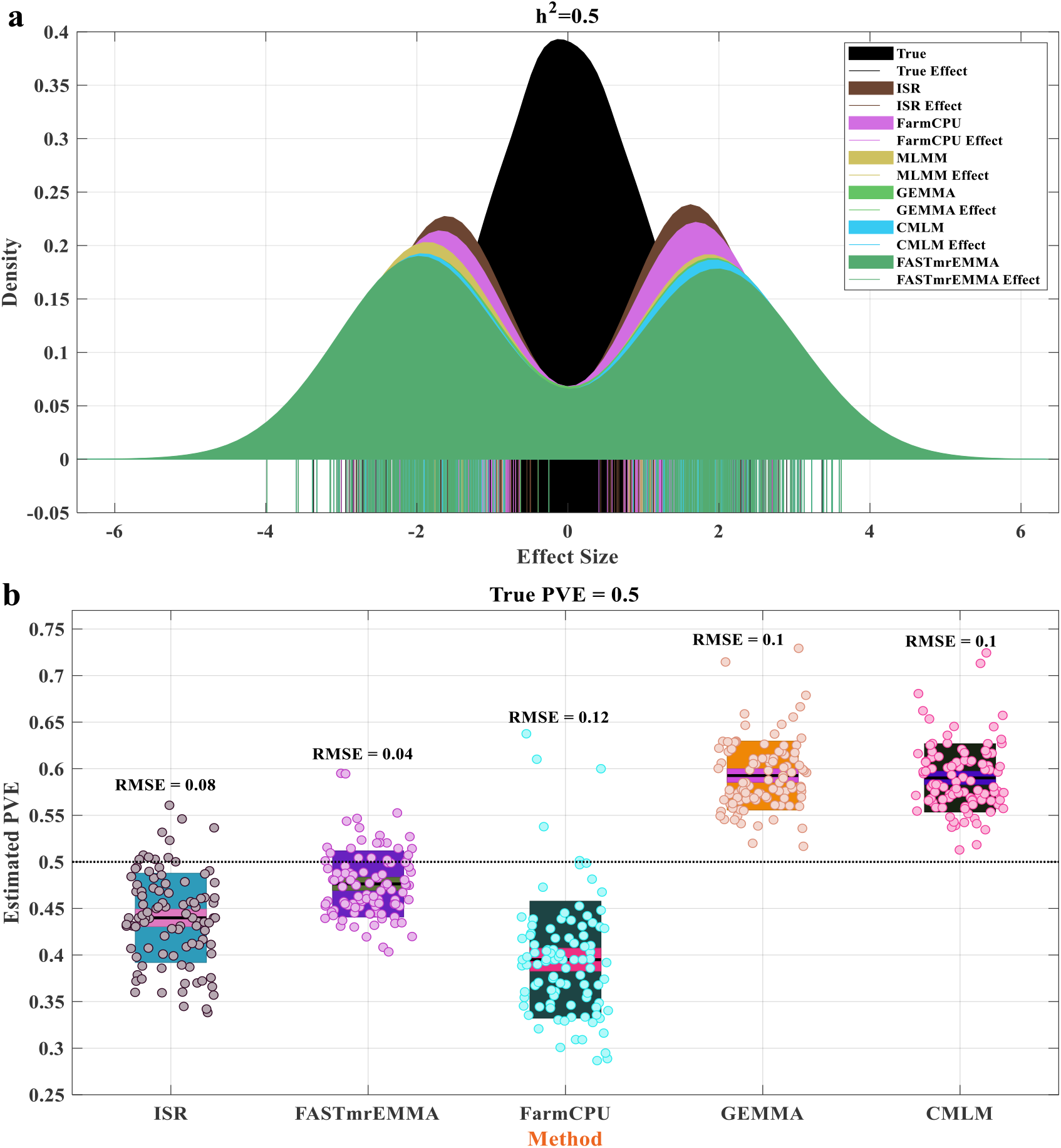
Comparison of detected effect and PVE estimates from five methods in the fourth simulation scenarios. **(a)** The distribution of all simulated effects (all true effect) and the distribution of effects of loci identified (100 casual loci within 100 simulations, and only true positive) by six methods. **(b)** The solid line shows the effect size by different methods and the phenotype with 50% of PVE. The bottom boxplot has explained the variance of the loci effect estimated by ISR, FASTmrEMMA, FarmCPU, GEMMA, and CMLM within the 100 simulations. Performance of estimating PVE is measured by the root of mean square error (RMSE), where a lower value indicates better performance. The true PVEs are shown as the horizontal dash lines.

**Fig.9.**
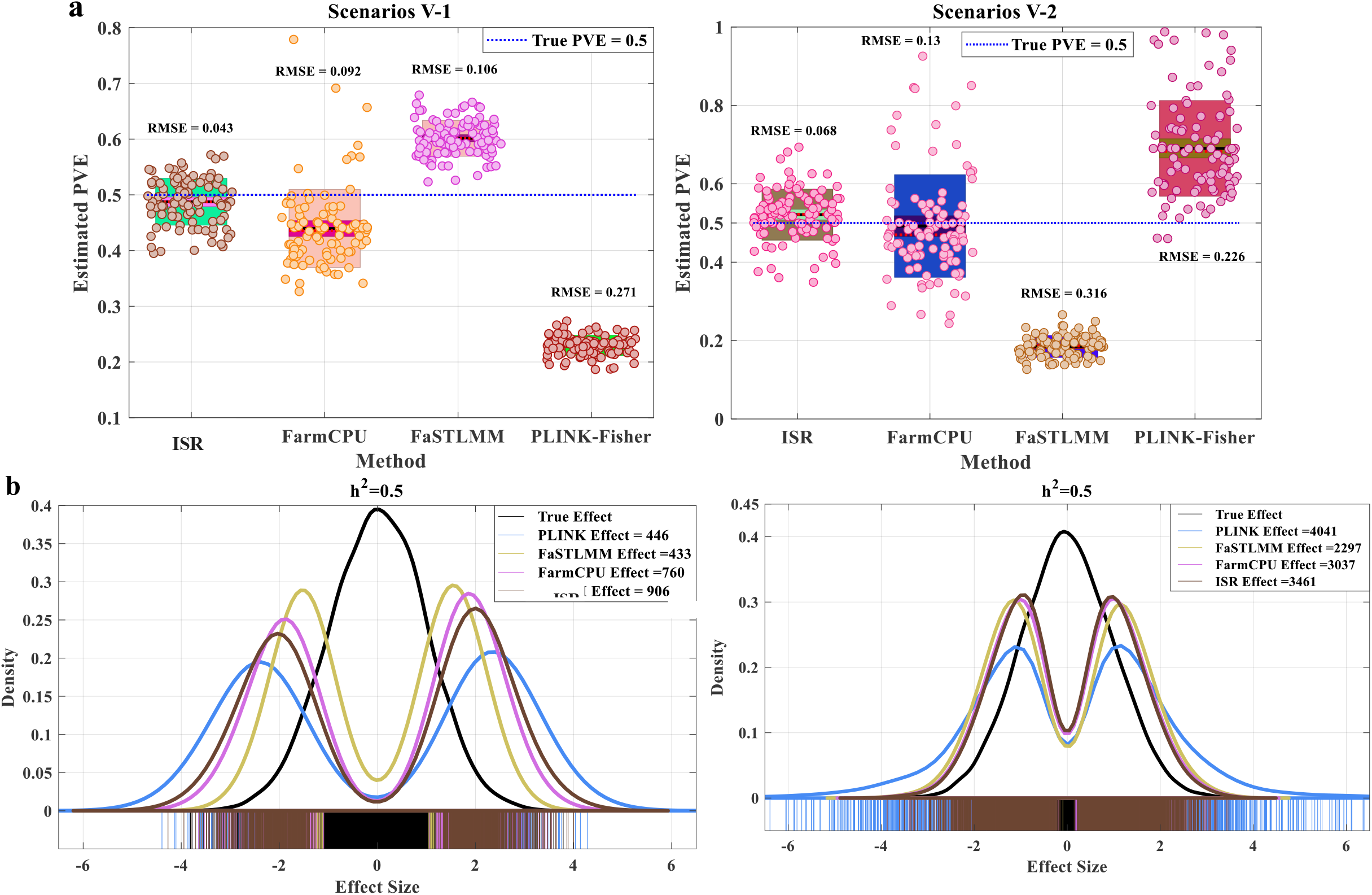
Analysis of the results of GWAS simulations using human dataset. **a** The explained variance of the casual loci effects estimated by ISR, FarmCPU, FaSTLMM, and PLINK-Fisher within the 100 simulations (The two simulations of Scenarios V (1-2)). **b** The distribution of all simulated effects (True Effect, black line) and the distribution of effects of loci identified (after 0.05 Bonferroni correction) by ISR (906 loci and 3461 loci), FarmCPU (760 loci and 3037 loci), FaSTLMM (433 loci and 2297 loci) and PLINK-Fisher (446 loci and 4041), respectively (The two simulations of Scenarios V (1-2).

### Applying ISR to real datasets

To validate and gain further insight ISR, it’s along with FarmCPU, GEMMA, CMLM, MLMM and FASTmrEMMA was used to reanalyze the *A. thaliana* dataset^52^ for all phenotypes related to flowering time and others (Defense-related, Ionomics and Developmental phenotypes, only chosen one). We excluded phenotypes measured for less than 160 accessions to avoid possible ‘small sample size effects, resulting in 13 flowering times phenotypes that were considered. The relatedness between individuals ranges in a wide spectrum leading to a complex population structure^53^. The SNPs identified that using six methods is listed in (Supplementary Table 3). The dataset is characterized by high heritabilities (0.89~1.00), except for the At1CFU2 trait with relatively low heritability (0.54). Moreover, both were small sample sizes (147~194).

Having shown the accuracy of ISR more than other methods in recovering causal SNPs in simulation, we now demonstrate that the ISR better models the genotype-to-phenotype map in Arabidopsis thaliana. ISR methods detected the most SNPs significantly associated close to or in known candidate genes with the above sixteen traits and significantly more than other methods (see Supplementary Fig.6 and Table 1). Such as, based on the SNPs detected by ISR, 13/13 genes were previously reported to be associated with the LN10 (leaf number at flowering time,10℃) trait, while 5/5, 0/0, 0/0, 3/3, and 5/9 genes detected by FarmCPU, GEMMA, CMLM, MLMM, and FASTmrEMMA, respectively ^54,55^. The same as the other traits in Table 1. As corresponding simulation result showed that ISR has higher detected power and lower false discovery rate than other methods in different heritabilities, especially, in high heritability. MLMM result indicated that at the EBIC and mBonf two different model selection criteria, which shown both detected the same genes.

**Table 1.** Comparison of six different methods the associations close to known candidate genes in Arabidopsis thaliana data

| Phenotype | ISR | FarmCPU | GEMMA | CMLM | MLMM(EBIC&mBonf) | FASTmrEMMA |
| --- | --- | --- | --- | --- | --- | --- |
| LD | 13/20 | 6/9 | 9/11 | 1/1 | 0/0 | 5/6 |
| LDV | 9/18 | 5/5 | 3/5 | 0/1 | 0/0 | 6/10 |
| SDV | 15/22 | 4/7 | 3/6 | 0/1 | 0/0 | 2/6 |
| SD | 15/21 | 6/7 | 1/1 | 0/0 | 0/0 | 1/3 |
| FLC | 16/23 | 0/2 | 1/3 | 0/0 | 0/0 | 3/5 |
| FRI | 9/15 | 1/3 | 2/9 | 1/4 | 0/1 | 5/8 |
| FT10 | 15/21 | 4/9 | 4/5 | 0/0 | 0/2 | 1/4 |
| FT16 | 7/14 | 1/2 | 1/2 | 1/1 | 1/1 | 4/8 |
| FT22 | 13/22 | 6/8 | 3/3 | 0/0 | 0/0 | 2/6 |
| FTGH | 12/21 | 2/6 | 13/17 | 0/0 | 0/0 | 2/3 |
| LN10 | 13/13 | 5/5 | 0/0 | 0/0 | 3/3 | 5/9 |
| LN16 | 14/22 | 5/7 | 2/2 | 0/0 | 2/2 | 6/10 |
| LN22 | 16/22 | 6/8 | 0/0 | 1/1 | 0/0 | 8/12 |
| 8WGHLN | 7/14 | 3/3 | 0/0 | 0/0 | 2/2 | 4/9 |
| At1CFU2 | 14/17 | 0/0 | 0/0 | 0/0 | 1/1 | 8/12 |
| RPGH | 12/19 | 0/0 | 0/0 | 0/0 | 0/0 | 7/12 |

The table lists the number of true positives/positives (TP/P) detected (passing the genome-wide significance threshold via Bonferroni correction) by six different methods for all phenotypes related to flowering time in Arabidopsis thaliana and others (Defense-related, Ionomics, and Developmental phenotypes). Pare all causal SNPs, and TP is all causal SNPs that are known candidate genes. All reported candidate genes and the reference literature could be sought on the website (https://www.arabidopsis.org/index.jsp). For each trait, we colored the best method with red and the second-best method with blue.

ISR outperformed other methods concerning controlling inflation of P values, identifying new associated markers, and covering with known loci. We take three phenotypes association test results in *A.thaliana* dataset as an example. The first is bacterial disease resistance (At1 CFU2) with low heritability ^52^, the second is leaf Na^+^ accumulation^56^, and the third is a cellular trait of meristem zone length^57^ both from worldwide *A. thaliana* accessions. The two latter phenotypes have already been reanalyzed by MLMM and QTCAT two methods, respectively.^45,58^. For the At1 CFU2 trait, FarmCPU, CMLM, and FASTmrEMMA both slightly under expected (Supplementary Fig.9), and except FASTmrEMMA identified no associated SNPs above the threshold of 5% after Bonferroni multiple test correction. MLMM and GEMMA controlled inflation well, while only MLMM identified one associated SNP above a threshold of 5% after Bonferroni multiple test correction. Furthermore, ISR not only controlled inflation well but also identified seventeen associated SNPs above the significance threshold, and only three loci out of the known candidate gene (Table 1, Supplementary Fig.8).

Sodium accumulation in the leaves of *A.thaliana* that had detected strongly associated with genotype and expression levels of the Na^+^ transporter *AtHKT1;1* ^56^. Extraordinarily, an SNP located in the first exon of the gene (chromosome4: 6,392,280) shows a highly significant association using an approximate mixed model. We reanalyzed the dataset used six different linear models (as above described). Both methods result indicated that identified the same most significant locus (chromosome4: 6,392,280), while in our study that detected more than the original research, except CMLM. The approximate mixed model CMLM showed the same result with^56^. Three methods CMLM, GEMMA, and FASTmrEMMA show slight inflation, while ISR, MLMM, and FarmCPU controlled inflation well. The two methods ISR and FarmCPU detected one same locus (chromosome2: 5,169,035), while ISR detected four loci in chromosome three which as same as MLMM identified three loci (total three loci) and two loci by FarmCPU (total two loci). Moreover, both one identified by CMLM (total one loci) and GEMMA (total four loci). ISR detected five loci in chromosome four which as same as MLMM identified four loci (total five loci) and also FarmCPU four loci (total five loci); ISR detected two loci in chromosome five only as same as MLMM detected one (total one loci), while between the others methods didn’t identify the same locus (Supplementary Fig.10 and Supplementary Table 4). In others words, except for as same as others methods detected genes, where it indicated that our model always detected more genes (Table 1).

In a recent GWA study^57^, authors using a worldwide collection of 201 natural Arabidopsis accessions to study the genetic architecture of root development. They also use the approximate mixed model and detected only one most significant association (at position 22244990 on chromosome one, an F-box gene). Natural genetic variation influences the meristem zone lengths in roots. Here, as above, our reanalyzed result showed that four methods included CMLM, FarmCPU, GEMMA, and MLMM control inflation well, while no identified association SNPs after 0.05 Bonferroni correction. The FASTmrEMMA showed under deflation, but the final result detected nine SNPs (Supplementary Fig.11 and Supplementary Table 4). While ISR not only controls inflation well but also identified fifteen SNPs also included the position 22244990 on chromosome one and all loci except one both in the candidate gene (Supplementary Fig.11 and Supplementary Table 4). Otherwise, Two methods ISR and FASTmrEMMA detected the same most significant association locus in chromosome three (Supplementary Fig.12).

Carworth Farms White (CFW) mice are a commercially available outbred mouse population. The dataset was previously reanalyzed to show the usefulness of the mixed model^59^. Here, we also reanalyzed the dataset used six different linear models that included a single locus linear model (CMLM and GEMMA) and multiple loci linear models (ISR, MLMM, FarmCPU, and FASTmrEMMA). Compared with the results that SNPs identified by six methods all were listed in (Supplementary Table.5). We also calculated a significance threshold via permutation, which is a standard approach for QTL mapping in mice that controls the type I error rate well (Supplementary Fig.13). We mapped QTLs for ten behavioral and physiological phenotypes, and mapping association results indicated that SNPs detected on different chromosomes by the single locus mixed model (GEMMA and CMLM) and associated by multiple loci linear model (ISR), while except the MLMM, FarmCPU, and FASTmrEMMA methods. Moreover, where the ISR always detected additional significate association locus. The results are mostly consistent with the simulations investigated. For example, QTL mapping for abnormal BMD phenotype that single-locus mixed model (GEMMA and CMLM) identifies two sharp peaks of significantly associated SNPs on chromosome five and eleven, and the most significant associated two loci were rs27024162 and rs32012436 (Supplementary Fig.14). Except for FarmCPU and FASTmrEMMA, those loci are both detected by multiple loci linear (mixed) model (ISR and MLMM) methods. Moreover, in contrast to that ISR, the visualization of Manhattan and QQ (Q stand for Quantile) plot showed that the ISR model fits more stable and control the population structure well than others (Supplementary Fig.14). Considering the lower error rates of ISR, those result promises to reveal genes that so far could not have been identified and more generally again shows the vast potential of ISR including its applicability to others species.

We further applied ISR to reanalyzed the rice dataset of grain length trait that owns a strong population structure, which the germplasm collections from all around the word^60^. After processing the data, including filtering for missing data, minor allele frequencies(MAF <0.05), the data were composed of m = 464,831 SNPs and n = 1,132 individuals. The data was previously reanalyzed to show the usefulness of the mixed model(EMMAX method^61^). Moreover, we used the same settings for mixed-model estimation here. We use the significance threshold level of 0.05 Bonferroni correction (P<1.08E-07) for comparative purposes and the significant SNPs for grain length trait identified by ISR and the others seven methods(except all above mentioned, also including the EMMAX^61^ and FASTLMM^49^ methods) are listed in (Supplementary Table.6). Here, all samples were evaluated together, and we can see two major GWAS peaks associated with grain length, one on chromosome 3 and one on chromosome 5 detected by the single-locus mixed model including GEMMA, EMMAX, FASTLMM, and CMLM methods. However, only the FASTLMM identified more than four SNPs in chromosome 10 (one) and 12 (three). The most significant SNPs were SNP-3.16732086 and SNP-5.5371749 from each of the major peaks on chromosomes 3 and 5, except for FASTmrEMMA, both identified by other methods (Supplementary Fig.15). Compared with the top ten SNPs detected by ISR, both different detected the same by GEMMA (two), EMMAX (two), FASTLMM (three), CMLM (two), FarmCPU (six), MLMM (four), and FASTmrEMMA (two).

## Discussion

Over the recent years, the prestigious GWAS methods development has been through several milestones from the single-locus model (mainly was a mixed model, such as EMMA^16^) to the multi-loci model (recently, BLINK^62^). Improvement of the LMM-based association approach has been proposed (included single-locus and multiple-loci linear model)^48,49,58,61,63^. All improvements are based on the assumption that population structure correction along with its negative effects cannot be entirely avoided (Supplementary Table 5, 6, and Supplementary Figs. 14, 15), part of the reasons that the trait is not approximately following an infinitesimal genetic architecture^63^. Otherwise, population structure leads to linkage disequilibrium (LD) between physically unlinked regions and thereby to correlations between markers of these regions. However, the multiple-loci linear model can conquer LD (Supplementary Fig.12b). The problem of population structure in GWAS is best viewed as one of model misspecification. Single-locus tests of the association are the wrong model to use when the trait is not attributable to a single locus.

Here, we have presented a novel statistical regression model. Based on that, derive a new set of methodologies, called a ‘multiple-locus linear model’ (ISR), and using it to the genetic association of complex traits. The method includes a significant locus in the model via a new iterative screen regression approach, which was continually changing the variable select criterion of the model at each step. ISR is a combined method with two stages, each of which needs a critical p-value. In the first step, a critical p-value 0.01 (methods default) (or 0.005 and 0.001, Supplementary Note Fig.2) were compared to obtain the best one. We divided variants into three types (Supplementary Note Fig.5 and Fig.1) and combined the expansion and contract screen procedure (Fig.1). Population structure is not species-specific but can be found in populations of any type. Moreover, we want to point out that the formulation of ISR can also be easily extended to accommodate other fixed effects (e.g., age, sex, or genotype principal components) that can be used to account for sample non-independence due to other genetic or shared environmental factors and similar to the LMM or LM approach. Otherwise, add fixed effects had no influenced the detected power (Supplementary Fig.17). ISR without fitting PCs as covariates still outperformed MLM that incorporated PCs as covariates (Supplement Fig.15). Fitting appropriate PCs as covariates in ISR further improved statistical power (Supplement Fig.17).

Our simulations showed that ISR is still very conservative, which indicates that such further development could lead to even more powerful methods. However, already in the current form, ISR correctly accounted for polygenic inheritance and facilitated to overcome the requirement for population structure correction. In any way, independent of the actual method, associating correlated markers will always be superior to the single-marker association. They are more consistent with the nature of quantitative traits (Supplementary Fig.15). ISR demonstrated that not only promising performance regarding power versus FDR and FPR in comparison with a single-locus mixed model scan(CMLM^22^, GEMMA^26^, FaSTLMM, and PLINK-Fisher) and multiple loci mixed model scan (FarmCPU^28^, MLMM^58^, and FASTmrEMMA^44^) but also had a higher accuracy effect estimated (PVE estimated). Particularly applying a relative conservative threshold, which can be effectuated with one of the proposed model quality criteria. ISR is not without its limitations. Perhaps the most significant burden is its computational cost. However, it still comparable with MLMM, CMLM, and faster than FASTmrEMMA (Supplementary Fig.19), when the individuals are a significant increase. On the other hand, it was built by MATLAB language, as we were known, which the M language with lower computer speed than other languages, such as, C and C++ and so forth, consider ISR itself, though both R and C++ program under development. Also, we will consider it combined with other technology like QTCAT^45^ to improve the power and achieve a lower false discovery rate.

We have focused on one application of ISR— genetic association of phenotypes. We were applying ISR to real data from A. thaliana, rice, and mice. Compared with other methods, our methods detected more known and unknown candidate genes (Supplementary Table 3), moreover, in contrast to the single-locus model that the visualization of the multiple-loci model (ISR, FarmCPU, and MLMM) results which the Manhattan plot and QQ plot showed reasonably and better illustrates the nature of quantitative traits (Supplementary Fig.15). Being with the marker density is multiply increasing, and no longer exist spikes and surprising^28^.

## Methods

### Overview of ISR

We provide a brief overview of ISR here. Detailed methods and algorithms are provided in the Supplementary Note. To model the relationship between phenotypes and genotypes, we consider the following multiple regression model:

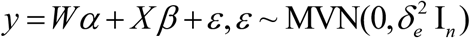

where *y* is an *n*-vector of phenotypes measured on *n* individuals; W=(w_1_,w_2_…w_c_) is an *n* by *c* matrix of covariates(fixed effects) including a column of ones for the intercept term; *α* is a *c*-vector of coefficients; X is an *n* by *p* matrix of genotypes; β is the corresponding *p*-vector of effect sizes; ε is an *n*-vector of residual errors where each element is assumed to be independently and identically distributed from a normal distribution with variance 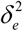; I_n_ is an *n* by *n* identity matrix and MVN denotes multivariate normal distribution. We used the proposed repeatedly screening stepwise linear regression model—effect size estimates obtained by the least-square method (LSM) and F-test P values for each SNP. The SNP with the most significant association is then added to the model as a cofactor for the next step. Combined the proposed repeatedly screening stepwise regression process, which makes it useful when *p*≫*n* (when the number of SNPs is much greater than the number of individuals).

We also proposed a new model selection criteria (RIC Fig.1) to select the most appropriate model (Supplementary Note). Without using the classic Bayesian information criterion (BIC)^64^ or Akaike information criterion (AIC)^65^, because they are too tolerant in the context of GWAS^58^, allowing for too many loci in the model.

### Simulations

GWA data from a set of 214,051 single-nucleotide polymorphism markers which surveys 248,584 SNPs after quality control, where were genotyped for 1,307 diverse Arabidopsis accessions showing strong population structure^66^ were used to perform two simulation experiments. Also, another outbred CFW (Carworth Farms White) mice population that including a set of 92,734 single-nucleotide polymorphism markers which were genotyped 1,161 individuals were also used to perform two simulation experiments. The human dataset derived from PLINK2^50^ included two real human genotype datasets. The first dataset included 1000 samples and 100000 makers (SNPs); The second included 10000 samples(6000 cases and 4000 control) and 88058 markers(SNPs), and only included in 19, 20, 21, and 22 chromosomes. The purpose was to compare ISR with the single-locus model methods (CMLM, GEMMA, LM) and the multi-locus model method (FarmCPU, FASTmrEMMA, MLMM). While for the human dataset, we only compare with FarmCPU, FaSTLMM^49^, and PLINK(version 1.9, and using Fisher’s exact test for association)^50,67^.

For the Arabidopsis dataset, the first two simulation experiments, a set of 20,000 SNPs and 1307 individuals were randomly sampled from the full dataset, seeing the density plot of SNPs (Supplementary Fig.20).

#### Scenario I

For the simple traits, following ^44,46,68^, we fixed two randomly chosen causal SNPs from each chromosome that were used to generate 100 phenotypes, where the phenotypes are simulated by the simple additive genetic model as following:

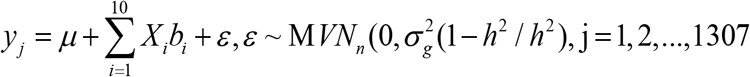

Where the average *μ* and heritability (total proportion of phenotypic variation explained) *h*^2^ were set at 10 and 0.25, respectively. The 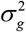 is the empirical variance of *X*_*i*_ *β*_*i*_ (i = 1, 2,…,10) and effects *β*_*i*_ (i = 1, 2,…,10) were generated from a normal distribution with means is 0 and variance is 4, where effects were 2.2386, −1.6089, 1.4445,−1.3338, −1.8779, 1.6808, −1.0891, 2.4238, 2.1443 and 1.8481, respectively (supplementary table1).

#### Scenario II

For the complex traits, following ^33^, we used an additive model with 100 randomly sampled causal SNPs having effect sizes *β*_*i*_ (i = 1, 2,…,100) drawn from an exponential distribution with a rate of 1. An additional random deviation *ε* was added, drawn from a normal distribution with a mean of zero and scaled identity matrix as a covariance matrix to fix the trait heritability *h*^2^ to 0.25, 0.5, and 0.75. For each phenotypic heritability, 100 phenotypes were simulated, the model as follows:

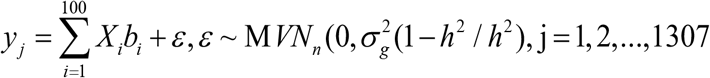

For the outbred CFW mice dataset, the first two simulation experiments, a set of 20,000 SNPs and 1161 individuals, were randomly sampled from the full dataset, seeing the density of SNPs (Supplementary Fig.21).

#### Scenario III

The first 100 phenotypes including 50 markers were randomly selected as causal loci. We assigned an additive effect randomly drawn from a standard normal distribution and added a random environmental term, such that *h*^2^ of the simulated traits was 0.25, 0.5, 0.75. Where the additive genetic model simulates the phenotypes as following:

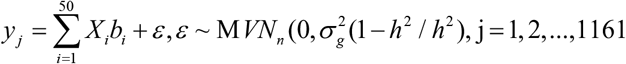

#### Scenario IV

The second 100 phenotypes used all CFW mice dataset that including 100 markers were randomly selected as causal loci, respectively. We also assigned an additive effect randomly drawn from a standard normal distribution and added a random environmental term, where the *h*^2^ of the simulated traits only was 0.5, here.

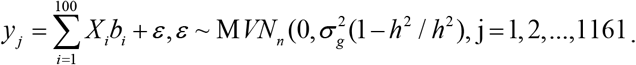

#### Scenario V

two 100 phenotypes used human dataset^50^ that including 100 markers were randomly selected as causal loci, respectively. We also assigned an additive effect randomly drawn from a standard normal distribution and added a random environmental term, where the *h*^2^ of the simulated traits only was 0.5, here.

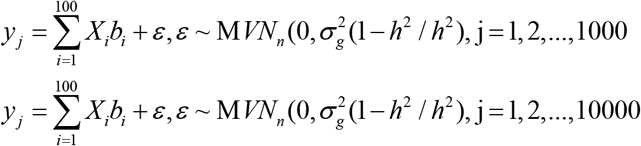

### Receiver operating characteristics

For each scenario, we examined statistical power (TPR, Ture Positive Rate) under different levels of FDR and FPR (Type I error). We defined FDR as the proportion of false positives among the total number of positives identified. Defined FPR as the proportion of false positives among the total number of negatives identified. Described the relationship between TPR and FDR or FPR uses the receiver operating characteristic (ROC) curves^69^. The method with a larger area under the curve (AUC) is preferred over the method with a smaller AUC.

### Other methods

We compared the performance of ISR mainly with six existing methods: (1) CLMM^22^ as implemented in the GAPIT^70^ R package; (2) LMM^66^ and LM as implemented in the GEMMA software (version 0.95alpha); (3) FarmCPU^28^ as implemented in the FarmCPU R package; (4) FASTmrEMMA as implemented in the mrMLM R package; (5) MLMM^33^ as implemented in the MLMM R package. We used default settings to fit all these methods and the details, as above stated.

## Code availability

ISR is available as an open-source MATLAB package at https://github.com/czheluo/ISR.

## Data availability

No data were generated in the present study. The 1,307 diverse Arabidopsis accessions data included genotype and phenotype is publicly available at https://1001genomes.org/data-center.html or http://bergelson.uchicago.edu/. The outbred CFW mice of genotype and phenotype data are publicly available at https://github.com/pcarbo/cfw. The human dataset derived from PLINK2^50^ included two real human genotype datasets only for the simulations.

## Author contributions

Shiliang Gu conceived the study and supervised statistical aspects of this work. Shiliang Gu and Meng Luo developed the software. Meng Luo designed the experiment and performed the simulations and data analyses. Meng Luo wrote the manuscript, and Shiliang Gu approved the final manuscript.

## Additional information

Supplementary Information accompanies this paper.

