## Supplementary Figures for "A new approach of dissecting genetic effects for complex traits"

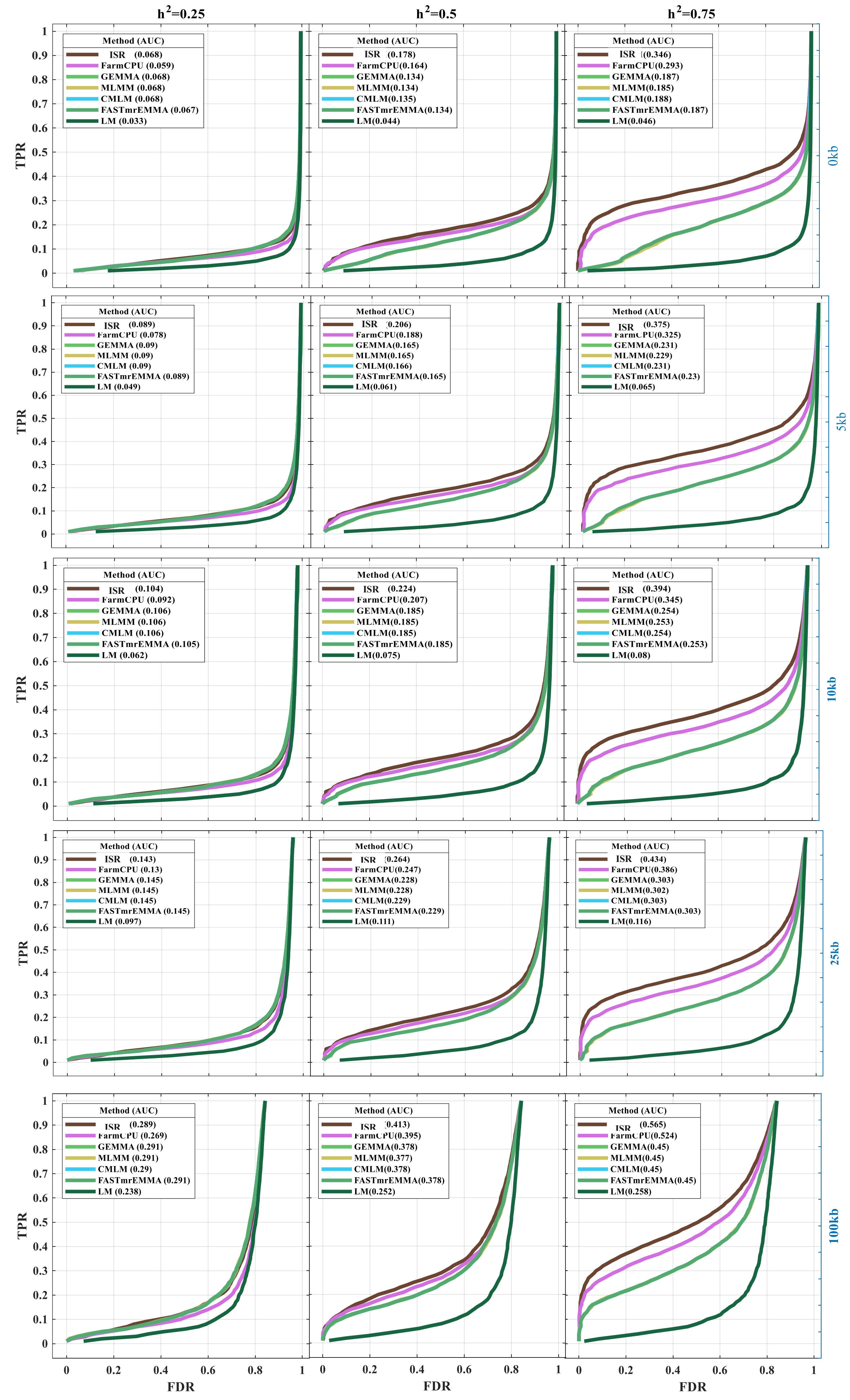

h2=0.25
h2=0.5
h2=0.75
TPR
FDR
0kb
5kb
10kb
25kb
100kb
Method (AUC)
ISR (0.068)
FarmCPU (0.059)
GEMMA (0.068)
MLMM (0.068)
CMLM (0.068)
FASTmrEMMA (0.067)
LM (0.033)
Method (AUC)
ISR (0.178)
FarmCPU(0.164)
GEMMA(0.134)
MLMM(0.134)
CMLM(0.135)
FASTmrEMMA(0.134)
LM(0.044)
Method (AUC)
ISR (0.346)
FarmCPU(0.293)
GEMMA(0.187)
MLMM(0.185)
CMLM(0.188)
FASTmrEMMA(0.187)
LM(0.046)
Method (AUC)
ISR (0.089)
FarmCPU (0.078)
GEMMA (0.09)
MLMM (0.09)
CMLM (0.09)
FASTmrEMMA (0.089)
LM (0.049)
Method (AUC)
ISR (0.206)
FarmCPU(0.188)
GEMMA(0.165)
MLMM(0.165)
CMLM(0.166)
FASTmrEMMA(0.165)
LM(0.061)
Method (AUC)
ISR (0.375)
FarmCPU(0.325)
GEMMA(0.231)
MLMM(0.229)
CMLM(0.231)
FASTmrEMMA(0.23)
LM(0.065)
Method (AUC)
ISR (0.104)
FarmCPU (0.092)
GEMMA (0.106)
MLMM (0.106)
CMLM (0.106)
FASTmrEMMA (0.105)
LM (0.062)
Method (AUC)
ISR (0.224)
FarmCPU(0.207)
GEMMA(0.185)
MLMM(0.185)
CMLM(0.185)
FASTmrEMMA(0.185)
LM(0.075)
Method (AUC)
ISR (0.394)
FarmCPU(0.345)
GEMMA(0.254)
MLMM(0.253)
CMLM(0.254)
FASTmrEMMA(0.253)
LM(0.08)
Method (AUC)
ISR (0.143)
FarmCPU (0.13)
GEMMA (0.145)
MLMM (0.145)
CMLM (0.145)
FASTmrEMMA (0.145)
LM (0.097)
Method (AUC)
ISR (0.264)
FarmCPU(0.247)
GEMMA(0.228)
MLMM(0.228)
CMLM(0.229)
FASTmrEMMA(0.229)
LM(0.111)
Method (AUC)
ISR (0.434)
FarmCPU(0.386)
GEMMA(0.303)
MLMM(0.302)
CMLM(0.303)
FASTmrEMMA(0.303)
LM(0.116)
Method (AUC)
ISR (0.289)
FarmCPU (0.269)
GEMMA (0.291)
MLMM (0.291)
CMLM (0.29)
FASTmrEMMA (0.291)
LM (0.238)
Method (AUC)
ISR (0.413)
FarmCPU(0.395)
GEMMA(0.378)
MLMM(0.377)
CMLM(0.378)
FASTmrEMMA(0.378)
LM(0.252)
Method (AUC)
ISR (0.565)
FarmCPU(0.524)
GEMMA(0.45)
MLMM(0.45)
CMLM(0.45)
FASTmrEMMA(0.45)
LM(0.258)

**Supplementary Figure 1. Performances of TPR (Power) versus FDR in the second simulation scenarios.** A receiver operating characteristic curve for seven methods were performed to test Power (TPR) versus FDR in the second simulation. Additive genetic effects controlled by 100 causal loci with three phenotypic heritabilities 0.25 (left), 0.5 (middle) and 0.75 (right), including ISR, FarmCPU, GEMMA, MLMM, CMLM, FASTmrEMMA and LM methods. The casual loci were randomly sampled from all the SNPs in each dataset. We examined the power under different levels of FDR and FPR. The causal SNPs were considered to be detected if an SNP within 0kb, 10kb, 25kb, and 100kb on either side was determined to have a significant association. Moreover, we measured the performance of detecting associations is using the area under the curve (AUC), where a method with a higher value indicates better performance.

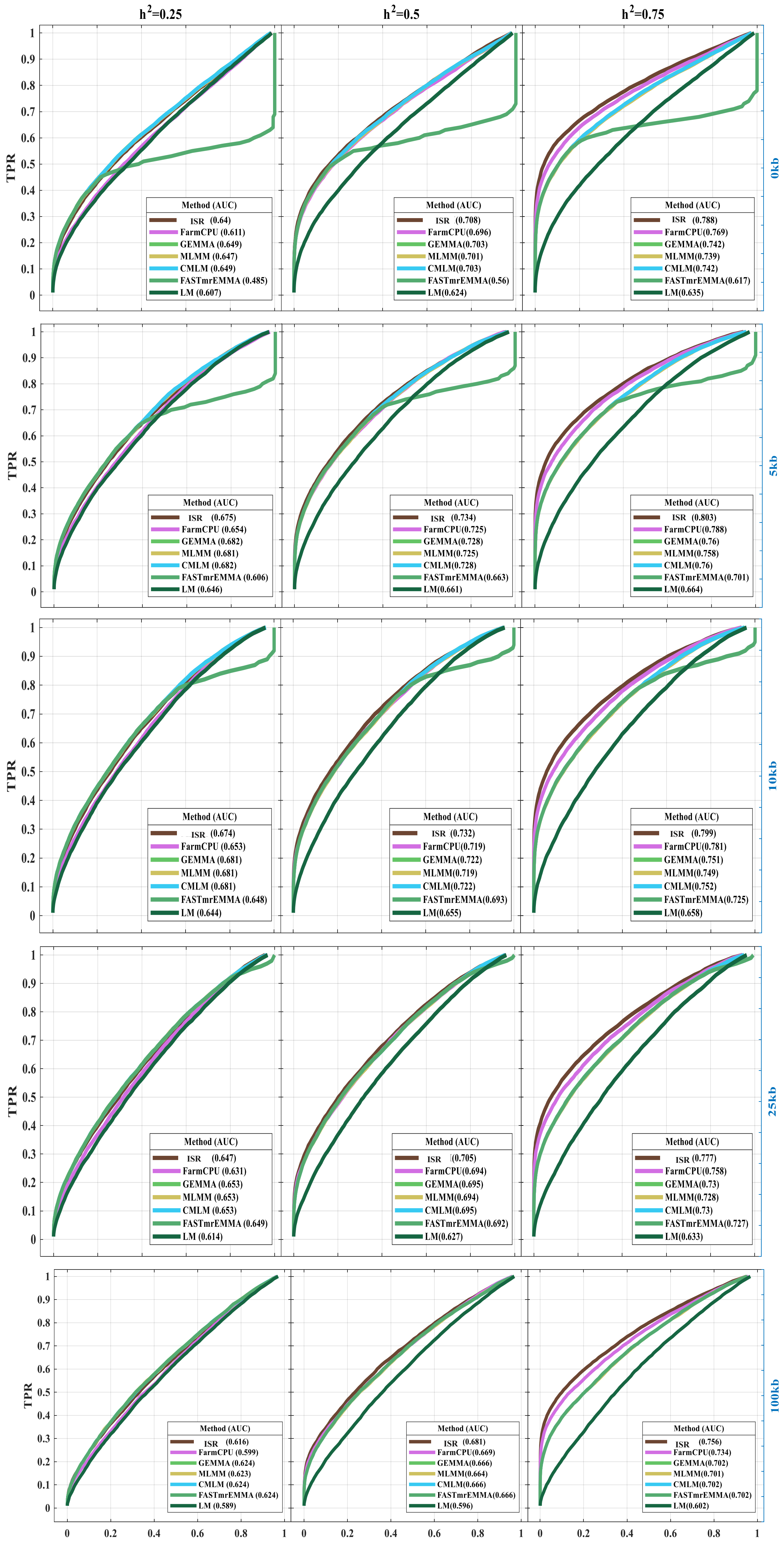


**Supplementary Figure 2. Performances of TPR (Power) versus FPR in the second simulation scenarios.** A receiver operating characteristic curve for seven methods were performed to test Power (TPR) versus FPR in the second simulation. Additive genetic effects controlled by 100 causal loci with three phenotypic heritabilities 0.25 (left), 0.5 (middle) and 0.75 (right), including ISR, FarmCPU, GEMMA, MLMM, CMLM, FASTmrEMMA and LM methods. The casual loci were randomly sampled from all the SNPs in each dataset. We examined the power under different levels of FPR. A causal SNP was considered to be detected if an SNP within 0kb, 10kb, 25kb, and 100kb on either side was determined to have a significant association; otherwise, it is considered a false positive. Moreover, we measured the performance of detecting associations is using the area under the curve (AUC), where a method with a higher value indicates better performance.

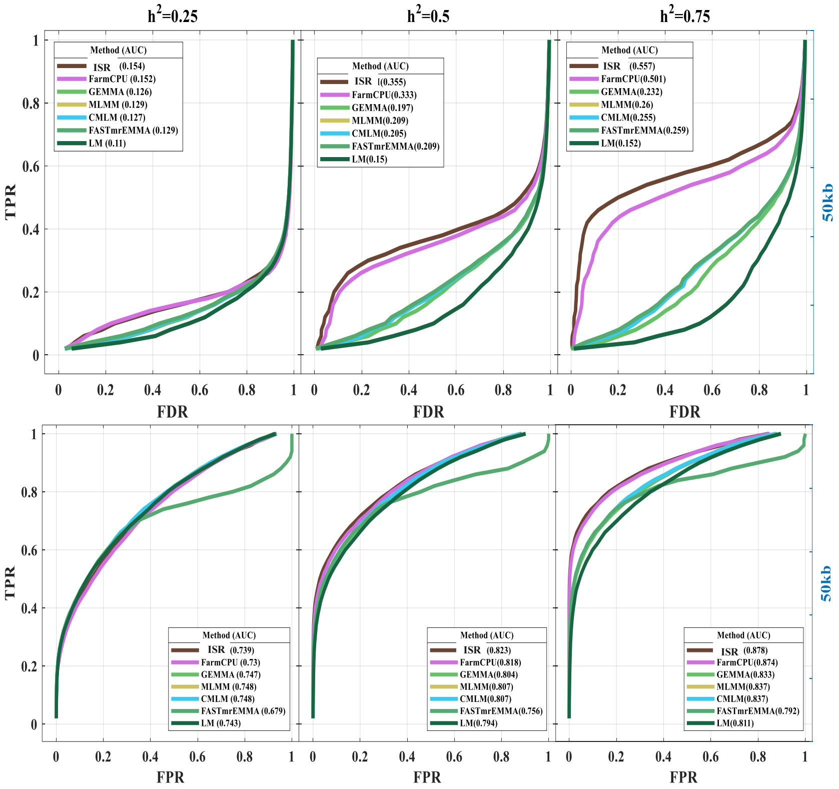


**Supplementary Figure 3. Performances of TPR (Power) versus FDR and FPR in third simulation scenarios.** A receiver operating characteristic curve for seven methods were performed to test Power (TPR) versus FPR in the third simulation. Additive genetic effects controlled by 50 causal loci with three phenotypic heritabilities 0.25 (left), 0.5 (middle) and 0.75 (right), including ISR, FarmCPU, GEMMA, MLMM, CMLM, FASTmrEMMA and LM methods. The casual loci were randomly sampled from all the SNPs in each dataset. We examined the power under different levels of FDR and FPR. A causal SNP was considered to be detected if an SNP within 50kb on either side was determined to have a significant association; otherwise, it is considered a false positive. Moreover, we measured the performance of detecting associations is using the area under the curve (AUC), where a method with a higher value indicates better performance.

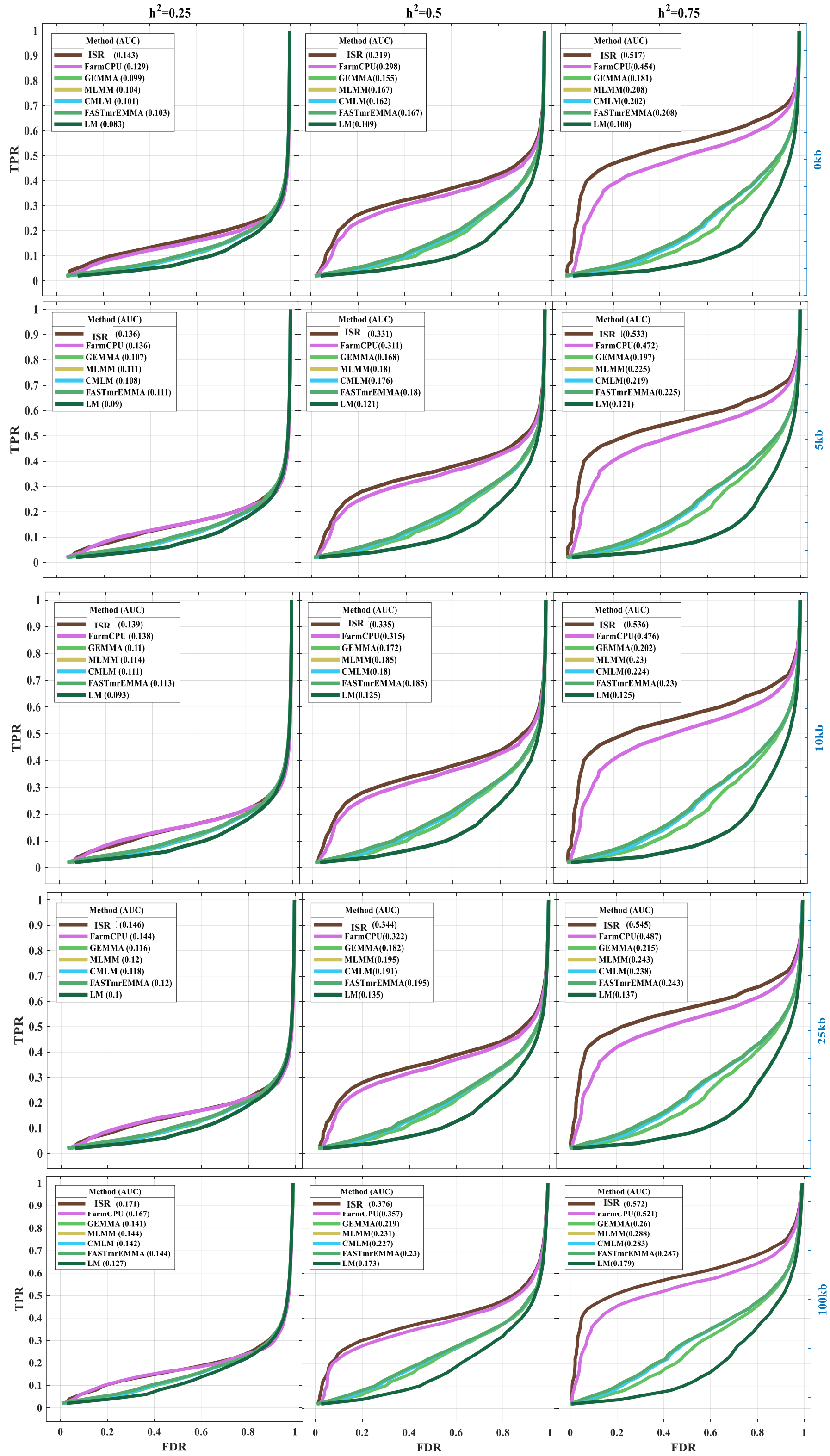
$h^2$=0.25
$h^2$=0.5
$h^2$=0.75
Method (AUC)
ISR (0.143)
FarmCPU (0.129)
GEMMA (0.099)
MLMM (0.104)
CMLM (0.101)
FASTmrEMMA (0.103)
LM (0.083)
Method (AUC)
ISR (0.319)
FarmCPU(0.298)
GEMMA(0.155)
MLMM(0.167)
CMLM(0.162)
FASTmrEMMA(0.167)
LM(0.109)
Method (AUC)
ISR (0.517)
FarmCPU(0.454)
GEMMA(0.181)
MLMM(0.208)
CMLM(0.202)
FASTmrEMMA(0.208)
LM(0.108)
0kb
Method (AUC)
ISR (0.136)
FarmCPU (0.136)
GEMMA (0.107)
MLMM (0.111)
CMLM (0.108)
FASTmrEMMA (0.111)
LM (0.09)
Method (AUC)
ISR (0.331)
FarmCPU(0.311)
GEMMA(0.168)
MLMM(0.18)
CMLM(0.176)
FASTmrEMMA(0.18)
LM(0.121)
Method (AUC)
ISR (0.533)
FarmCPU(0.472)
GEMMA(0.197)
MLMM(0.225)
CMLM(0.219)
FASTmrEMMA(0.225)
LM(0.121)
5kb
Method (AUC)
ISR (0.139)
FarmCPU (0.138)
GEMMA (0.11)
MLMM (0.114)
CMLM (0.111)
FASTmrEMMA (0.113)
LM (0.093)
Method (AUC)
ISR (0.335)
FarmCPU(0.315)
GEMMA(0.172)
MLMM(0.185)
CMLM(0.18)
FASTmrEMMA(0.185)
LM(0.125)
Method (AUC)
ISR (0.536)
FarmCPU(0.476)
GEMMA(0.202)
MLMM(0.23)
CMLM(0.224)
FASTmrEMMA(0.23)
LM(0.125)
10kb
Method (AUC)
ISR (0.146)
FarmCPU (0.144)
GEMMA (0.116)
MLMM (0.12)
CMLM (0.118)
FASTmrEMMA (0.12)
LM (0.1)
Method (AUC)
ISR (0.344)
FarmCPU(0.322)
GEMMA(0.182)
MLMM(0.195)
CMLM(0.191)
FASTmrEMMA(0.195)
LM(0.135)
Method (AUC)
ISR (0.545)
FarmCPU(0.487)
GEMMA(0.215)
MLMM(0.243)
CMLM(0.238)
FASTmrEMMA(0.243)
LM(0.137)
25kb
Method (AUC)
ISR (0.171)
FarmCPU (0.167)
GEMMA (0.141)
MLMM (0.144)
CMLM (0.142)
FASTmrEMMA (0.144)
LM (0.127)
Method (AUC)
ISR (0.376)
FarmCPU(0.357)
GEMMA(0.219)
MLMM(0.231)
CMLM(0.227)
FASTmrEMMA(0.23)
LM(0.173)
Method (AUC)
ISR (0.572)
FarmCPU(0.521)
GEMMA(0.26)
MLMM(0.288)
CMLM(0.283)
FASTmrEMMA(0.287)
LM(0.179)
100kb
TPR
FDR

**Supplementary Figure 4. Performances of TPR (Power) versus FDR in third simulation scenarios.** A receiver operating characteristic curve for seven methods were performed to test Power (TPR) versus FDR in the third simulation. Additive genetic effects controlled by 50 causal loci with three phenotypic heritabilities 0.25 (left), 0.5 (middle) and 0.75 (right), including ISR, FarmCPU, GEMMA, MLMM, CMLM, FASTmrEMMA and LM methods. The casual loci were randomly sampled from all the SNPs in each dataset. We examined the power under different levels of FDR. A causal SNP was considered to be detected if an SNP within 0kb, 10kb, 25kb, and 100kb on either side was determined to have a significant association; otherwise, it is considered a false positive. Moreover, we measured the performance of detecting associations is using the area under the curve (AUC), where a method with a higher value indicates better performance.

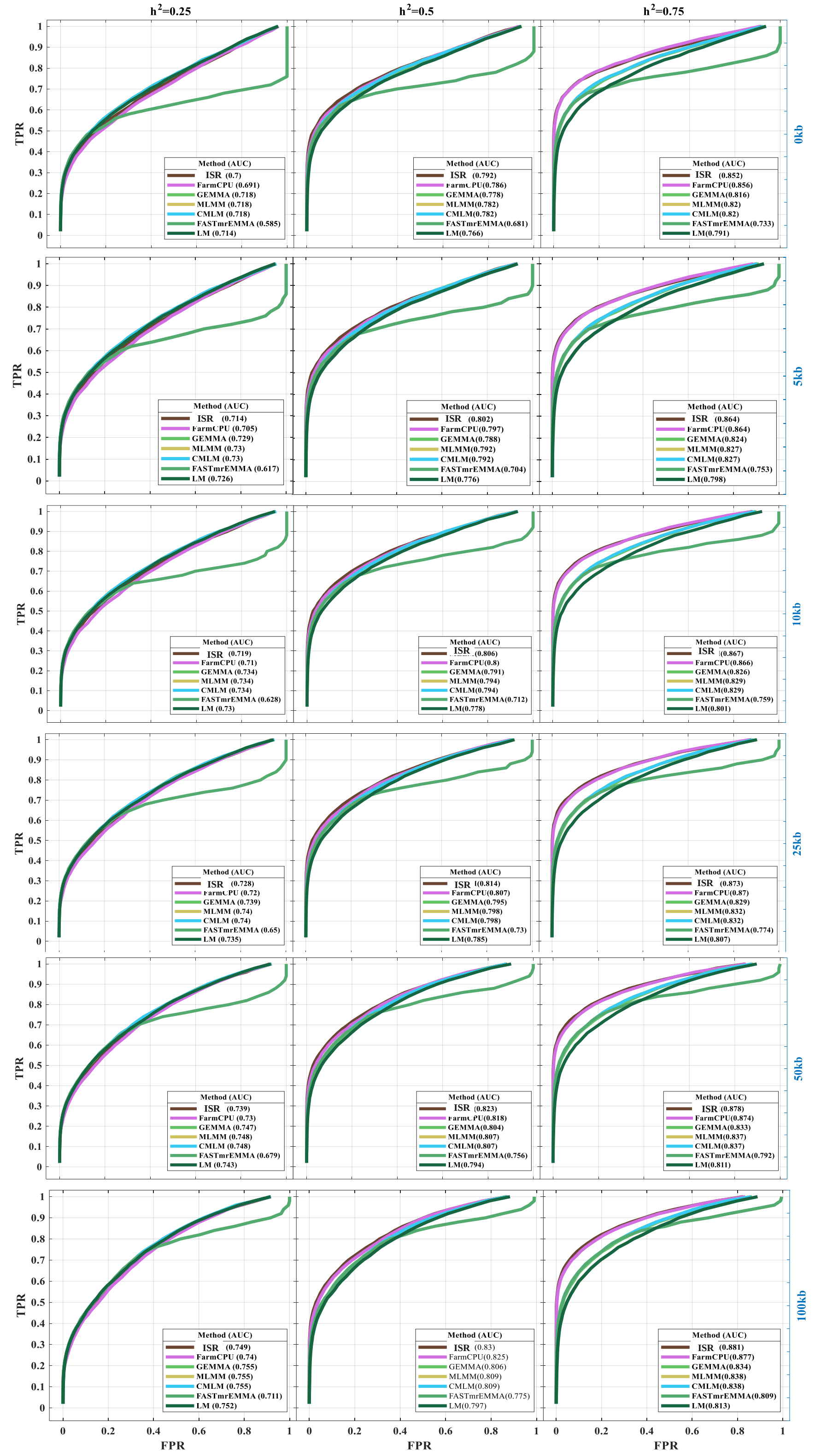
h2=0.25
h2=0.5
h2=0.75
0kb
5kb
10kb
25kb
50kb
100kb
TPR
FPR
Method (AUC)
ISR (0.7)
FarmCPU (0.691)
GEMMA (0.718)
MLMM (0.718)
CMLM (0.718)
FASTmrEMMA (0.585)
LM (0.714)
ISR (0.792)
FarmCPU(0.786)
GEMMA (0.778)
MLMM(0.782)
CMLM(0.782)
FASTmrEMMA(0.681)
LM(0.766)
ISR (0.852)
FarmCPU(0.856)
GEMMA(0.816)
MLMM(0.82)
CMLM(0.82)
FASTmrEMMA(0.733)
LM(0.791)
ISR (0.714)
FarmCPU (0.705)
GEMMA (0.729)
MLMM (0.73)
CMLM (0.73)
FASTmrEMMA (0.617)
LM (0.726)
ISR (0.802)
FarmCPU(0.797)
GEMMA(0.788)
MLMM(0.792)
CMLM(0.792)
FASTmrEMMA(0.704)
LM(0.776)
ISR (0.864)
FarmCPU(0.864)
GEMMA(0.824)
MLMM(0.827)
CMLM(0.827)
FASTmrEMMA(0.753)
LM(0.798)
ISR (0.719)
FarmCPU (0.71)
GEMMA (0.734)
MLMM (0.734)
CMLM (0.734)
FASTmrEMMA (0.628)
LM (0.73)
ISR (0.806)
FarmCPU(0.8)
GEMMA(0.791)
MLMM(0.794)
CMLM(0.794)
FASTmrEMMA(0.712)
LM(0.778)
ISR (0.867)
FarmCPU(0.866)
GEMMA(0.826)
MLMM(0.829)
CMLM(0.829)
FASTmrEMMA(0.759)
LM(0.801)
ISR (0.728)
FarmCPU (0.72)
GEMMA (0.739)
MLMM (0.74)
CMLM (0.74)
FASTmrEMMA (0.65)
LM (0.735)
ISR (0.814)
FarmCPU(0.807)
GEMMA(0.795)
MLMM(0.798)
CMLM(0.798)
FASTmrEMMA(0.73)
LM(0.785)
ISR (0.873)
FarmCPU(0.87)
GEMMA(0.829)
MLMM(0.832)
CMLM(0.832)
FASTmrEMMA(0.774)
LM(0.807)
ISR (0.739)
FarmCPU (0.73)
GEMMA (0.747)
MLMM (0.748)
CMLM (0.748)
FASTmrEMMA (0.679)
LM (0.743)
ISR (0.823)
FarmCPU(0.818)
GEMMA(0.804)
MLMM(0.807)
CMLM(0.807)
FASTmrEMMA(0.756)
LM(0.794)
ISR (0.878)
FarmCPU(0.874)
GEMMA(0.833)
MLMM(0.837)
CMLM(0.837)
FASTmrEMMA(0.792)
LM(0.811)
ISR (0.749)
FarmCPU (0.74)
GEMMA (0.755)
MLMM (0.755)
CMLM (0.755)
FASTmrEMMA (0.711)
LM (0.752)
ISR (0.83)
FarmCPU(0.825)
GEMMA(0.806)
MLMM(0.809)
CMLM(0.809)
FASTmrEMMA(0.775)
LM(0.797)
ISR (0.881)
FarmCPU(0.877)
GEMMA(0.834)
MLMM(0.838)
CMLM(0.838)
FASTmrEMMA(0.809)
LM(0.813)

**Supplementary Figure 5. Performances of TPR (Power) versus FDR in third simulation scenarios.** A receiver operating characteristic curve for seven methods were performed to test Power (TPR) versus FDR in the third simulation. Additive genetic effects controlled by 50 causal loci with three phenotypic heritability 0.25 (left), 0.5 (middle) and 0.75 (right), including ISR, FarmCPU, GEMMA, MLMM, CMLM, FASTmrEMMA and LM methods. The casual loci were randomly sampled from all the SNPs in each dataset. We examined the power under different levels of FDR. A causal SNP was considered to be detected if an SNP within 0kb, 10kb, 25kb, and 100kb on either side was determined to have a significant association; otherwise, it is considered a false positive. Moreover, we measured the performance of detecting associations is using the area under the curve (AUC), where a method with a higher value indicates better performance.

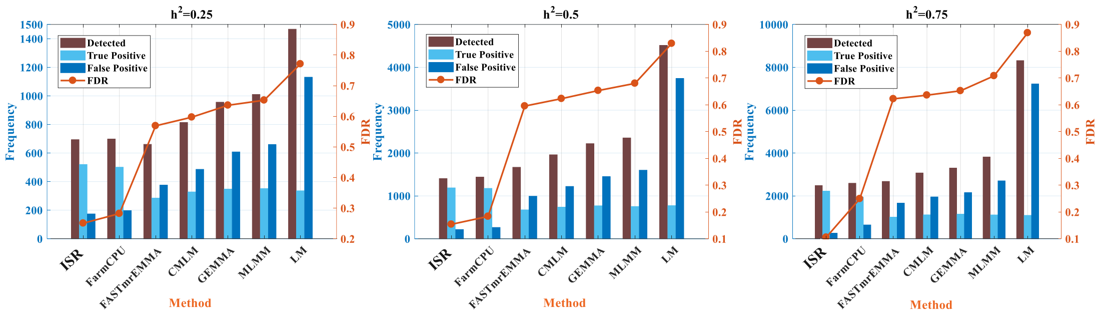


**Supplementary Figure 6. Performance of seven methods under different heritabilities in the third simulation scenarios.** The panel displays the numbers of detected, true positive, and false-positive SNPs that passed a threshold of 0.05 after a Bonferroni multiple test correction. A positive SNP is a true positive if a QTN is within a 50kb distance; otherwise, it is a false positive. The line chart displays FDR for each method.

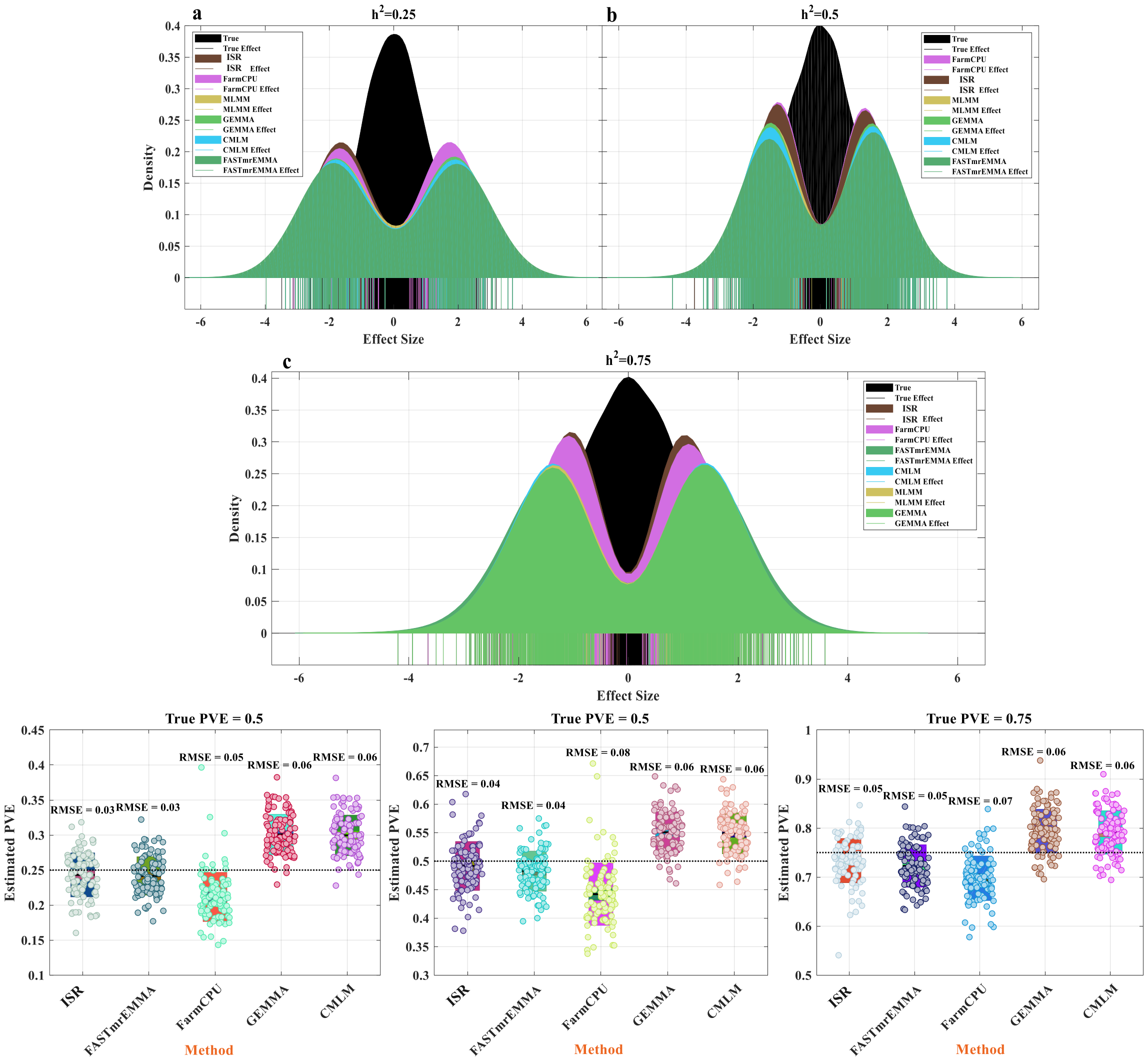


**Supplementary Figure 7. Comparison of detected effect and PVE estimates from five methods in the third simulation scenarios.** The distribution of all simulated effects (all true effect) and the distribution of effects of loci identified (50 casual loci for every 100 simulations, and only true positive) by six methods. The solid line shows the effect size by different methods. (**a**) The phenotype with 25% of PVE, (**b**) the phenotype with 50% of PVE, (**c**) the phenotype with 75% of PVE. The bottom boxplot has explained the variance of the loci effect estimated by ISR, FASTmrEMMA, FarmCPU, GEMMA, and CMLM within the 100 simulations. Comparing ISR, the performance of estimating PVE with the other four methods was using the root of mean square error (RMSE), where a method with lower value indicates better performance. The true PVEs are shown as the horizontal dash lines.

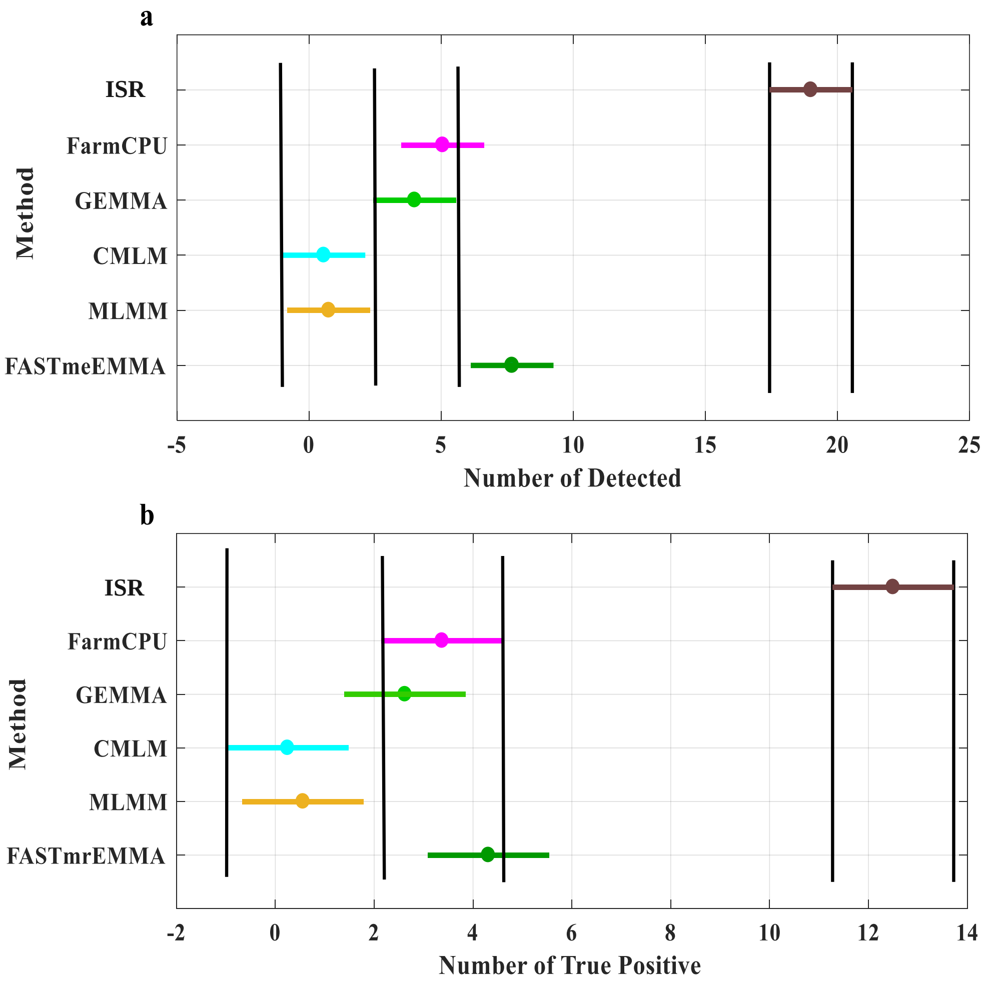


**Supplementary Figure 8. Multiple compare for the number of identified association SNPs with six statistics methods. (a)** plot show that reaches the detected number of SNPs by six methods, and (**b**) plot shows that true casual SNPs are in known candidate genes interval. All reported candidate genes and the reference literature could be found on the website (https://www.arabidopsis.org/index.jsp). The filled circle on the vertical line is the mean of the number of each group, and the length of the line represents the confidence interval. Two groups not sharing a horizontal dashed line are significantly different at Turkey 0.05.

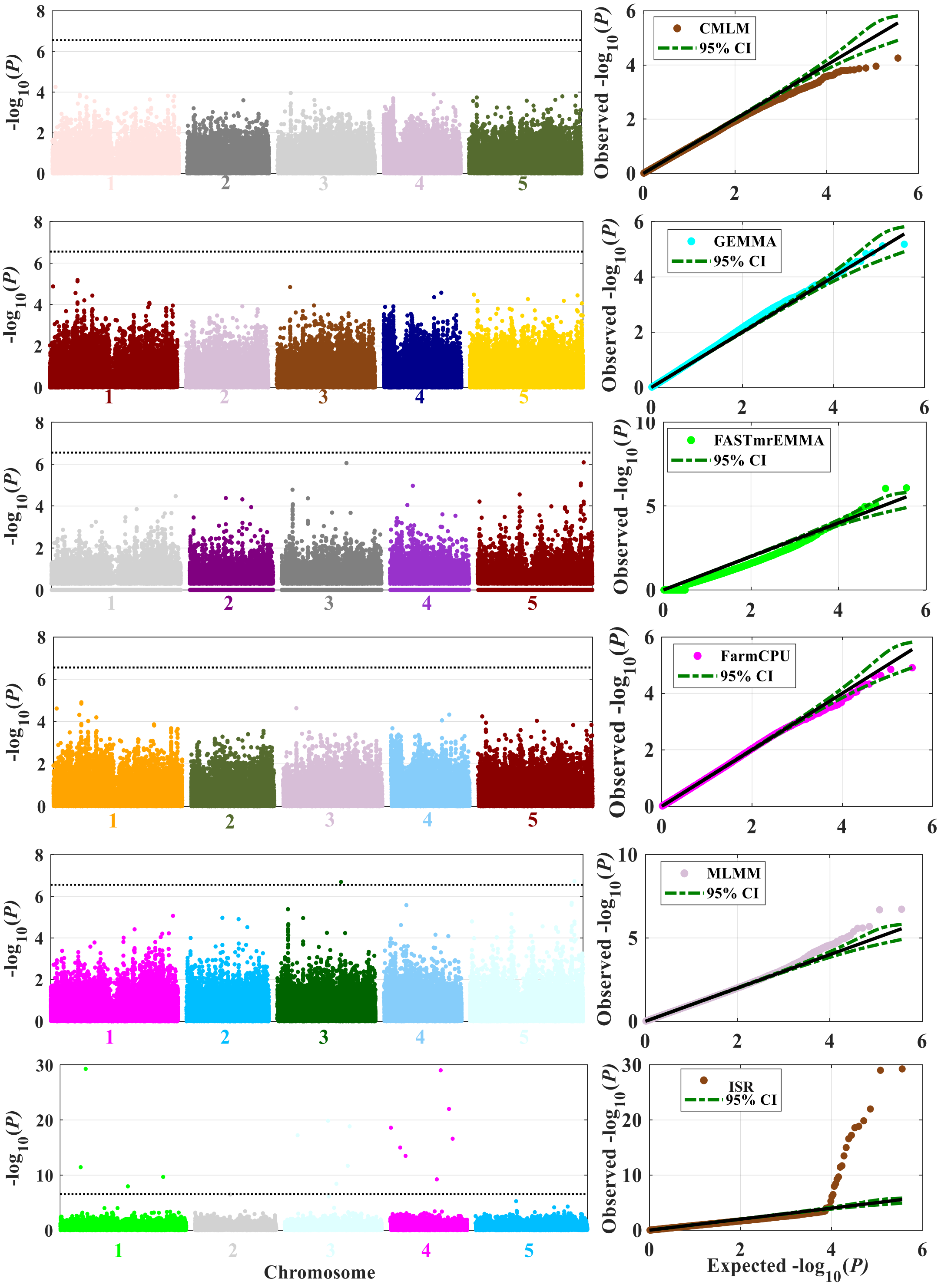

CMLM
95% CI
GEMMA
95% CI
FASTmrEMMA
95% CI
FarmCPU
95% CI
MLMM
95% CI
ISR
95% CI
-log10(P)
Observed -log10(P)
Expected -log10(P)
Chromosome

**Supplementary Figure 9. Association studies of bacterial disease resistance (At1 CFU2) in the *Arabidopsis thaliana* dataset.** The trait of At1 CFU2 was measured on 175 Arabidopsis thaliana individuals genotyped with 216,130SNPs and excluded non-polymorphic SNPs and SNPs with minor allele frequency less than 0.10 leave a total of 178,384 SNPs were used to calculate. Six statistical methods were employed to conduct the association studies: (1) FarmCPU; (2) CMLM; (3) GEMMA; (4) MLMM; (5) FASTmrEMMA; (6) ISR. The black dash line in the Manhattan plot shows that the value of Bonferroni multiple test correction (0.05/p, p is the number of SNPs), and the green dash lines in the QQ plot show that the 95% confidence intervals. All methods, except FarmCPU, CMLM, and GEMMA, and both identified associated SNPs after Bonferroni multiple test correction (Supplementary Table 3).

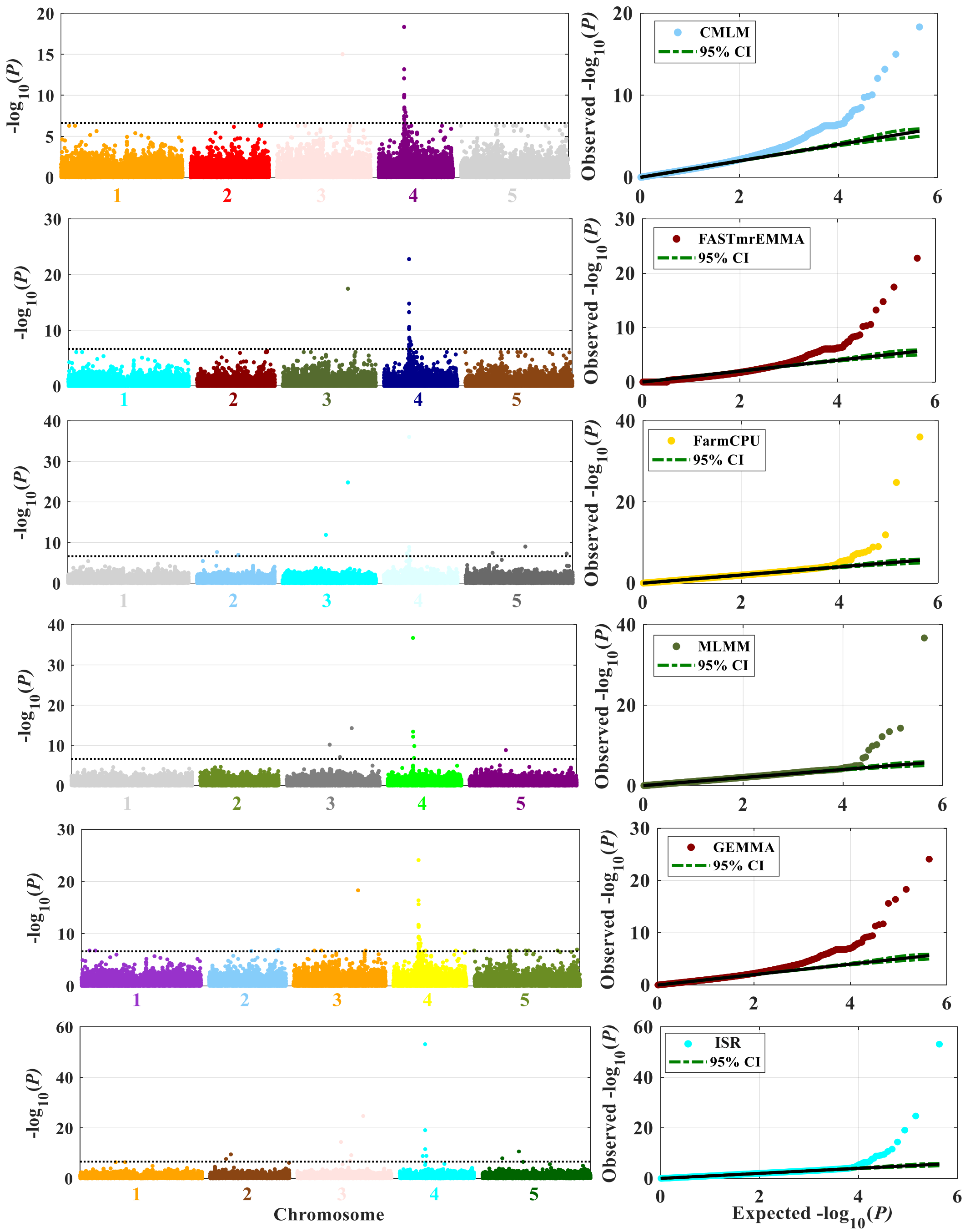


**Supplementary Figure 10. Association studies of leaf $Na^+$ accumulation in A.thaliana accessions.** The trait was measured on 349 Arabidopsis thaliana individuals genotyped with 214,051 SNPs, but excluded eight individuals with missing phenotypes. As above, six statistical methods were used to conduct the association studies. All methods that didn't include the PCs as covariates to control population structure. The dataset was derived from1.

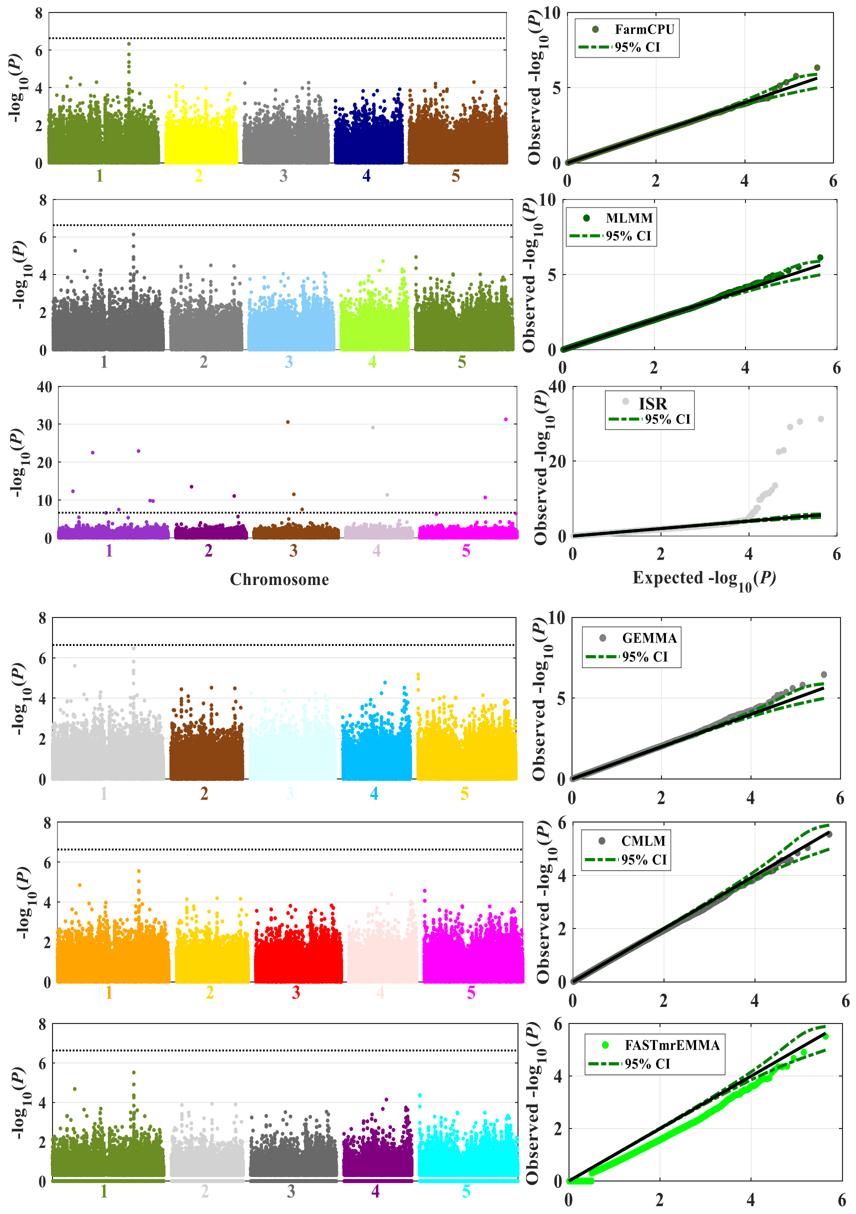


**Supplementary Figure 11. Association studies for cellular traits of meristem zone length in A. thaliana accessions.** The trait was measured on 201 Arabidopsis thaliana individuals genotyped with 214,051 SNPs. As above, six statistical methods were used to conduct the association studies. All methods included the first five PCs derived from all genetic markers as covariates to control population structure, PC-Select recommended by2.

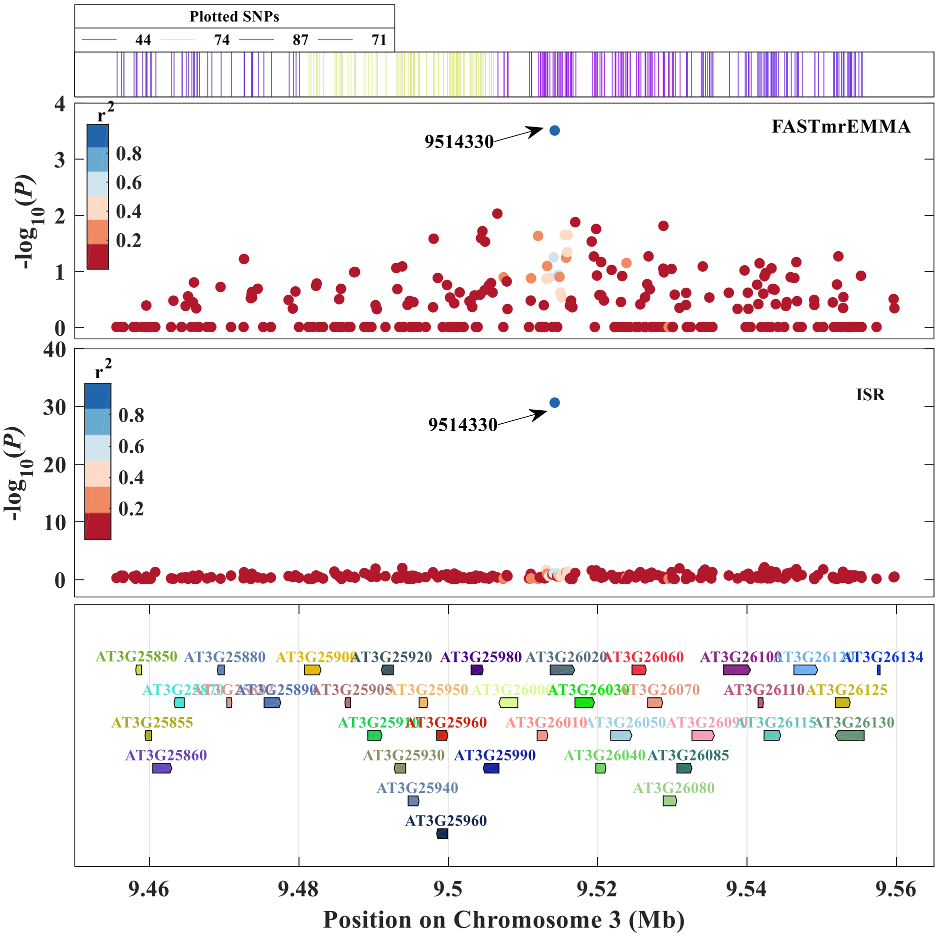


**Supplementary Figure 12. LocusZoom plots for genome-wide significant SNP are both found by ISR and FASTmrEMMA.** The locuszoom plots showing the zoom-in of the most significant SNP are both seen by FASTmrEMMA and ISR two methods. The points for each SNP are colored by the level of the linkage disequilibrium ($r^2$) with the index SNP, the SNP with the highest association to the quantitative trait meristem zone length.

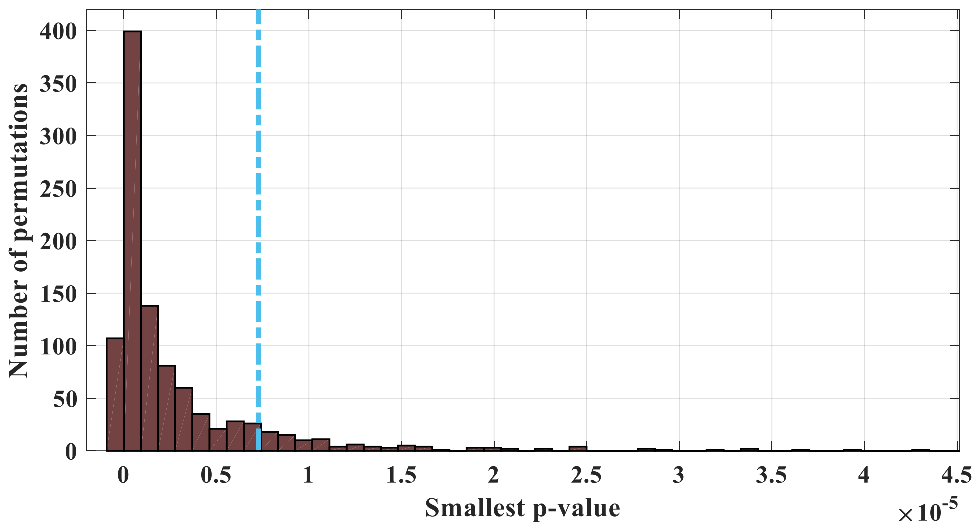


**Supplementary Figure 13. The results of permutation analysis for association mapping.** This plot summarizes the distribution of minimum p-values from mapping QTLs in 1,000 permuted data sets. The minimum p-value in each permuted data set is the smallest p-value from the 92,734 SNPs tested for association. This permutation test is meant to simulate the distribution of p-values under the null hypothesis. The dashed blue line depicts the 90th percentile of this distribution, which is approximately 7.7 × 10-6.

**BMD**

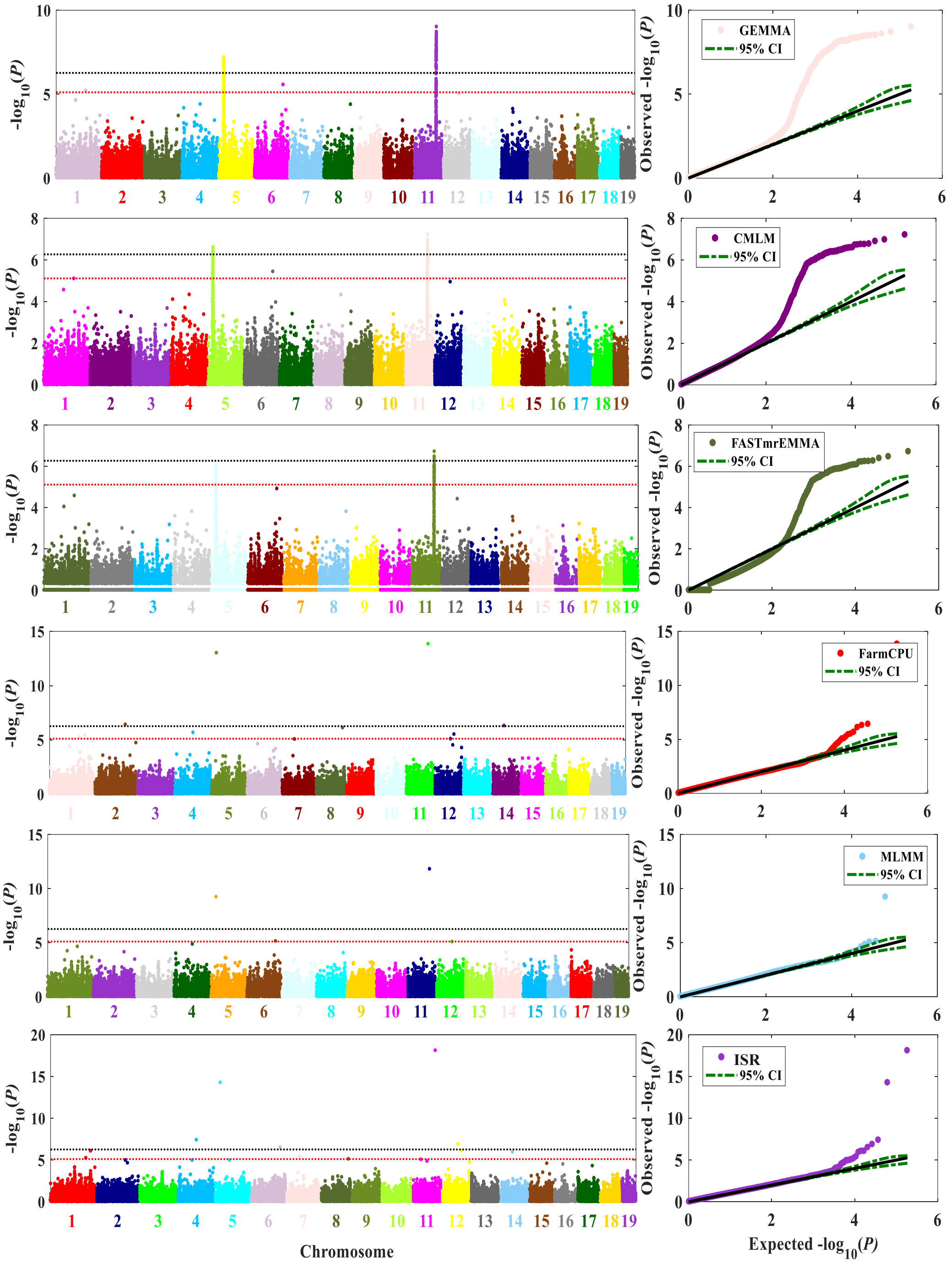
GEMMA
95% CI
CMLM
95% CI
FASTmrEMMA
95% CI
FarmCPU
95% CI
MLMM
95% CI
ISR
95% CI
-log10(P)
Observed -log10(P)
Chromosome
Expected -log10(P)

**EDL**

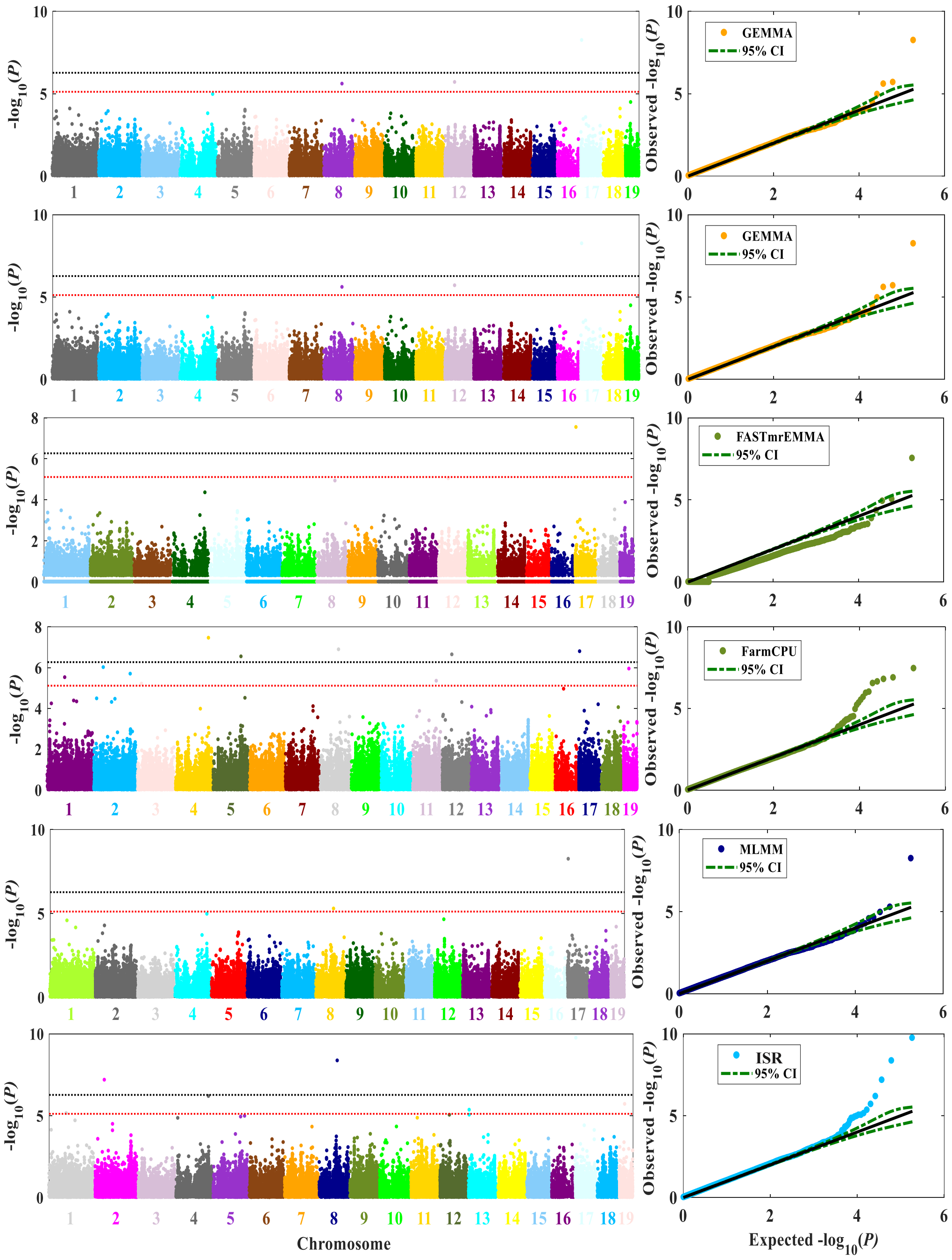

GEMMA
95% CI
GEMMA
95% CI
FASTmrEMMA
95% CI
FarmCPU
95% CI
MLMM
95% CI
ISR
95% CI
$-\log_{10}(P)$
Observed $-\log_{10}(P)$
Chromosome
Expected $-\log_{10}(P)$

**PPI12**

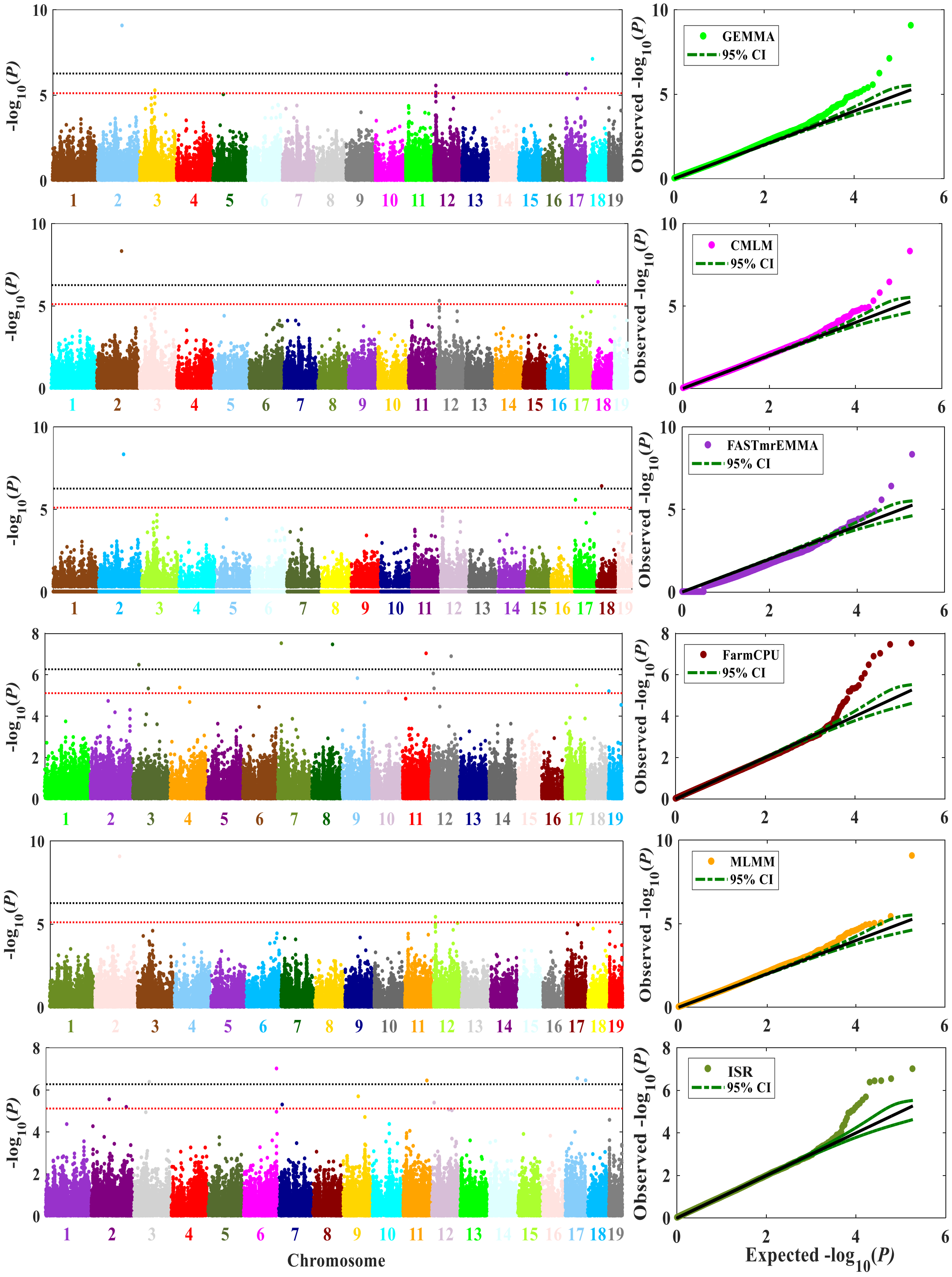

-log10(P)
Observed -log10(P)
GEMMA
95% CI
CMLM
95% CI
FASTmrEMMA
95% CI
FarmCPU
95% CI
MLMM
95% CI
ISR
95% CI
Chromosome
Expected -log10(P)

TA

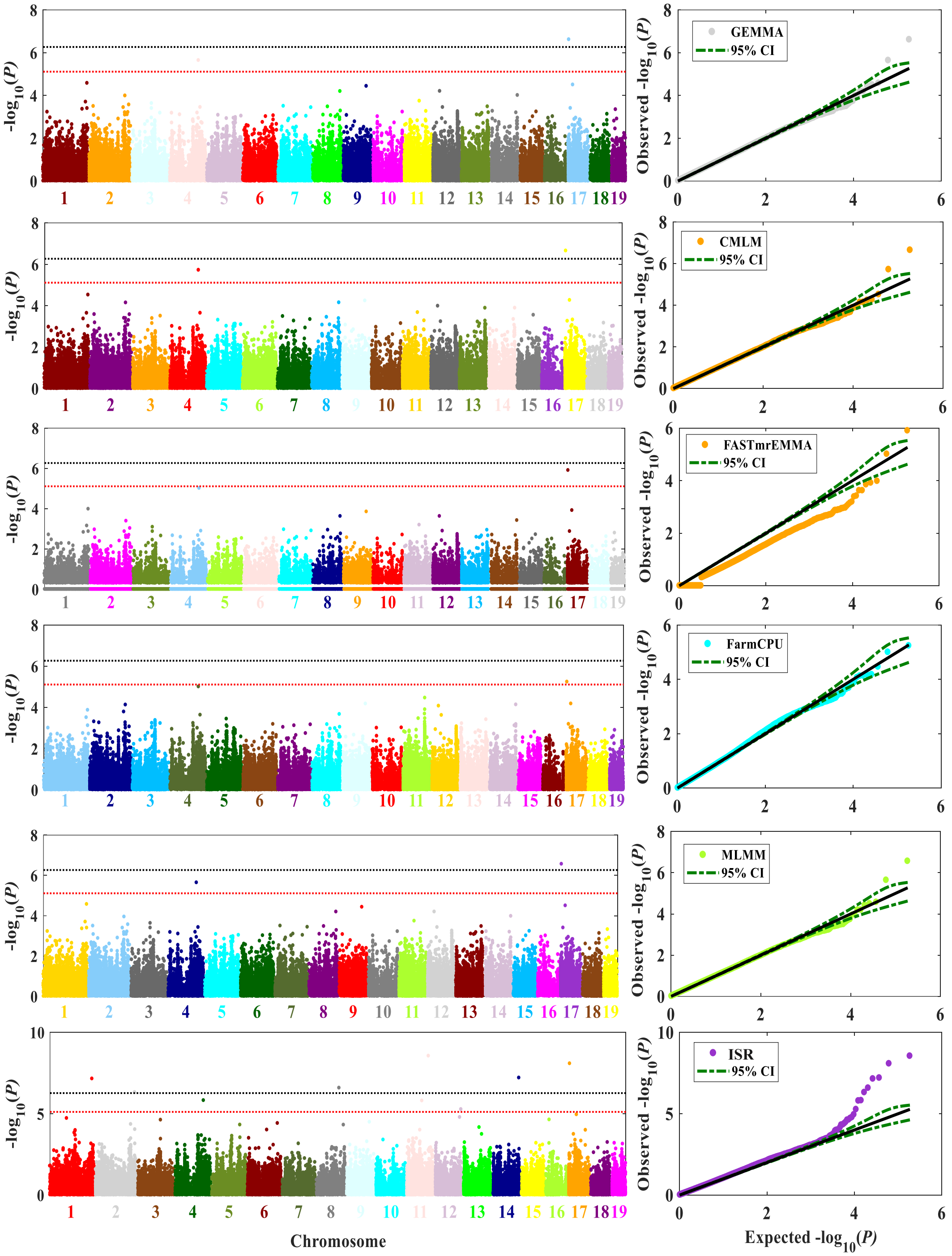
-log10(P)
Chromosome
GEMMA
CMLM
FASTmrEMMA
FarmCPU
MLMM
ISR
95% CI
Observed -log10(P)
Expected -log10(P)

### testiweight

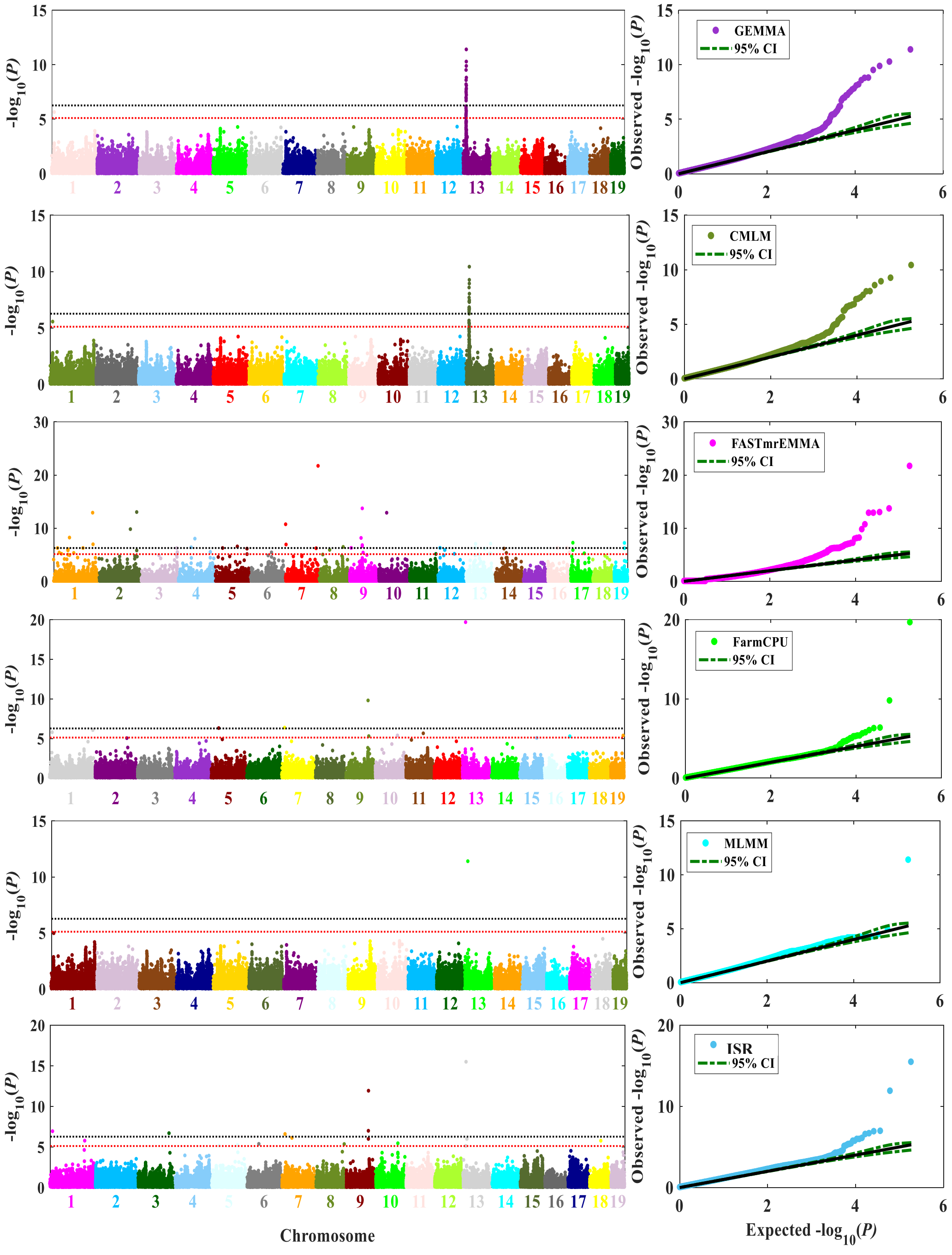

### Soleus

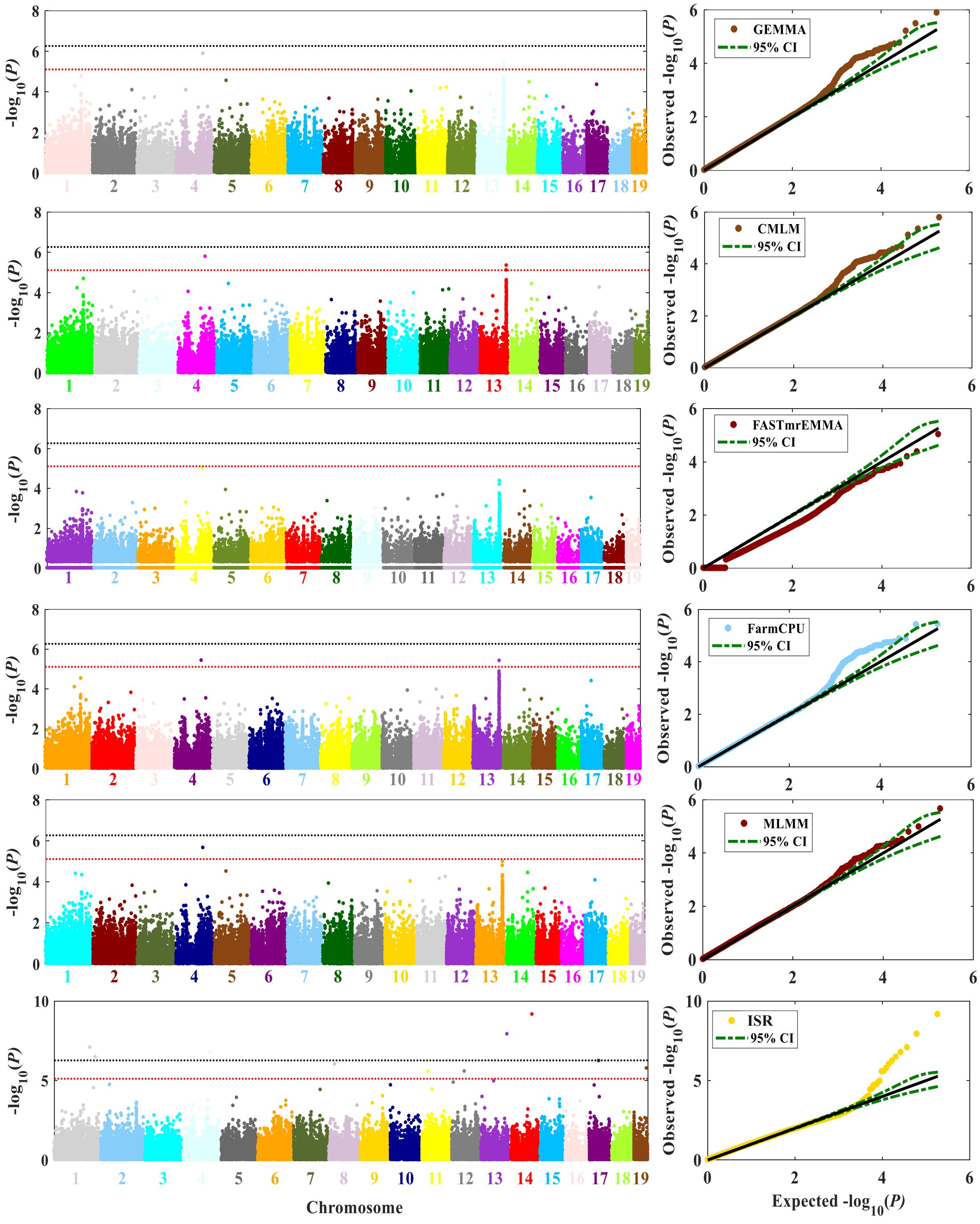

GEMMA
CMLM
FASTmrEMMA
FarmCPU
MLMM
ISR
95% CI
-log10(P)
Observed -log10(P)
Expected -log10(P)
Chromosome

Plantric

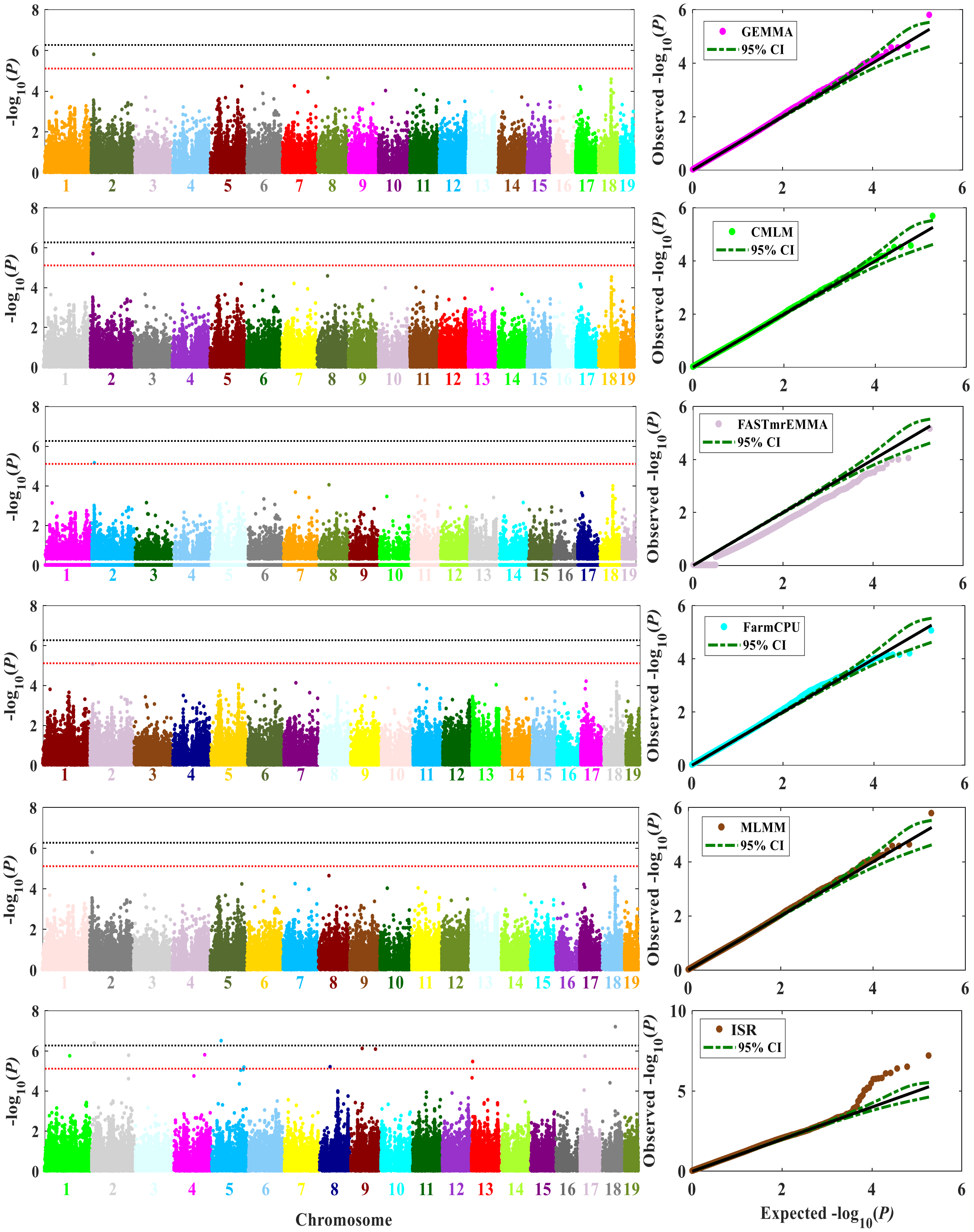

GEMMA
95% CI
CMLM
95% CI
FASTmrEMMA
95% CI
FarmCPU
95% CI
MLMM
95% CI
ISR
95% CI
-log10(P)
Observed -log10(P)
Chromosome
Expected -log10(P)

Fastgluose

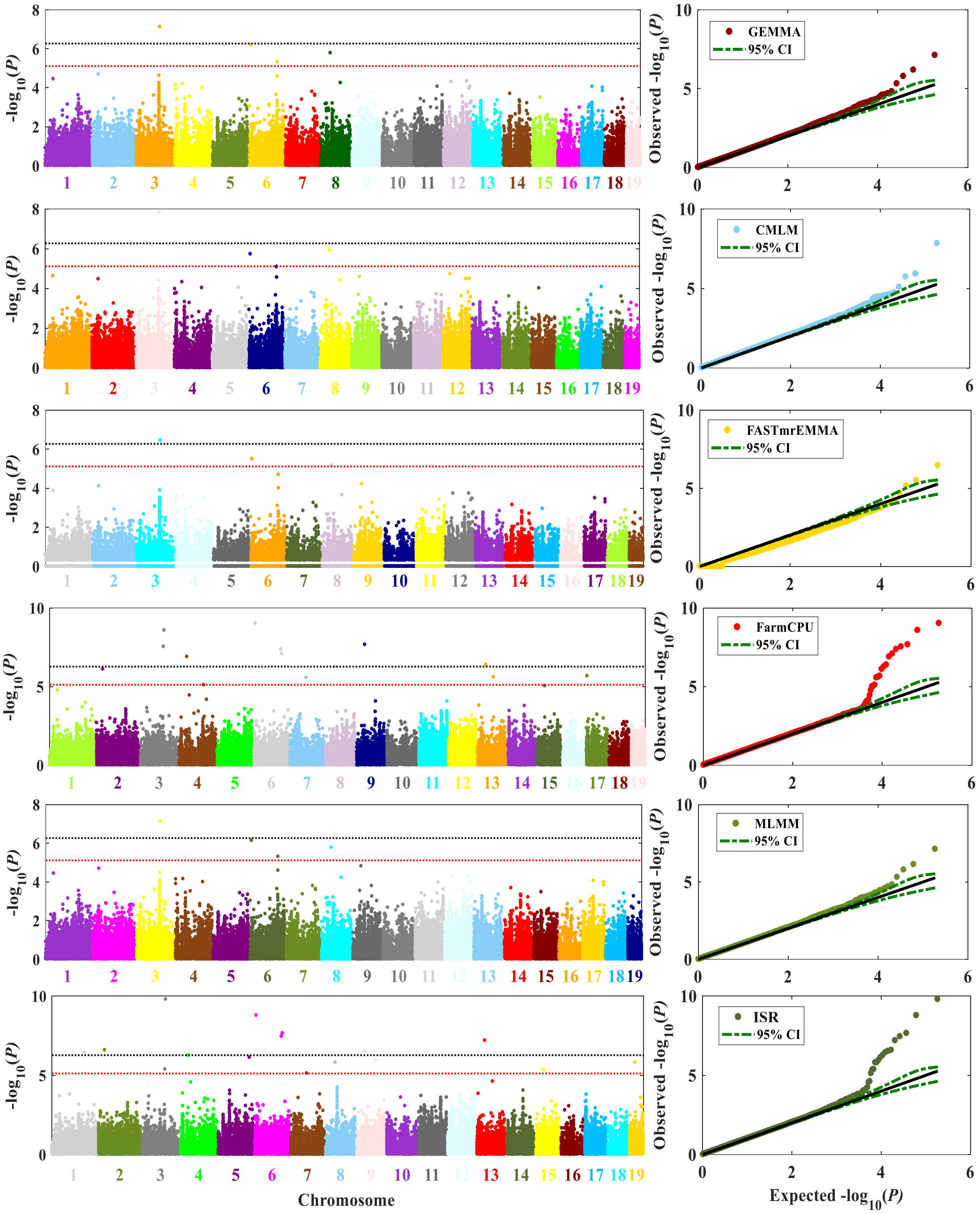

-log10(P)
Observed -log10(P)
GEMMA
95% CI
CMLM
95% CI
FASTmrEMMA
95% CI
FarmCPU
95% CI
MLMM
95% CI
ISR
95% CI
Chromosome
Expected -log10(P)

Sacweight

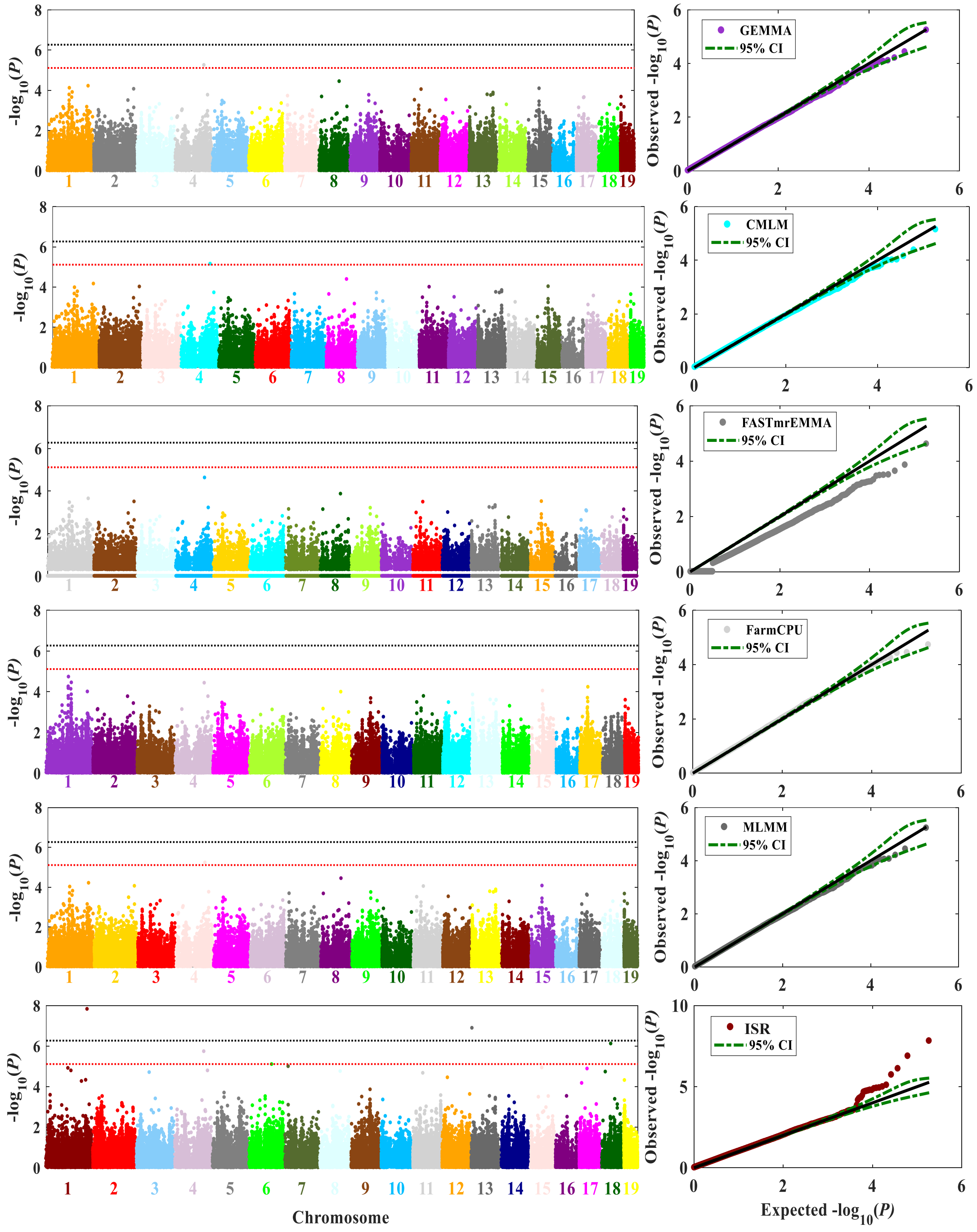
GEMMA
95% CI
CMLM
95% CI
FASTmrEMMA
95% CI
FarmCPU
95% CI
MLMM
95% CI
ISR
95% CI
$-\log_{10}(P)$
Observed $-\log_{10}(P)$
Chromosome
Expected $-\log_{10}(P)$

Taillenght

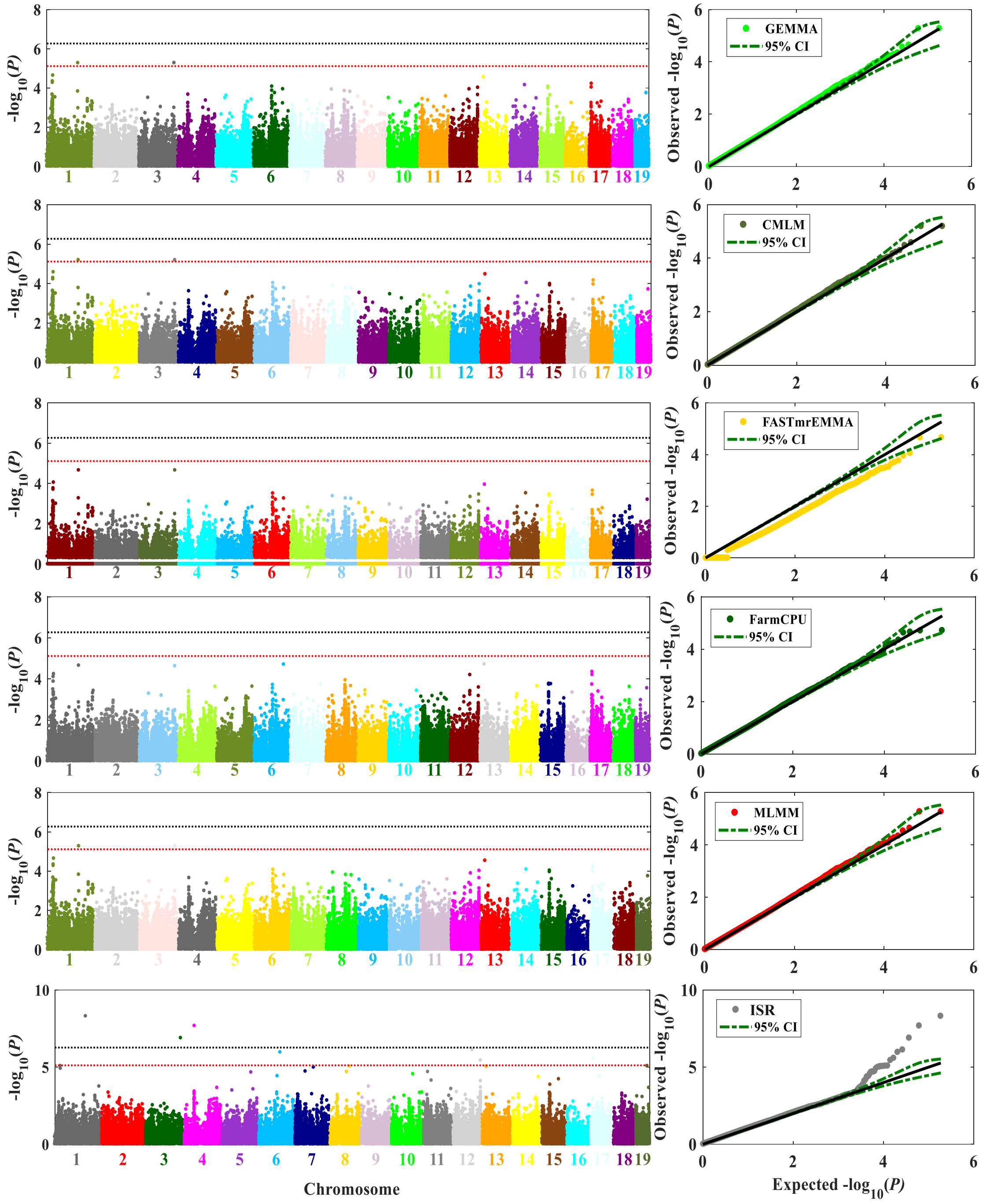


**Supplementary Figure 14. Genome-wide scans for ten phenotypes of the CFW mice dataset.** Six methods were used to perform GWAS, GEMMA, CMLM, FASTmrEMMA, FarmCPU, MLMM, and ISR. P-values quantify support for a QTL at each of the 92,734 candidate SNPs on autosomal chromosomes. Dotted red lines correspond to significance thresholds ($P < 0.1$, approaching $7.7 \times 10^{-6}$) estimated via permutation tests. Refer to Supplementary Table 5 for additional information.

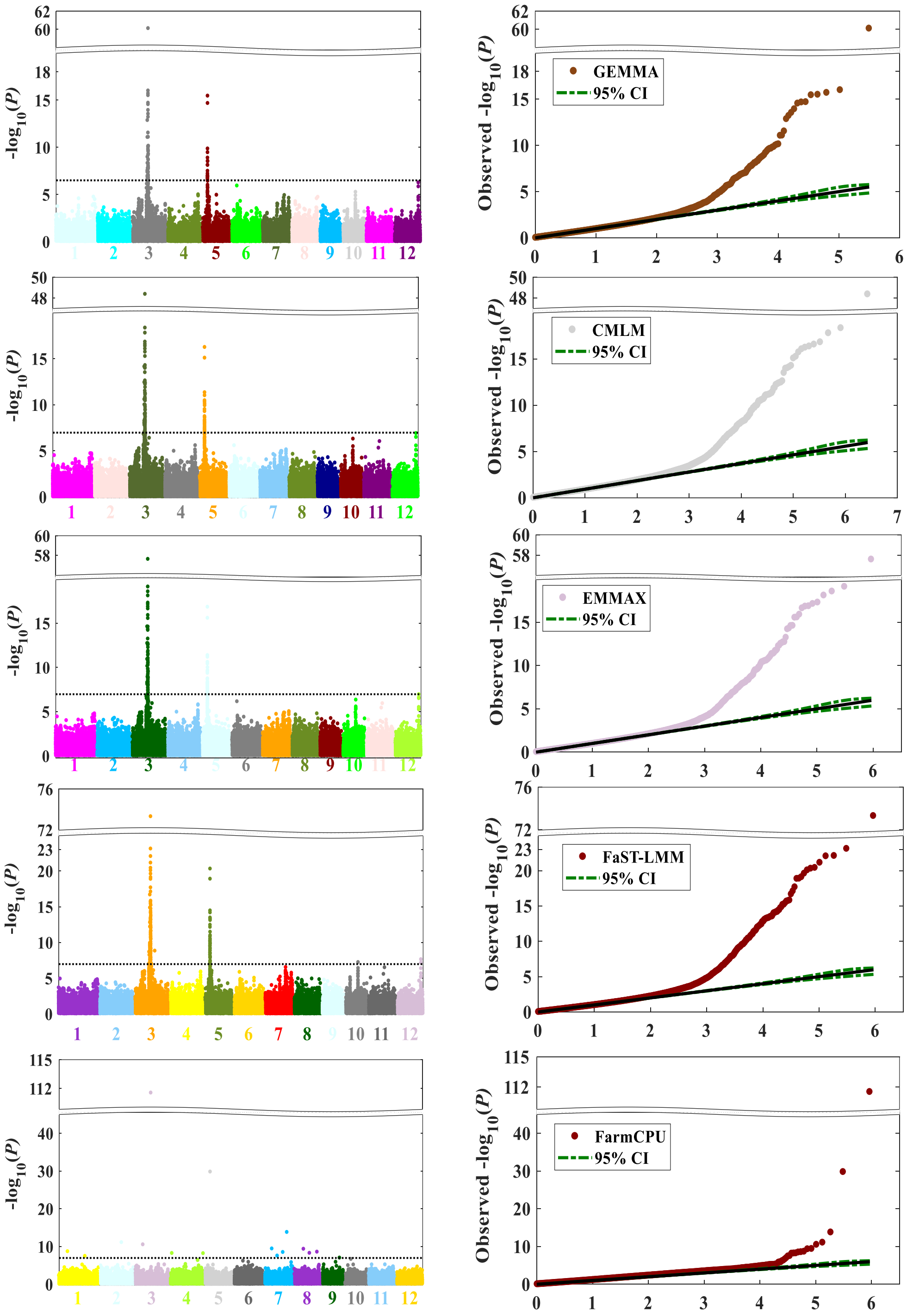

-log10(P)
Observed -log10(P)
GEMMA
95% CI
CMLM
95% CI
EMMAX
95% CI
FaST-LMM
95% CI
FarmCPU
95% CI
1 2 3 4 5 6 7 8 9 10 11 12

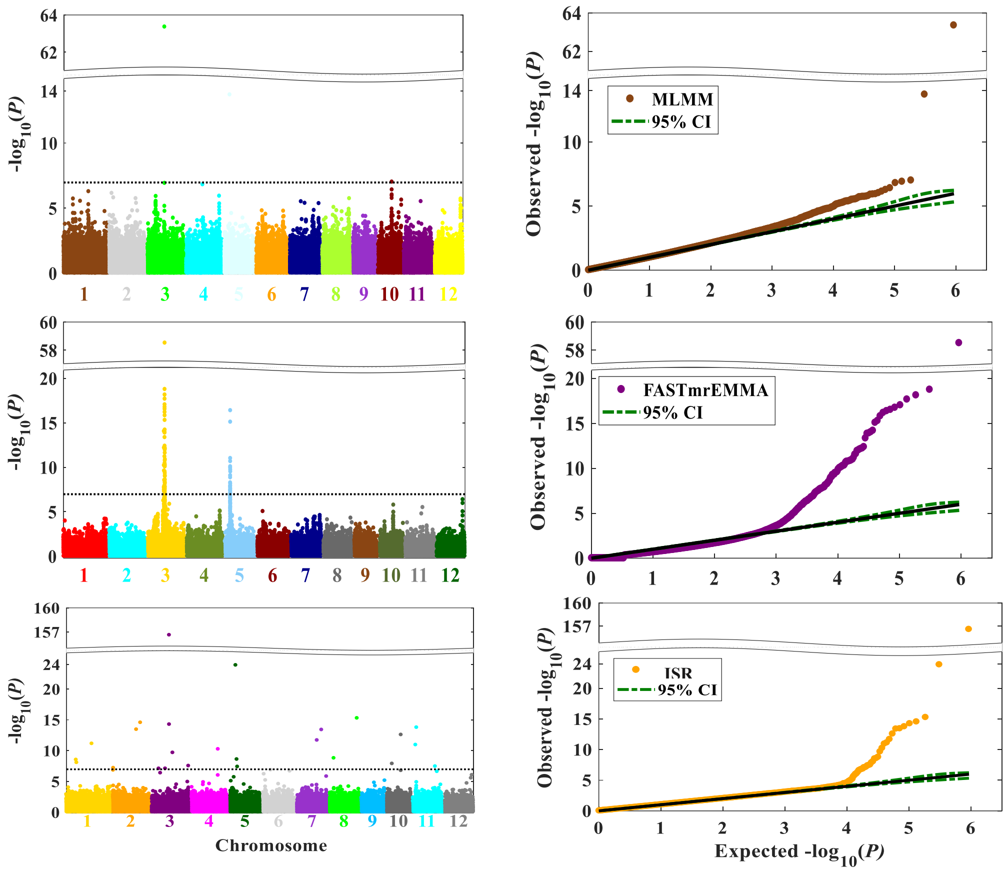


**Supplementary Figure 15. GWAS for rice grain length.** Manhattan plots (left panel) and quantile-quantile plots (right panel) for GWAS using eight different methods, GEMMA, CMLM, EMMAX, FaSTLMM, FarmCPU, MLMM, FASTmrEMMA, and ISR. Dotted black lines correspond to the significance thresholds level of 5% Bonferroni correction.

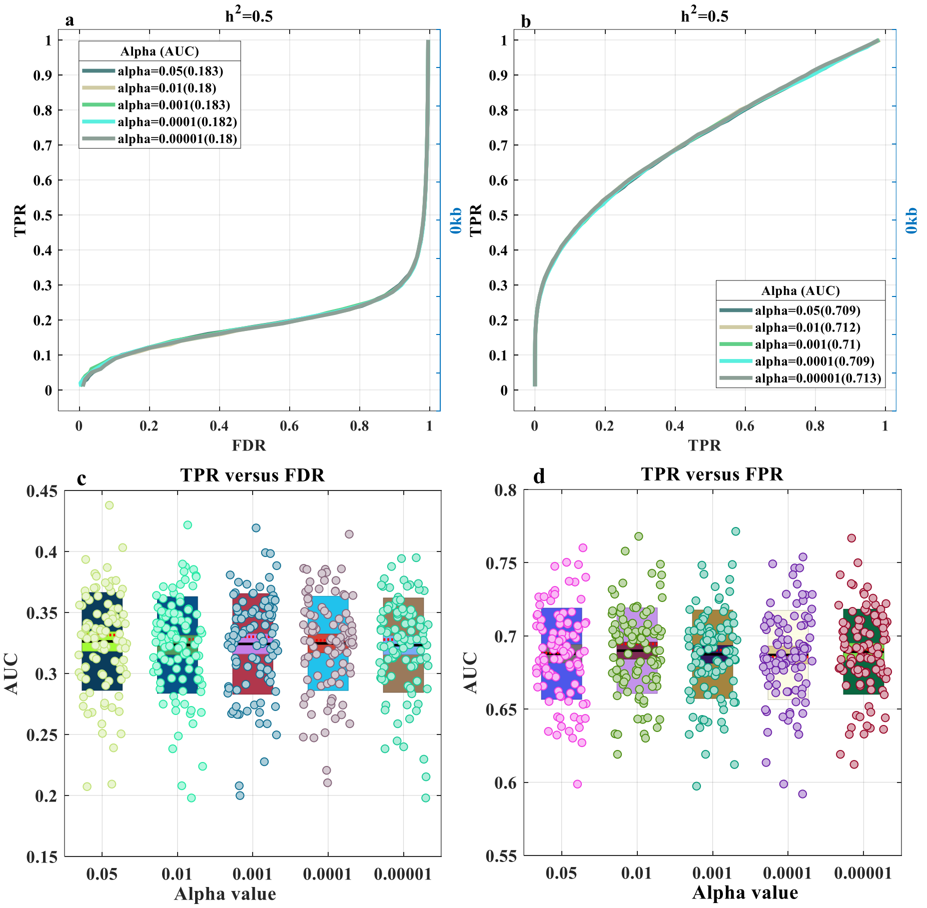


**Supplementary Figure 16. Performances of ISR with different alpha values in the second simulation scenarios.** Here, additive genetic effects are controlled by 100 causal loci with a phenotypic heritability of 0.5. We investigated the influence of dynamic significance level alpha value for model selection(0.05, 0.01, 0.001, 0.0001 and 0.0001) (Supplementary Note Fig.2). A causal SNP was considered to be detected if an SNP within 0kb (only casual SNP itself) on either side was determined to have a significant association, otherwise, it was considered a false positive. **(a,b)** The two types of Receiver Operating Characteristic (ROC) curves are displayed separately for TPR (power) versus FDR and FPR. **(c,d)** The Area Under the Curves (AUC) is also displayed separately for TPR (power) versus FDR and FPR for 100 simulations. The area measures the performance of detecting associations under the curve (AUC), where a higher value indicates better performance. It indicated that the alpha value merely slight (non-significant after ANOVA, p=0.6665, p=0.7394, respectively) impact the detected power.

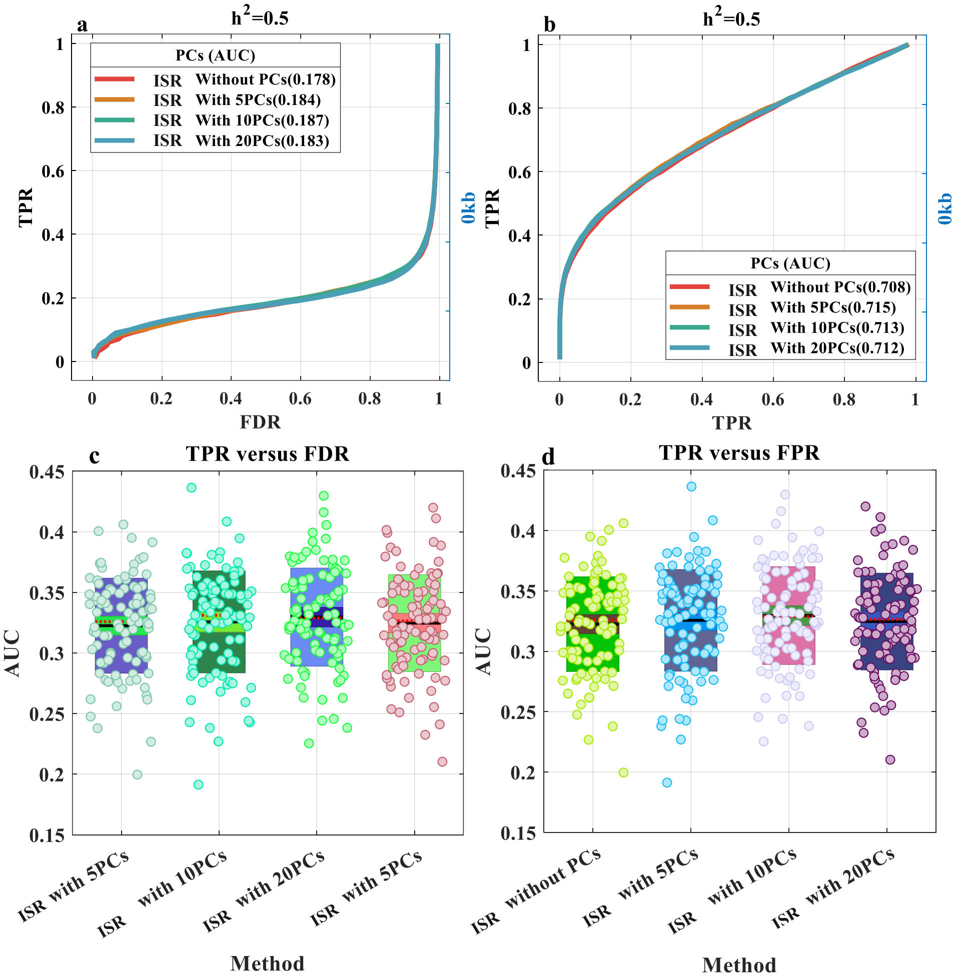


**Supplementary Figure 17. Effect of including population structure as covariates in ISR methods in the second simulation scenarios.** The simulated trait was controlled by 100 casual loci with a heritability of 0.5. Population structure was calculated as PCs, derived from all of SNPs. Different numbers of PCs (5, 10, and 20) were fitted in ISR. A causal SNP was considered to be detected if an SNP within 0kb (only casual SNP itself) on either side was determined to have a significant association; otherwise, it was considered a false positive. **(a,b)** The two types of Receiver Operating Characteristic (ROC) curves are displayed separately for TPR (power) versus FDR and FPR for 100 simulations. **(c,d)** The Area Under the Curves (AUC) is also displayed separately for TPR (power) versus FDR and FPR for 100 simulations. The area measures the performance of detecting associations under the curve (AUC), where a higher value indicates better performance. It also indicated that add PCs as covariates did not impact the detected power (p=0.6583, p=0.596, ANOVA, **c,d**).

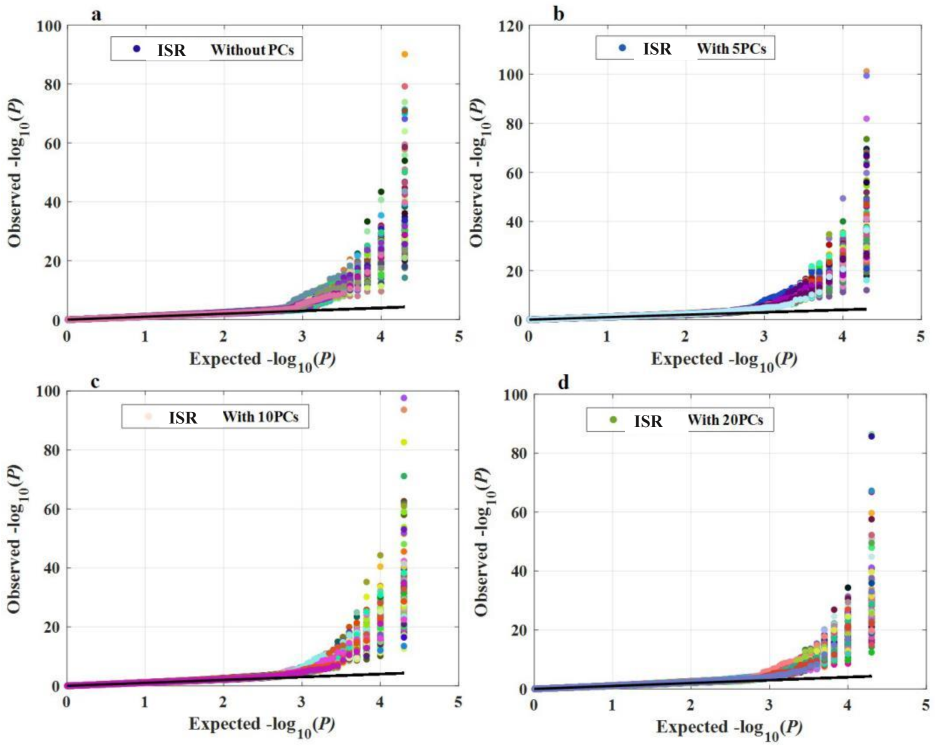


**Supplementary Figure 18. Comparing ISR with or without PCs for controlling inflation.** QQ plots of 100 randomly generated phenotypes under the second simulation scenarios. (**a)** ISR without PCs; (**b)** ISR with 5PCs; (**c)** ISR with 10PCs; (**d)** ISR with 20PCs; In contrast, ISR without PCs results in P values close to the expected distribution as same as others adjustment.

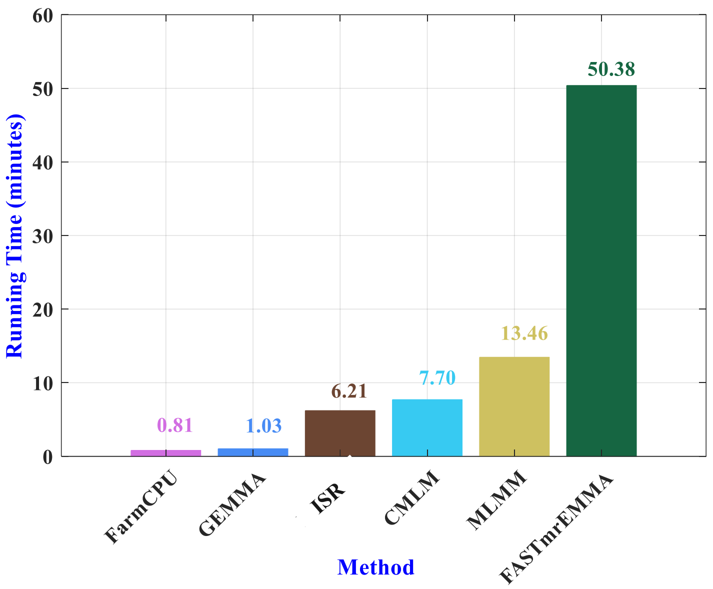


**Supplementary Figure 19. Comparison of computing time of ISR and other methods in the third simulation scenarios.** The average of computing times using five methods (FarmCPU-R, GEMMA-C++, CMLM-R, MLMM-R, and FASTmrEMMA-R) are compared with ISR-M (MATLAB language) for 100 simulations. The dataset containing 1161 individuals genotyped with 20000 markers. Simulation studies using a computer with an Inter(R) i3-6100 @3.7GHz CPU in windows 10.

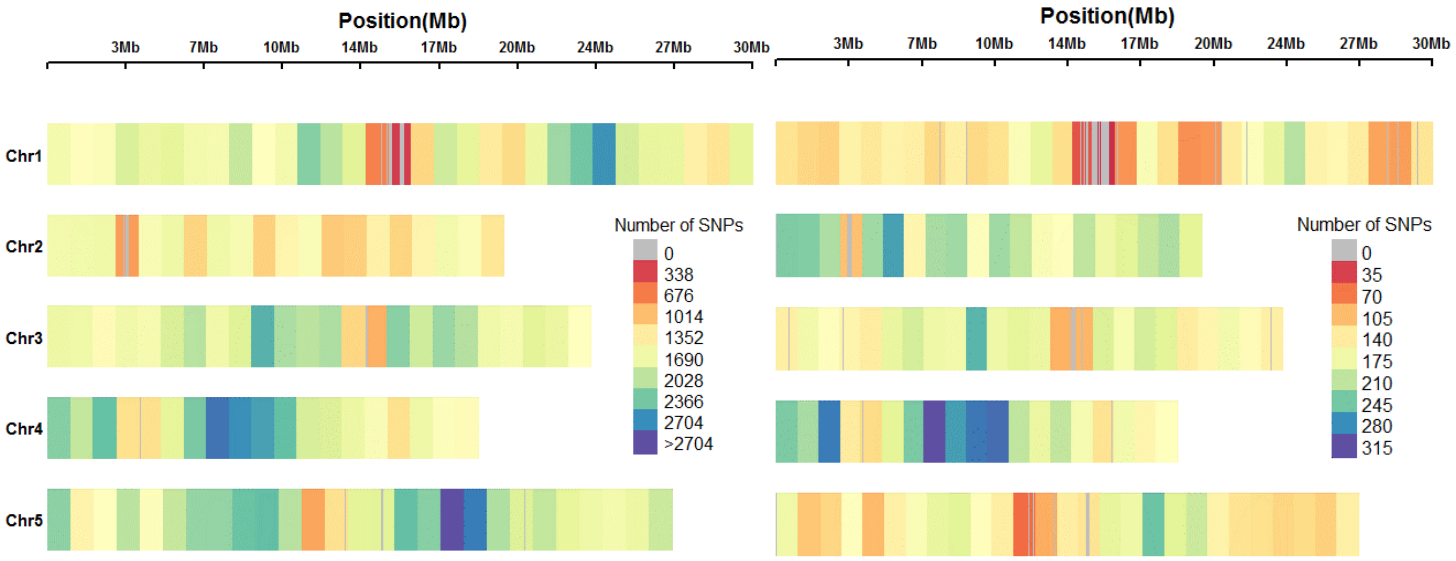


**Supplementary Figure 20. The density of SNPs discovered in 1,307 diverse Arabidopsis accessions.** The colored bars represent the number of SNPs, and each bar represents a 1Mb window size. The left figure that represents the density of the genome-wide 214,051 single-nucleotide polymorphism marker discovered, where the right figure that represents the randomly chose density of 20,000 SNPs to simulation discovered.

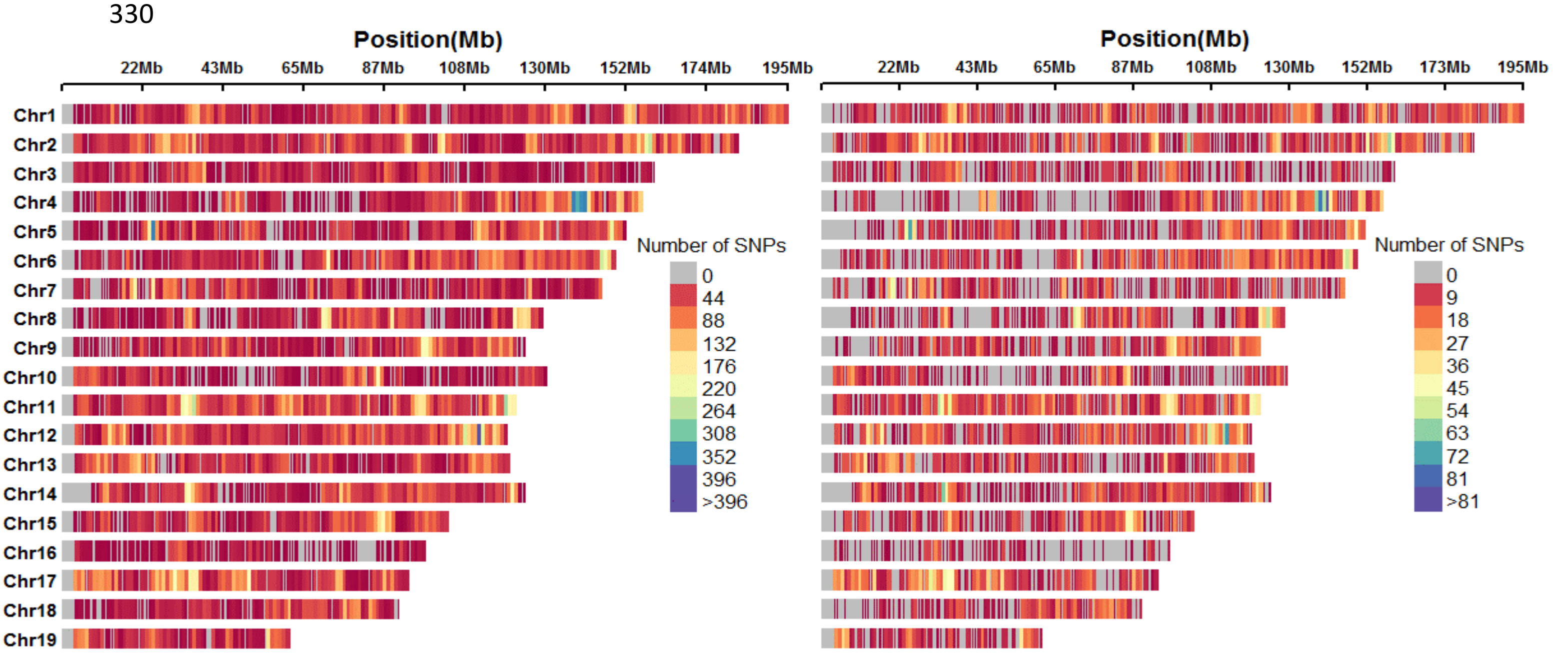


**Supplementary Figure 21. The density of SNPs was discovered in the CFW 1161 population.** The colored bars represent the number of SNPs, and each bar represents a 1Mb window size. The colored bars represent the number of SNPs. Each bar represents a 1Mb Window size. The left figure that represents the density of the genome-wide 92,734 single-nucleotide polymorphism marker discovered, where the right figure that represents the randomly chose density of 20,000 SNPs to simulation discovered.

1. Horton, M.W. *et al.* Genome-wide patterns of genetic variation in worldwide Arabidopsis thaliana accessions from the RegMap panel. *Nat Genet* **44**, 212-216 (2012).
2. Tucker, G., Price, A.L. & Berger, B. Improving the Power of GWAS and Avoiding Confounding from Population Stratification with PC-Select. *Genetics* **197**, 1045 (2014).
