## Supplementary Notes for "A new approach of dissecting genetic effects for complex traits"

### Supplementary Note: Model and Algorithm Details for ISR

#### The latent screen stepwise regression model

We consider the following multiple linear regression model:

$$y = W\alpha + X\beta + \varepsilon, \varepsilon \sim \mathrm{MVN}(0, \delta_e^2 \mathrm{I}_n) \quad (1)$$

where $y$ is an $n$-vector of phenotypes measured on $n$ individuals; $W=(w_1,w_2\ldots w_c)$ is an $n$ by $c$ matrix of covariates(fixed effects) including a column of ones for the intercept term; $\alpha$ is a $c$-vector of coefficients; X is an $n$ by $p$ matrix of genotypes; β is the corresponding $p$-vector of effect sizes; ε is an $n$-vector of residual errors where each element is assumed to be independently and identically distributed from a normal distribution with a variance $\delta_e^2$; $I_n$ is an $n$ by $n$ identity matrix and MVN denotes multivariate normal distribution.

The goal of GWAS is to detect loci associated with variation (SNPs) in traits of interest. Finding which of 100,000—1,000,000 or more loci has a practically significant effect is a challenging statistical problem, like finding a needle in a haystack[1]. Such that large $p$ small $n$ problems are challenging ($p$ is the number of SNPs, and $n$ is the number of individuals): there are may a strong correlation between the explanatory variables (QTNs) (in statistics called Multicollinearity). So the existence of linear or almost linear relations among covariates causes a big imprecision in the estimation of coefficients of regression. If the relations are rigorously linear, the coefficients would be biased. Herein, we proposed a new iterative screen stepwise regression model, where the ideal was from the stepwise regression[2], but the standard approach did not work well for large datasets, especially for GWAS. Caused being limited in exploring the model space, we proposed a new strategy that made the advantage of being computationally efficient and applicable to GWAS in a large dataset. The following procedures accurately describe the new model. Notably, our regression model is not limited to associations between phenotypes and genetic markers. Therefore, as following, we refer to them as response variable (phenotype) and covariates (markers).

#### Build the screening criterion of regression for model selection

In some way, AIC (Akaike information criterion)[3], BIC (Bayesian information criterion)[4] and EBIC[5] did not work well in model selection[6,7]. Here, we formulated a regression information criterion (RIC) and this criterion as the objective function of the entire computational procedures. The second built a unique and appropriate independent variable (single locus, SNP) screening technology, quietly said that selected from all the covariates have significant effects, simultaneously eliminating the effects of weak arguments. Still, it is different from general stepwise regression, including forward and backward[6,8]. So, we proposed a novel optimal algorithm ideal called expansion and contract variable screening procedure. Consequently, according to the continual changing regression information criterion to iterative screening variable, and eventually achieved optimal subsets (the subset of markers (SNPs) that together explains the phenotype best in GWAS[6,9,10]).

We consider the objective function of the following:

$$RIC = f\left(sd\right) \times F\left(p\right) \quad (2)$$

Where $F(p)$ is $F$ test value that when model selected $p$ covariates and defined $f$(sd) is approximate exponent function of saturating information function that is responded to the model saturation degree $sd\left(sd = \left(p+1\right)/\ n\right)$ (Fig.1). Then, we obtained the $f$(sd), which was a reverse $J$ graphical function inspired by $F_\alpha$ test critical value after reduplicative simulation testing (Fig.1). Apparently, the $f$(sd) value will tend to zero, when the $sd$ is largest (means model tends to saturate). Avoiding the phenomenon happened that infinity selected optimal subset as a cofactor. Diametrically, there is no chosen subset when $f$(sd) reaching its maximum. When the saturation degree of the model tends to zero, the $f$(sd)

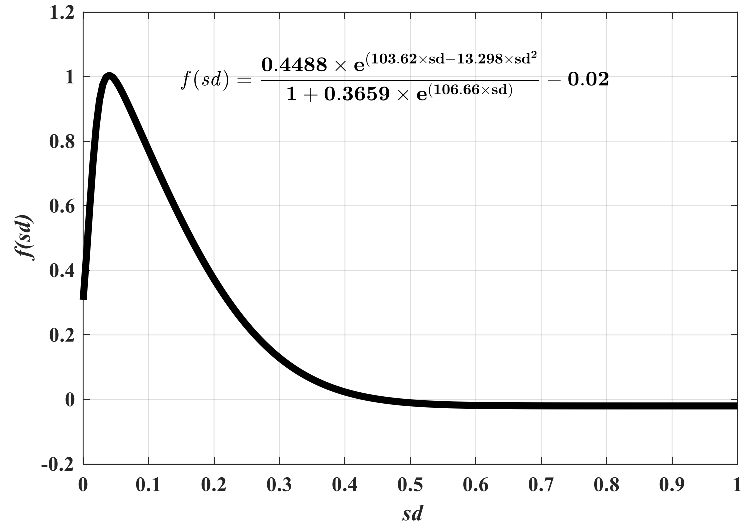


Figure 1. Equation of saturated information functions $f(\text{sd})$

the function appears a modest recovery (appear a peak) (Fig.1), such that when the number of covariates is already reached to appropriate small numbers, and the objective function will be stabilized at a peak. If the covariates have a weak effect on response (phenotypic variation, defined as following), they will not be removed. Moreover, only when all covariates are no effect that means effect equal zero, they also can be removed entirely. which can eliminate the phenomenon that happened that all covariates will be excluded. It can be seen that using the objective function as a model screening criterion is appropriate. This procedure will enable us to achieve the objective of the selected effect of SNPs smaller.

**Significant levels of dynamic regulation**

One of the main issues with stepwise regression is that it searches a large space of possible models. Thus an enormous search space can lead to overfitting and high variance of the coefficient estimates. Hence, after the objective function is determined, looking for the maximum objective function corresponding to the regression equation does not have to be related to a significant level. However, in finding the optimal objective function, selected covariates and removed covariates should be based on the probabilities screening. Inspired by the Bonferroni test, we combined significant level with the number of effects $p_c$ (or $sd$), and built a dynamic significance level functions $af(\alpha, \text{sd})$, here,

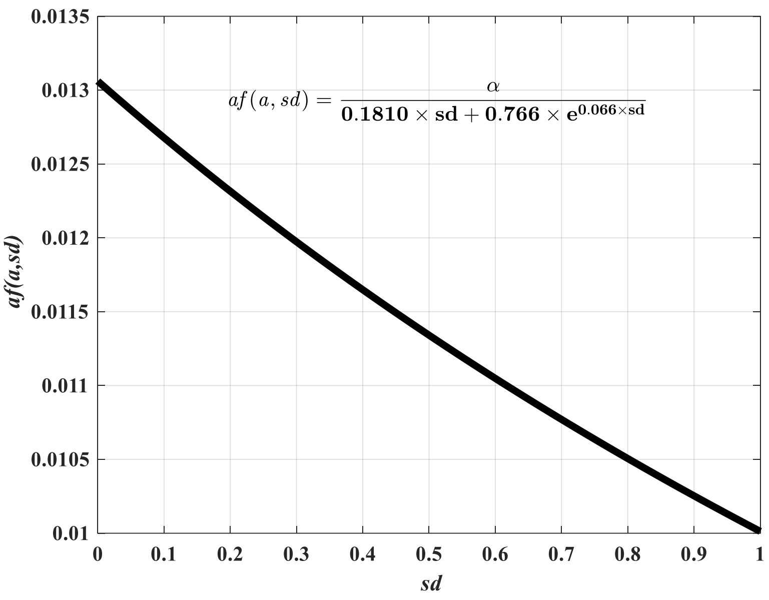


Figure 2. Dynamic significance level functions $af\,(0.01, \mathrm{sd})$

α=0.01(Fig.2), or can be 0.05, 0.001, 0.0001 (seeing the discussion section for arguing). When in a certain equation of saturates degree $sd$ , the covariates are removed and selected dynamics of significant level function (Fig.2). However, when the regression model had selected numerous covariates, that saturation degree of the model is high, which increases the covariate further selected threshold is the best choice to achieve better results.

**Calculate the number of effects $p_c$ and append effect terms $p_d$**

$p_c$ Is the total number of effects in the regression model, if an effect is a linear main effect, then the count is 1. Simulation research found that if the model unsaturated, but the determination of coefficient close to 1 (this is another form of saturation, we called interpretation saturation degree), which the deviation from the regression model is smaller, and existing some very weak effects will become a significant effect term and incorrect included the regression model. Here, we combined interpretation saturation degree (determination of coefficient $R^2$ ) with the number of effects and append effect terms $p_d$ , and built the $p_c$ of append function $p_d$=$fr2(\mathrm{R}^2)$ (Fig.3), well, the proceeding $p_c$ was as above describe and $fr2(\mathrm{R}^2)$ value. Which can regulate and refrain from continuing to increase covariates selections of a vicious circle, when the determination coefficient close to 1.

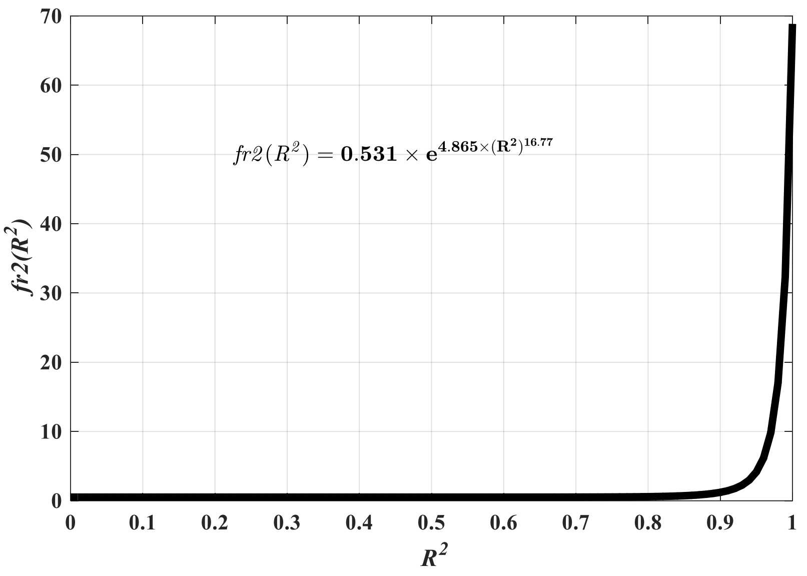


Figure 3. Append effects function $p_d = fr2(\mathrm{R}^2)$

#### The relationship between the objective function and optimal subset

In practical simulation analysis, we found that the objective function and the optimized subset is not one-to-one relations. That is to say, existing an objective function is similar or even equal while providing the optimized subsets was significantly different situations. In such cases, we adhere to the only standard of principle that taken the objective function, optimizing the objective function, and make it more rational and universal. By doing this, the correlation between the objective function and optimal subsets will be stronger and stronger.

#### The procedure of iterative screen stepwise regression

After model selection criteria are established, screening optimal subsets from all covariates is divided into the following three stages.

##### *Basic regression stage*

We consider the following standard linear model:

$$y_j = \sum_{i=1}^{p} X_{ij}\beta_i + \varepsilon_j$$

Where the continuous response variable *y* is a vector of *n* phenotypic observation, $X_{ij} \in \{0,1,2,\ or, 0,0.5,1\}, i = 1,2,\dots,p, j = 1,2,\dots,\mathrm{n}$ is an *n* by *p* design matrix of covariates (markers, $p$ is the number of SNPs in GWAS), $\beta$ is a vector of $p$ effects, and $\varepsilon$ is a vector of $n$ residuals.

If the number of covariates ($m$) less than the number of observations ($n$) ($m<n$), then all the covariates only perform classical multivariate linear regression analysis ($p=m$). While if the number of covariates more than or equal the number of observations ($m \geq n$), choosing the covariates less than the number of observations performs classical regression analysis (p<n<m). Synchronously, obtained each covariate of the partial regression coefficient $\beta_i (i=1,2,...p, p+1, \beta_1 \sim \mathrm{int}\,ercept)$ , partial regression sum of squares $U_{p_i} = \frac{\beta_{i+1}^2}{C_{i+1,i+1}} (\mathrm{j}=1,2,...,\mathrm{p})$ , the sum of squares due to deviation from the regression $Q = Y'Y - \beta' K$, the deviation regression variances $MS_Q = \frac{Q}{n-p-1}$ .

***Remove and select procedures***

After the previous step, removed from the selected covariates with partial regression sum of squares minimum and no significant effects (if selected effect terms are significant, won't have to remove any covariates), then continuing to select previous not be selected. Do the same last step after obtaining the new model (primary regression analysis). Keep repeating this process until all the covariates have been trying selected. The specific details described by the following:

( I ) Calculated the initially selected covariates of partial regression sum of squares and find the covariate $X_l \left(l=1,\ 2,\ \ldots, p\right)$ with minimum partial regression sum of squares :

$$U_{p_l} = \min_{1 \leq i \leq p}(U_{p_l}), (i=1,2,...,\mathrm{p}) \qquad (1)$$

(II) F-test:

$$F_l = \frac{U_{P_l}/1}{MS_Q} = \frac{B_{l+1}^2 / C_{l+1,l+1}}{Q/(n-p-1)} \qquad (2)$$

If $F_l < F_\alpha$, then, removed covariate $X_l$ from the new model ( $p = p-1$ ). If $F_l \geq F_\alpha$, keep the covariate $X_l$, and turn into the next step.

(III) Selected covariate in sequence and simultaneously change the data structure matrix, which has been selected from the total number of variables $p = p+1$, calculated the new model of regression statistics, and other summaries statistics. However, if the new model leads to dissatisfaction with the rank of $X$, it needs to be removed, and other data structure correspondingly changes. Similarly, repeat the above three steps until all the covariates have been trying selected. Moreover, these procedures require attention to the following two issues:

1. Setting the significance level is slightly larger, so avoiding the real effect would be not removed in the first step.
2. Recording the removed covariates, which will be used for the next rescreen step.

***Iterative screen procedure***

In ultra-high dimensional feature space[11], such as GWAS, closely linked markers (covariates) are highly correlated due to the lack of sufficient recombination between them[9,12]. Except for correlations based on physical linkage, population structure introduces correlations between markers even if they are weakly linked or not linked at all. These correlations can lead to spurious associations between markers (covariates) and phenotypes (response variable) and thus need to be corrected in any screen (testing) procedure. Moreover, in the above steps, the setting of significant levels is too big; it is also easier to improperly select some small effects (weak effects). Therefore, result in a subset of covariates is far from to obtain optimal subsets. Here, we proposed a novel two-steps to correction reselected procedures (Fig.4). Details describe as following:

(i) expansion screen step

Using remove and select procedures, which one of its chosen covariate and own partial regression sum of squares smallest was removed, and reselecting previously unselected all covariates. Usually, setting a relative bigger significant levels for this step (generally, establishment function relation between $\alpha$ and $P_c$, $\alpha' = \alpha / P_c$, the size of $\alpha$ is influenced by $P_c$ adjustment, where the $\alpha$ was from Fig.2), leading to rediscover the previous covariates could not be selected and have large effect (strong effect) to the response variable. By this step, numerous covariates will be reelected included in the models and were considerably increased, meantime, expanded the regression function. Worthy of noting that after at the end of each round, recorded the objective function value (RIC) of in the new model. By comparing each recoded model and chose the best model to obtain optimal subsets.

(ii) contraction select step

After the expansion screen step, the number of covariates was multiply increased. At this point, we used a smaller significant level (Fig.2) to reselect these covariates. Removed the small effect of covariates from the models to the contract regression function. To make the number of optimal of covariates that contains at a reasonable amount in the model. At the end of each round, recorded the new model of the value of the objective function (RIC).

Taken these two steps (Fig.4) as iterative screen procedure of one round. However, obtaining the optimal subset of covariates not only running this process one time, but it needs to keep repeating the process until the objective function value is stable after six to eight-round, and basically can be considered as an optimal subset of covariates, and to end this process.

The iterative screen procedure makes it more possible that obtained a surer optimal subset of covariates. However, the vast data (especially in GWAS, could be millions dataset) need to do more rounds of calculations were very time-consuming (computation burden). However, in linear regression, the magnitude of regression coefficients (i.e. the effect size of a covariate) also provides useful information for variable selection. Therefore, in

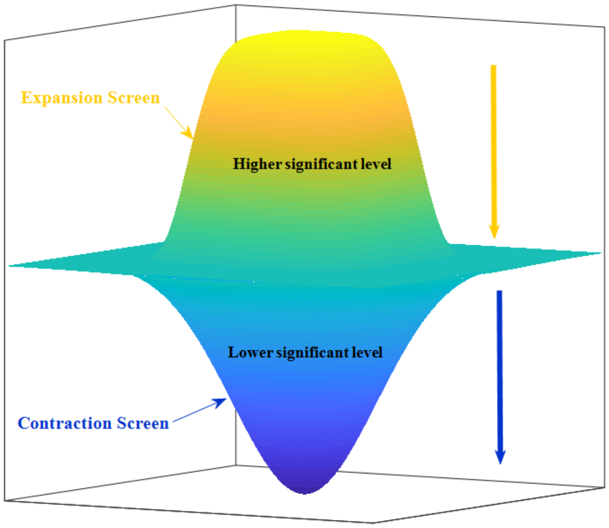


Figure 4. The diagram of re-screen steps

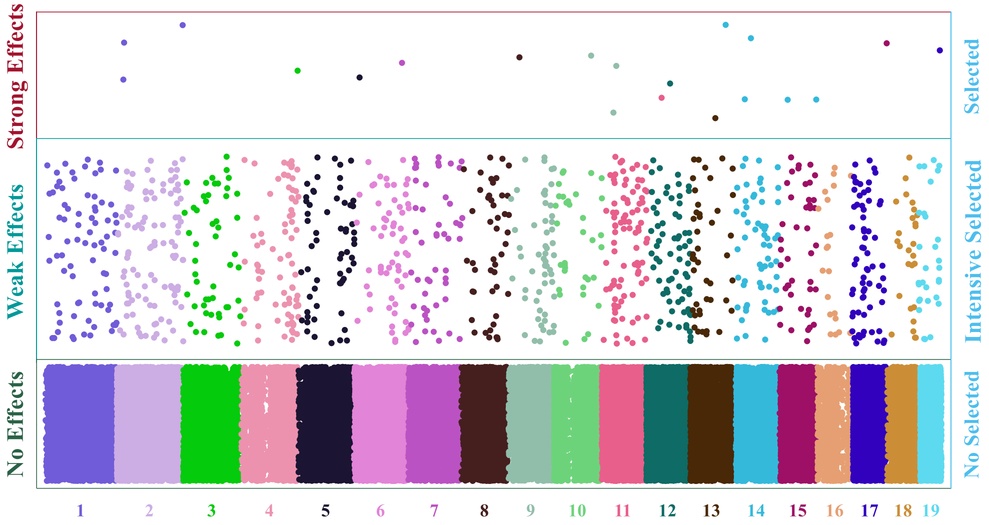


Figure 5. The diagram of three classified

response to this problem, which we have adopted the following strategies to specific define the covariates of unselected. In the searching process, according to the size of the F-test value for each effect term (here, mainly was additive effects), and all the covariates are divided into three categories (strong effects, week effects, and no effect term) (Fig.5). The first part of the effect terms are extremely significant of covariates to response variable and call strong effect term (effect terms appeared or ever appeared in optimization model), usually, these covariates can be selected in the model quickly. The second part of the effect terms are significant (or $F_{test}$>=1) of covariates to response variable and call week effect term (effect terms appeared or ever appeared in suboptimization model). However, it is a bit of a difference from the first part that is selected in the model or not. The third part of the effect terms are no significant ($F_{test}$<1) of covariates to response variable and call no effect term; generally, they cannot be selected in the model.

Based on the above reasons, those covariates that are not selected that according to its probability of partial regression tests (F-test) divided into weak effect and no effect two parts (Fig.5), and intensive selected the weak effect of covariates. However, the weak effects of selection should have an appropriate amount if the numbers are too small, some effects could not be selected, and unable to get the optimal subset of covariates (reasonable or representative covariates). On the contrary, it will cause a giant increase in the computation burden and low efficiency. On the other hand, it will also affect the optimal subset of covariates obtained. Through simulation analysis and summary, generally considered the weak effect of re-selected of the number to a total of about 5%-10% (the number of points in Fig.5). Moreover, this section of intensive selected should according to the number of weak effects and no effects of covariates feedback regulation.

As above stated, between the covariates, they may are exist correlated. In this condition, screening between covariates will affect the return of the other covariates associated with regression significant, and previously identified weak effects or no effects of covariates may be biased. Therefore, after a certain round of repeated screen obtained a new model, which needs to reauthenticate the weak effects and no effect and reclassified.

In the global optimization process, removed or selected some covariates in the model still cannot make the objective function feature optimization that production local optimum (no matter how many rounds/iterations that still were unable to acquire the optimal subset of circumstance). Included our new method iterative screen stepwise regression model. In order to solve this problem, in the process of obtaining optimal subsets, having taken two different expansion and contraction optimization technology into consideration. The first is expansion screening and contraction screening procedure (Fig.4). The second is excluding the effects of covariates not always with partial regression sum of squares minimum, sometimes, choosing the covariates that have medium size of partial regression sum of squares $U_{p_i} (\mathrm{j}=1,2,...,\mathrm{p})$. By adopting these strategies can jump out from the local optimum to get an optimal subset of covariates.

***The expansion ratio and contraction ratio of feedback regulation***

Expansion ratio (ER) means the initial expansion screen step and the final step of the ratio of the number of covariates contained in the model. Similarly, the contraction ratio (CR) means the initial contraction screen step and the final step of the ratio of the number of covariates contained in the model. We found that the size expansion ratio and contraction ratio greatly influenced the ability to achieve an optimal model. Our methods regulation the expansion ratio and compression ratio, and imposed dynamic change (ER and CR) in the searching optimal model process. Which efficiently improved the ability to search the latent variables to achieve the optimal model.

***The storage capacity of weak effect term of feedback regulation***

Weak effect terms are the foremost part of strengthening screening, while the storage capacity of weak effect terms (the number of covariates that have the weak effect) is an influence on optimizing of ability and efficiency (search the best subset). One of the key points is maintained the storage capacity of weak effect terms to achieve global optimal objective function. When the storage capacity of weak effect terms is too small, an optimal regression model is not easy to obtain, oppositely will make searching optimize of ability and efficiency decline. Herein, we are using the critical value $c$ (the difference between optimization and sub-optimization model of the objective function is the critical value $c$ ). The storage capacity of a weak effect term by the critical value $c$ feedback regulation. However, it is difficult to determine appropriate critical value $c$. It needs to phase adjustment during the optimization process. Which was certain rounds in the searching optimization process, checking the size of the storage capacity of weak effect term, when storage capacity is too small, increasing the critical value $c$, on the contrary, reduce critical value $c$, and removed all the weak effect of covariates.


1. Gondro, C., Werf, J.V.D. & Hayes, B. *Genome-Wide Association Studies and Genomic Prediction*, 1-17 (Humana Press, 2013).
2. Hocking, R.R. A Biometrics Invited Paper. The Analysis and Selection of Variables in Linear Regression. *Biometrics* **32**, 1-49 (1976).
3. Akaike, H. Information Theory and an Extension of the Maximum Likelihood Principle. in *Selected Papers of Hirotugu Akaike* (eds. Parzen, E., Tanabe, K. & Kitagawa, G.) 199-213 (Springer New York, New York, NY, 1998).
4. Schwarz, G. Estimating the Dimension of a Model. *Ann. Statist.* **6**, 461-464 (1978).
5. Chen, J. & Chen, Z. Extended Bayesian information criteria for model selection with large model spaces. *Biometrika* **95**, 759-771 (2008).
6. Segura, V. *et al.* An efficient multi-locus mixed-model approach for genome-wide association studies in structured populations. *Nat Genet* **44**(2012).
7. Huang, M., Liu, X., Zhou, Y., Summers, R.M. & Zhang, Z. BLINK: A Package for Next Level of Genome Wide Association Studies with Both Individuals and Markers in Millions. *bioRxiv* (2017).
8. Knüppel, S. *et al.* Multi-locus stepwise regression: a haplotype-based algorithm for finding genetic associations applied to atopic dermatitis. *BMC Medical Genetics* **13**, 8 (2012).
9. Klasen, J.R. *et al.* A multi-marker association method for genome-wide association studies without the need for population structure correction. **7**, 13299 (2016).

10. Rakitsch, B., Lippert, C., Stegle, O. & Borgwardt, K. A Lasso multi-marker mixed model for association mapping with population structure correction. *Bioinformatics* **29**, 206-214 (2013).
11. Fan, J. & Lv, J. A Selective Overview of Variable Selection in High Dimensional Feature Space. *Statistica Sinica* **20**, 101-148 (2009).
12. Wasserman, L. & Roeder, K. High-dimensional variable selection. *Ann. Statist.* **37**, 2178-2201 (2009).
